# Plant breeding simulations with AlphaSimR

**DOI:** 10.1101/2023.12.30.573724

**Authors:** Jon Bančič, Philip Greenspoon, Chris R. Gaynor, Gregor Gorjanc

**Affiliations:** The Roslin Institute and Royal (Dick) School of Veterinary Studies, University of Edinburgh, Easter Bush, Edinburgh, EH25 9RG, Midlothian, United Kingdom.; Bayer Crop Science, 700 Chesterfield Pkwy W., Chesterfield, MO 63017, Missouri, USA.

**Keywords:** AlphaSimR, plant breeding, stochastic simulation, self-pollinated crop breeding, hybrid breeding, clonal breeding

## Abstract

Plant breeding plays a crucial role in the development of high-performing crop varieties that meet the demands of society. Emerging breeding techniques offer the potential to improve the precision and efficiency of plant breeding programs; however, their optimal implementation requires refinement of existing breeding programs or the design of new ones. Stochastic simulations are a cost-effective solution for testing and optimizing new breeding strategies. The aim of this paper is to provide an introduction to stochastic simulation with software AlphaSimR for plant breeding students, researchers, and experienced breeders. We present an overview of how to use the software and provide an introductory AlphaSimR vignette as well as complete AlphaSimR scripts of breeding programs for self-pollinated, clonal, and cross-pollinated plants, including relevant breeding techniques, such as backcrossing, speed breeding, genomic selection, index selection, and others. Our objective is to provide a foundation for understanding and utilizing simulation software, enabling readers to adapt the provided scripts for their own use or even develop completely new plant breeding programs. By incorporating simulation software into plant breeding education and practice, the next generation of plant breeders will have a valuable tool in their quest to provide sustainable and nutritious food sources for a growing population.

## 1 Introduction

Stochastic simulations are a cost-effective tool for the development, validation, and optimization of new breeding methods to enable the continuous refinement of real breeding programs. They can also serve as a powerful educational platform to learn the principles of plant breeding and quantitative genetics. In this contribution, we present the process of building a plant breeding program simulation and provide complete AlphaSimR scripts of different breeding programs and techniques, suitable for training, research, or even practical implementation.

Plant breeding has gone through a significant transformation since its inception. The first form of selective breeding by humans started about 10,000 years ago with selection on traits such as non-shattering, seed/fruit appearance, and plant stature. This plant domestication process lasted for millennia, transmitting knowledge and improved plant materials through generations, and developed a set of plant species that are still widely cultivated today to feed our societies. With an improved understanding of the principles of inheritance and genetics, as well as improved management practices, plant breeding has increased in sophistication, precision, and scale, particularly in the last century. For example, maize and wheat productivity has nearly tripled and rice and soybean crop yields have doubled between 1961 and 2021 [1]. As such, modern selective breeding has become effective in developing varieties that are precisely tailored to specific environments, production systems, and consumer preferences.

Despite significant achievements in modern plant breeding, annual genetic gains in most staple crops fall short of the recommended targets stipulated by the Food and Agriculture Organization of the United Nations (FAO). On average, annual gains in staple crops are estimated to be around 1%, while FAO guidelines stipulate a minimum of 2% to meet the ever-growing global food demand [2, 3]. Several factors contribute to the limited annual genetic gains in plant breeding. First, breeders may not be able to develop new stable varieties that keep up with increasingly changing environmental conditions. Second, due to the decreasing availability of arable land in some regions, breeders are under pressure to develop varieties that are not only higher yielding, but also more resource efficient, stress tolerant and nutritious [4]. Breeding for multiple traits simultaneously reduces yield due to trade-offs, requires maintenance of extensive genetic diversity, and sometimes involves lengthy cycles to introgress donor material into elite material [5]. Finally, long breeding cycles, lasting 7-12 years in many cereals and even more in perennials, make it difficult to deliver improved varieties in a timely manner [6]. To address these challenges, it is necessary to efficiently use the limited resources available to a breeding program.

Running breeding program experiments with simulation prior to implementing them in the field allows rapid evaluation of which promising avenues to be pursued and dead ends to be avoided.

A typical breeding cycle of a modern plant breeding program involves common steps regardless of the species considered. These steps include (Figure 1):

**Fig. 1.**
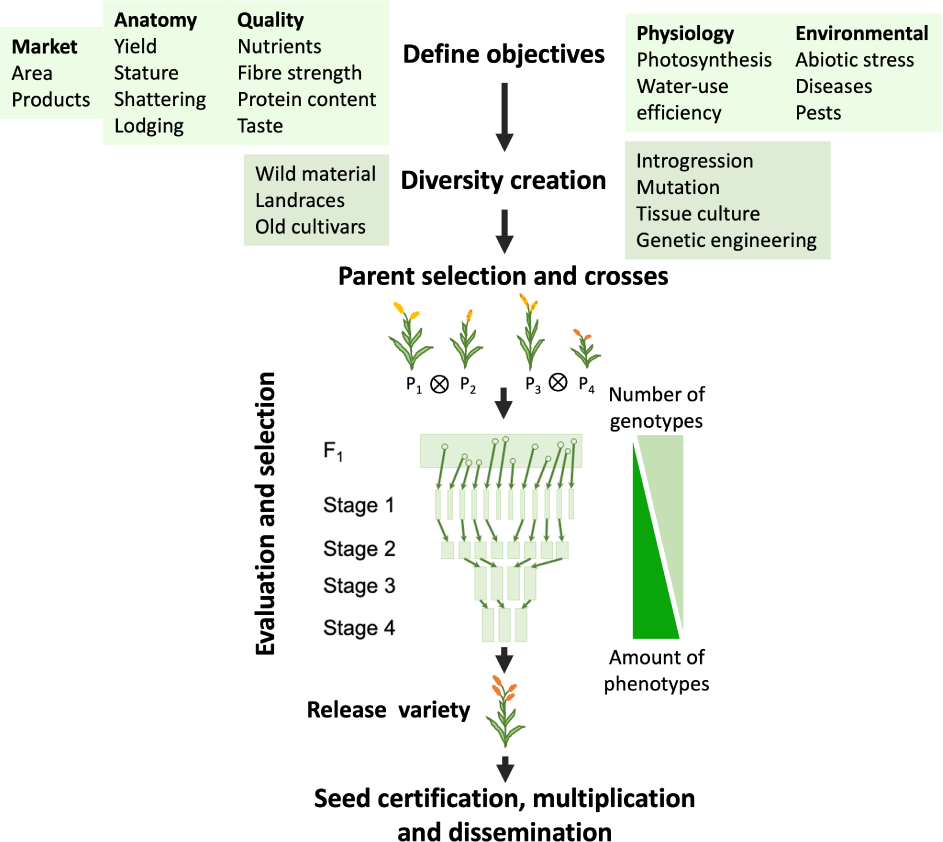
Common steps of a plant breeding program.

i) defining breeding objectives - identifying the traits to be improved in the breeding program and the targets to aim for; ii) creating diversity - evaluating a range of plant genetic resources to identify potential breeding material; iii) parent selection - choosing best genotypes with desired traits and using these genotypes as parents of the next generation; iv) crossing - creating progeny by crossing the parents to generate the genetic diversity in a candidate population needed for selection, and v) evaluation and selection - assessing candidate genotypes and identifying superior ones for a wide range of traits over multiple stages, locations, and years. The final step of a breeding pipeline involves seed multiplication and commercial release of a newly developed variety after it has successfully passed the registration and certification process. Apart from the obvious differences that can arise among breeding programs due to available resources, differences in programs also arise from factors such as the mode of reproduction (e.g., self-pollinated, cross-pollinated, or clonally propagated), the desired level of genetic uniformity and heterozygosity in the released variety, and the importance of dominance genetic variation driving both heterosis and inbreeding depression [7]. Due to the long process of variety development, breeding programs adopt new methods to efficiently generate and identify the best genotypes with available resources.

Recent years have seen a surge in the development and adoption of new methods that shorten the length of breeding cycle and improve the accuracy of selection to help accelerate genetic progress. These new methods include the use of pre-breeding, bridging, and introgression of exotic/landrace material [8–10], genome sequencing and molecular markers [11–13], speed breeding [14], genome editing and targeted recombination [15, 16], advanced statistical techniques for efficient field trial analysis and prediction (e.g., [17, 18]), and the use of remote sensing for high-throughput phenotyping, image analysis, and environmental data [19–21]. The application of such new methods in breeding programs is often constrained by uncertainties about the potential impact on genetic progress, logistics, and costs. Simulation can provide insights about these uncertainties by indicating potential benefits, which can guide the design of validation experiments and downstream practical implementation.

Stochastic simulations are a popular tool for the development, validation, and optimization of breeding methods and pipelines. Through abstract modelling of the real system under a large number of scenarios, stochastic simulations provide researchers with an approximation of what could happen in reality [22]. Simulations enable researchers and breeders to make informed decisions before spending time and resources on field experiments. This power of simulation has motivated the release of several simulation software that are available as standalone programs or as packages for the R, Python, or Julia environment. Some of the prominent examples are: AlphaSim [23, 24], AdamPlant [25], ChromaX [26], MoBPS [27], PyBrOpS [28], QU-GENE [29], and XSim [30]. Most of this software can accommodate features such as additive, dominance, and epistatic genetic effects; complex family structures; selection on multiple traits controlled by different numbers of quantitative trait loci (QTL); and to some extent, genotype by environment interaction. Numerous publications demonstrate the usefulness of simulations. For example, simulations have been used to test and compare different breeding strategies [31, 32], including designing novel breeding systems [33]; optimize the implementation of genomic selection and mating strategies in different crop species [11, 34]; find the best statistical models for prediction and analysis of experimental data [35, 36]; evaluate gentoyping and imputation strategies [37, 38]; and test different strategies that introduce and manage genetic variation [9, 12, 39]. As a learning resource, Table 1 summarizes simulation studies relevant to plant breeding that used AlphaSimR. With the growing use of simulation in breeding program development, it is important for students, researchers, and practitioners to gain proficiency in the use of simulations and to understand how simulation is used to shed light on real-world problems. Although there are now a number of studies employing stochastic simulations, perspective papers on how to approach such studies [40, 41], and online course material (https://www.edx.org/course/breeding-programme-modelling-with-alphasimr), there remains a lack of (and demand for) literature providing guidance on how to practically implement and deploy such plant breeding simulations for various crops.

**Table 1:**
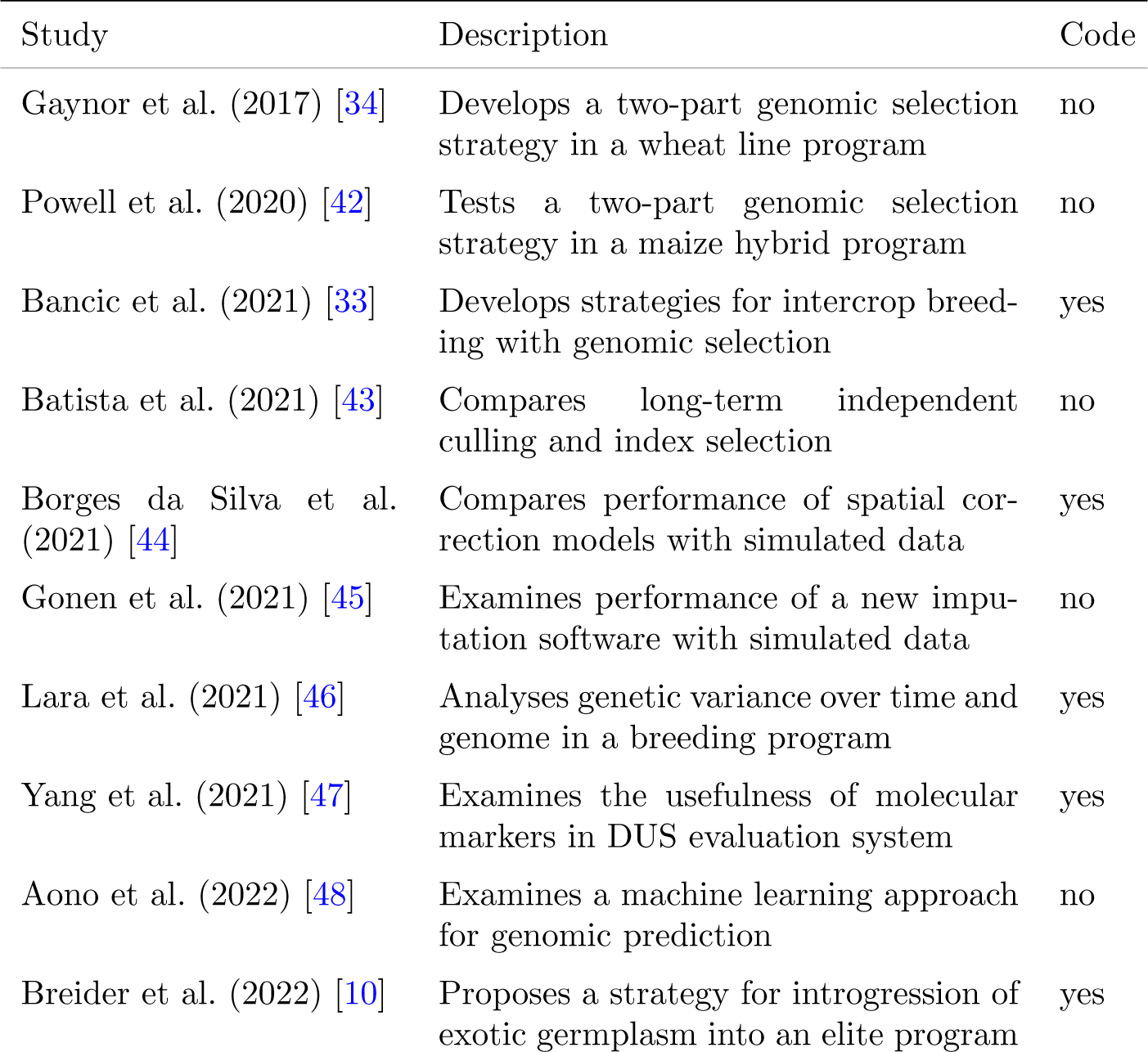
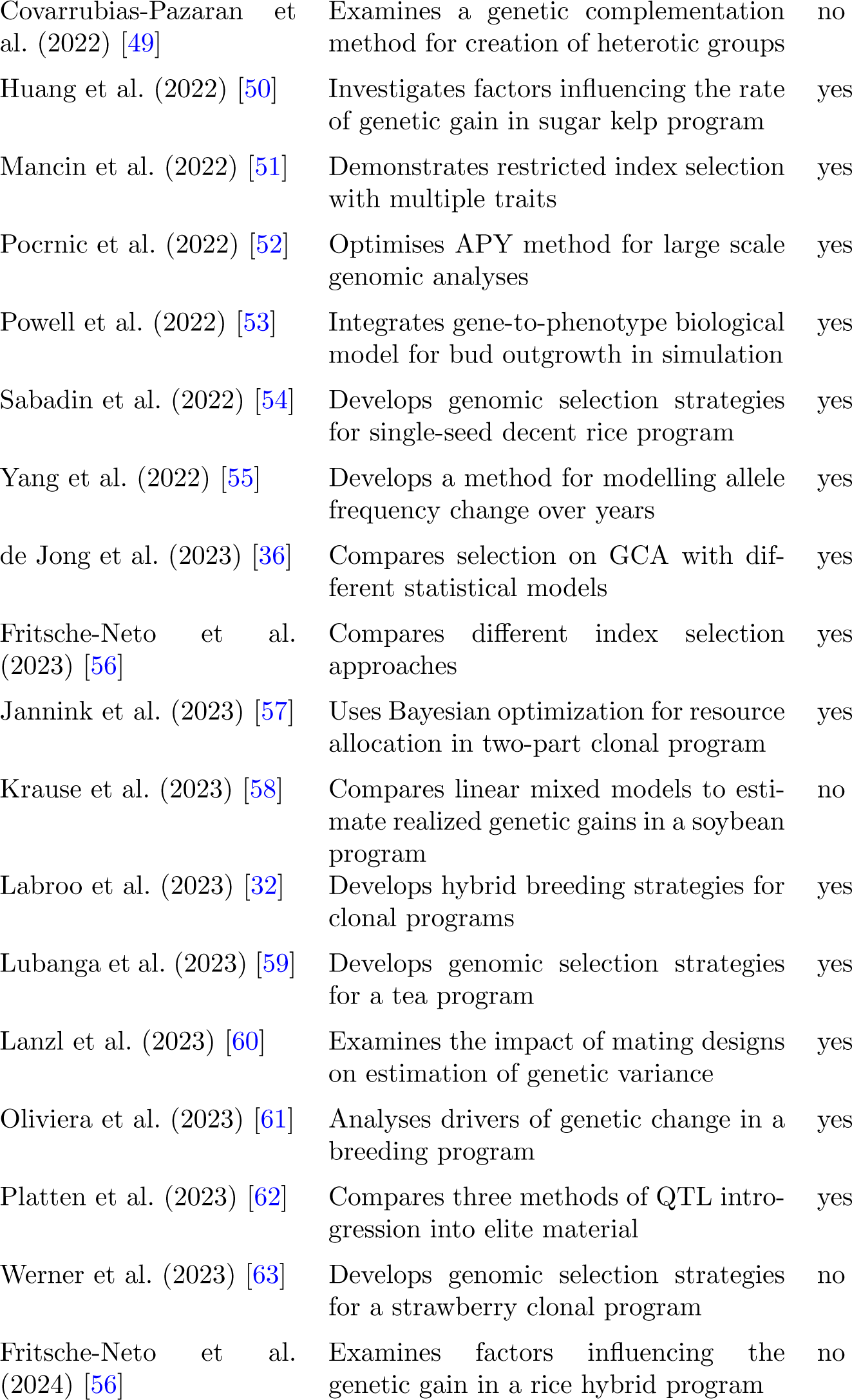
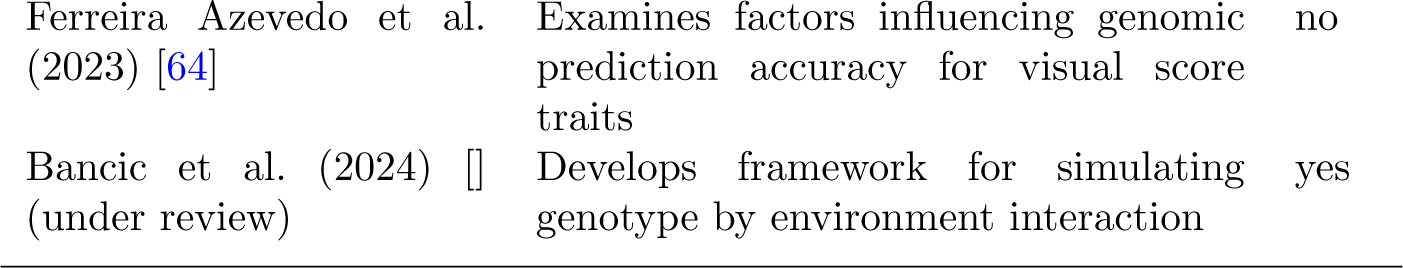
Simulation studies relevant to plant breeding using AlphaSimR.

| Study | Description | Code |
| --- | --- | --- |
| Gaynor et al. (2017) [34] | Develops a two-part genomic selection strategy in a wheat line program | no |
| Powell et al. (2020) [42] | Tests a two-part genomic selection strategy in a maize hybrid program | no |
| Bancic et al. (2021) [33] | Develops strategies for intercrop breeding with genomic selection | yes |
| Batista et al. (2021) [43] | Compares long-term independent culling and index selection | no |
| Borges da Silva et al. (2021) [44] | Compares performance of spatial correction models with simulated data | yes |
| Gonen et al. (2021) [45] | Examines performance of a new imputation software with simulated data | no |
| Lara et al. (2021) [46] | Analyses genetic variance over time and genome in a breeding program | yes |
| Yang et al. (2021) [47] | Examines the usefulness of molecular markers in DUS evaluation system | yes |
| Aono et al. (2022) [48] | Examines a machine learning approach for genomic prediction | no |
| Breider et al. (2022) [10] | Proposes a strategy for introgression of exotic germplasm into an elite program | yes |
| Covarrubias-Pazaran et al. (2022) [49] | Examines a genetic complementation method for creation of heterotic groups | no |
| Huang et al. (2022) [50] | Investigates factors influencing the rate of genetic gain in sugar kelp program | yes |
| Mancin et al. (2022) [51] | Demonstrates restricted index selection with multiple traits | yes |
| Pocrnic et al. (2022) [52] | Optimises APY method for large scale genomic analyses | yes |
| Powell et al. (2022) [53] | Integrates gene-to-phenotype biological model for bud outgrowth in simulation | yes |
| Sabadin et al. (2022) [54] | Develops genomic selection strategies for single-seed decent rice program | yes |
| Yang et al. (2022) [55] | Develops a method for modelling allele frequency change over years | yes |
| de Jong et al. (2023) [36] | Compares selection on GCA with different statistical models | yes |
| Fritsche-Neto et al. (2023) [56] | Compares different index selection approaches | yes |
| Jannink et al. (2023) [57] | Uses Bayesian optimization for resource allocation in two-part clonal program | yes |
| Krause et al. (2023) [58] | Compares linear mixed models to estimate realized genetic gains in a soybean program | no |
| Labroo et al. (2023) [32] | Develops hybrid breeding strategies for clonal programs | yes |
| Lubanga et al. (2023) [59] | Develops genomic selection strategies for a tea program | yes |
| Lanzl et al. (2023) [60] | Examines the impact of mating designs on estimation of genetic variance | yes |
| Oliviera et al. (2023) [61] | Analyses drivers of genetic change in a breeding program | yes |
| Platten et al. (2023) [62] | Compares three methods of QTL introgression into elite material | yes |
| Werner et al. (2023) [63] | Develops genomic selection strategies for a strawberry clonal program | no |
| Fritsche-Neto et al. (2024) [56] | Examines factors influencing the genetic gain in a rice hybrid program | no |
| Ferreira Azevedo et al. (2023) [64] | Examines factors influencing genomic prediction accuracy for visual score traits | no |
| Bancic et al. (2024) [] (under review) | Develops framework for simulating genotype by environment interaction | yes |

This paper serves as an introductory guide to stochastic simulation of plant breeding programs using the R package AlphaSimR [24]. Our target readers are plant breeding students and experienced breeders or researchers interested in integrating simulations into their toolkit. The readers are expected to have a foundational understanding of quantitative genetics and basic R coding skills. Through a detailed walk-through of an AlphaSimR wheat breeding program simulation, we share our past experiences and best practices in building a plant breeding program simulation. Additionally, we provide scripts for various breeding programs (self-pollinated, clonal, and cross-pollinated plants) and techniques (backcrossing, speed breeding, genomic selection, index selection, etc.) commonly used in plant breeding to highlight AlphaSimR’s flexible nature and plasticity of R scripting. These resources are intended for use as educational and reference materials that offer a starting point for designing bespoke plant breeding programs.

## 2 Material and Methods

This section consists of three parts. In the first part, we present the key steps for building a plant breeding simulation. In the second part, we summarize examples of breeding programs for self-pollinated, clonal, and cross-pollinated crop species. We consider different selection strategies for each breeding program. In the third part, we describe several examples of common breeding techniques that can be implemented within a breeding program or used alone for hypothesis testing. All examples are supported with AlphaSimR scripts, which are available in the supplement and at GitHub repository https://github.com/HighlanderLab/jbancicalphasimr plants. The purpose of these scripts is to empower new users, but also to encourage collaborative code enhancement over time with contributions from the readers. Throughout, we focus on a simple narrative without mathematical descriptions of the simulation steps involving quantitative genetics and statistics. We direct the reader to previous publications for these details [23, 24, 34].

We suggest that readers initially review the Material and Methods section to gain an understanding of the paper’s content and then proceed to the Results section, which provides a detailed description of a wheat breeding program simulation with key results. This simulation description is supplemented by a walk-through R Markdown vignette (see supplementary files LineBreeding.{Rmd,html}).

### 2.1 Key simulation steps

The process of building a breeding program simulation can be organized into seven steps:

1. Outlining the breeding program,
2. Specifying global parameters,
3. Simulating genomes and founders,
4. Filling the breeding pipeline,
5. Running the burn-in phase,
6. Running the future phase with competing scenarios, and
7. Replication and statistical comparison of the scenarios.

Detailed explanations of these steps are in the Results section 3 along with a practical example of a wheat breeding program and the corresponding walk-through R Markdown vignette in the supplement. Although the work with the seven steps may seem linear, it typically involves continuous revisions and iterations, which all require a considerable amount of time to complete. This approach to simulation is well supported by modern computing architecture using R, Python, or Julia environments. While the focus here is on AlphaSimR [24], the steps and best practices are also applicable to other simulation software. We used AlphaSimR version 1.5.3 and expect future versions to retain backward compatibility with the code presented.

### 2.2 Common breeding programs

This subsection introduces three distinct plant breeding programs for self-pollinated, clonal, and cross-pollinated crop species. Each breeding program is briefly introduced and supported with a corresponding AlphaSimR script (Table A1). The AlphaSimR scripts for each breeding program adhere to the steps outlined above and report trends in genetic mean, genetic variance, and selection accuracy.

#### 2.2.1 Breeding for self-pollinated species

Breeding programs for self-pollinated species include those of wheat, rice, barley, oat, and soybean [65–67]. The breeding cycle starts by crossing genetically diverse inbred parental lines to produce a segregating population. From these, candidate plants are selected for line development which lasts multiple stages with repeated growing and selection on traits of interest until they reach a desirable level of homozygosity. Line breeding programs leverage additive and epistatic genetic variation, and the release variety is a homozygous genotype, or inbred/pure line, which is multiplied through repeated selfing and bulking. There are several different breeding methods to produce inbred lines, including mass, single seed descent (SSD), and the pedigree method. These methods typically require several rounds of selfing to develop inbred lines. Advanced breeding programs use doubled haploid (DH) technology to replace the obtain true homozygous lines in less than two years after parental crosses, thereby substantially shortening the breeding cycle. In contrast to traditional methods which rely on F_2_ segregating populations, DH lines are directly generated from F_1_ plants. The process involves crossing F_1_ seed with an inducer line (e.g., wheat, corn) or employing anther or microspore culture (e.g., oilseed rape) to produce haploids. Haploid seeds (corn) or tissue cultures (other crops) are then grown as haploid plants and chromosome doubling is induced using a chemical agent (e.g., colchicine).

AlphaSimR scripts (Table A1, Programs 1 – 4) demonstrate the simulation of a hypothetical wheat breeding program employing the mass selection method, SSD method, pedigree method, and DH technology. The mass selection method is the simplest case, which involves recurrent selection of best individual plants or families, followed by intercrossing to form the next generation (or cycle) of selection with the aim to rapidly increase the frequency of desirable genes. The pedigree method is a slow and extensive method, which involves specific parental crosses to generate segregating F_2_ families to which both family and within-family selection are applied in each selfing round until plants are sufficiently inbred for multi-environment trials. Parent-progeny information is recorded for every candidate line to trace back to the initial parents with the aim to maximize variation produced in the next crossing cycle. The SSD method, typically performed in greenhouse/nursery conditions, involves selecting a single seed from each plant in every selfing round. This method achieves inbreeding of lines faster than the pedigree method and preserves genetic variability as every individual traces back to a different F_2_ individual. In each selfing round, the collected seed can be either bulked together (resulting in the breakup of the family structure) or maintained separately for each F_2_ plant using the Single-Hill procedure. Once sufficiently inbred, plants advance to multi-environment trials for further line development. A program shown in Figure 2 creates DH lines instead of selfing so that the line development and multi-environment trials commence immediately after their creation. The scripts highlight AlphaSimR’s flexibility to simulate these different breeding methods.

**Fig. 2.**
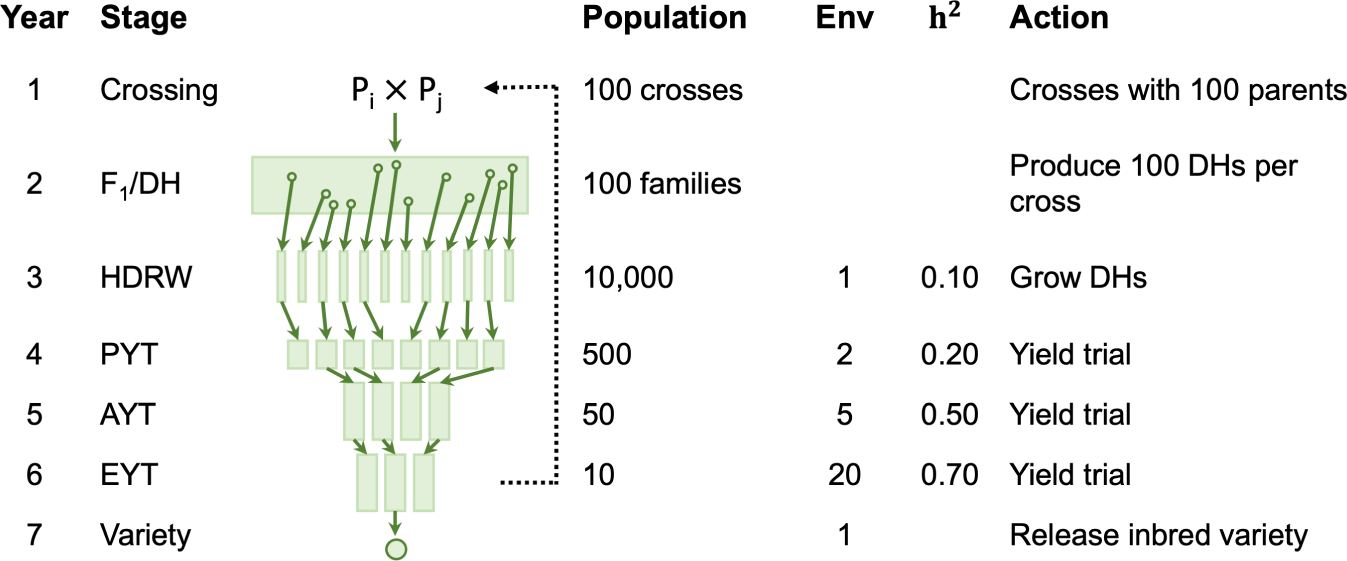
Key features of a hypothetical wheat line breeding program following [34]. Presented are key breeding stages with different populations (DH - doubled haploid, HDRW - headrow, PYT - preliminary yield trial, AYT - advanced yield trial, EYT - elite yield trial), each involving different numbers of plant genotypes in a population, environments (Env), heritability of yield, and action taken. The black dotted line indicates the stage where new parents are selected.

In addition to different selfing methods with phenotypic selection, we provide AlphaSimR scripts for two breeding scenarios that implement genomic selection with DH technology (Table A1, Programs 5 and 6). The first uses a genomic selection strategy within the context of a traditional wheat breeding program with concurrent product development and population improvement, while the second uses genomic selection to separate the components of population improvement and product development into the two-part strategy of [34]. To prioritize learning over maximizing genomic prediction accuracy, both genomic selection scenarios utilize limited training population sizes with only two years of historical data. As a result, the simulations run quickly on personal computers but may not yield optimal results.

#### 2.2.2 Breeding for clonally propagated species

Breeding programs for clonally propagated species include those of tea, strawberries, citrus trees, and bananas [65–67]. The breeding cycle starts by crossing genetically diverse parental genotypes through sexual reproduction to induce recombination and produce a segregating F_1_ population. Note that each cross produces a unique and distinct F_1_ seed (true seed) that has the potential to become a new variety as soon as it is made. The newly generated F_1_ seedlings are tested in an unreplicated trial to select promising F_1_ plants for multiplication through clonal/vegetative propagation (e.g. cuttings, corms, tubers, etc.) for subsequent testing in multi-year multi-environment trials. Clonal breeding programs leverage the total (additive and non-additive) genetic variation and the release variety is an outbred genotype, which is multiplied through repeated vegetative propagation.

AlphaSimR scripts (Table A1, Programs 7 – 9) demonstrate simulations of a hypothetical tea breeding program that uses phenotypic selection as shown in Figure 3 and two program variations using either pedigree-based or genome-based selection strategy as described in [59]. Both scenarios use limited training population sizes for faster execution on personal computers.

**Fig. 3.**
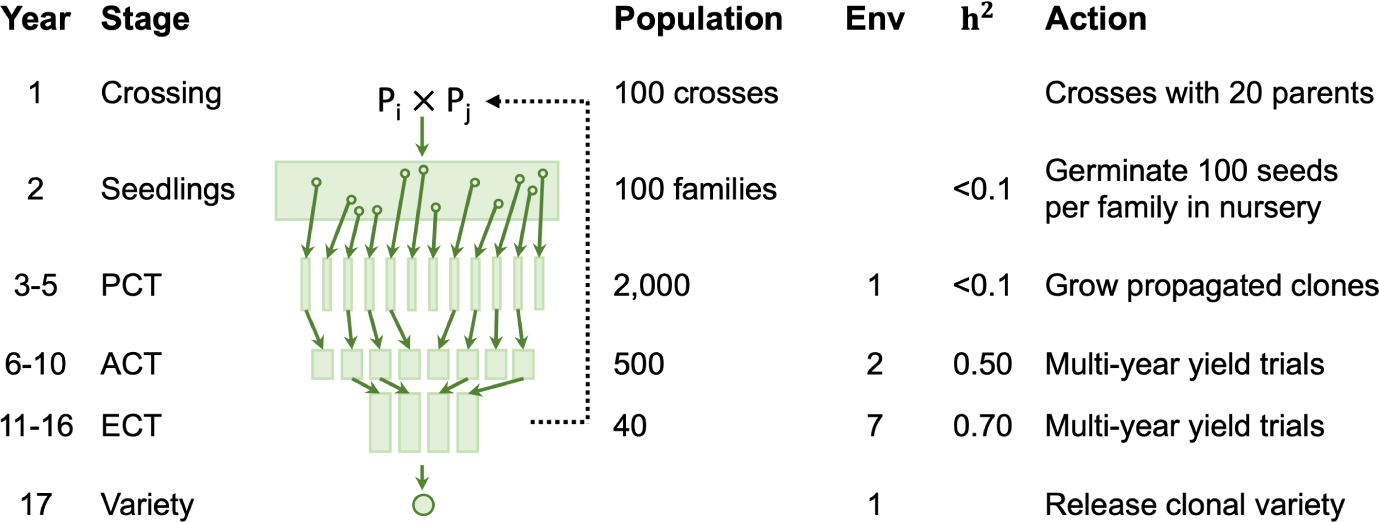
Key features of a hypothetical tea clonal breeding program following [59]. Presented are key breeding stages with different populations (PCT - preliminary clonal trial, ACT - advanced clonal trial, ECT - elite clonal trial), each involving different numbers of plant genotypes in a population, environments (Env), heritability, and action taken. The black dotted line indicates the stage where new parents are selected.

#### 2.2.3 Breeding for hybrid cultivars

Breeding for hybrid cultivars is used in self-pollinated and out-crossing species, including maize, sorghum, wheat, rice, sunflower, and various vegetables [65–67]. The process of developing hybrid cultivars uses reciprocal recurrent selection which consists of multiple steps. First, inbred lines within two (or more) genetically distinct heterotic groups are crossed to produce F_1_ populations. New inbred lines are derived from this population, commonly using DH technology. Second, selected lines from one heterotic group are crossed with tester lines from the opposite group, and vice versa, to determine their general combining ability (GCA; i.e. mean performance of an inbred in across hybrid combinations). Tester lines from each heterotic pool were previously selected for consistently high GCA and possessing key traits. Third, inbred lines with the highest GCA from each heterotic group are intercrossed to produce F_1_ hybrids. The crossing between heterotic groups results in F_1_ progeny with a high proportion of heterozygous loci that generally exhibit some level of dominance, which can generate heterosis (i.e. the F_1_ progeny outperforms the mean of their inbred parents). Hybrid programs therefore leverage both additive and dominance variation and the release variety is an outbred hybrid F_1_ genotype, which is multiplied through repeated crossing of known inbred parents.

AlphaSimR scripts (Table A1, Programs 10 – 12) demonstrate the simulation of a hypothetical maize hybrid breeding program with phenotypic selection as shown in Figure 4 and two scenarios with genomic selection scenarios. The first scenario applies genomic selection in the context of a traditional hybrid breeding program with concurrent product development and population improvement, while the second scenario applies genomic selection to separate the components of population improvement and product development into the two-part strategy of [34, 42]. Both genomic selection scenarios use limited training population sizes, employing two years of historical data, for faster execution on personal computers. The R scripts also highlight AlphaSimR’s features such as the specification of heterotic groups, performing testcrosses and hybrid crosses, as well as calculating, predicting, and selecting on GCA.

**Fig. 4.**
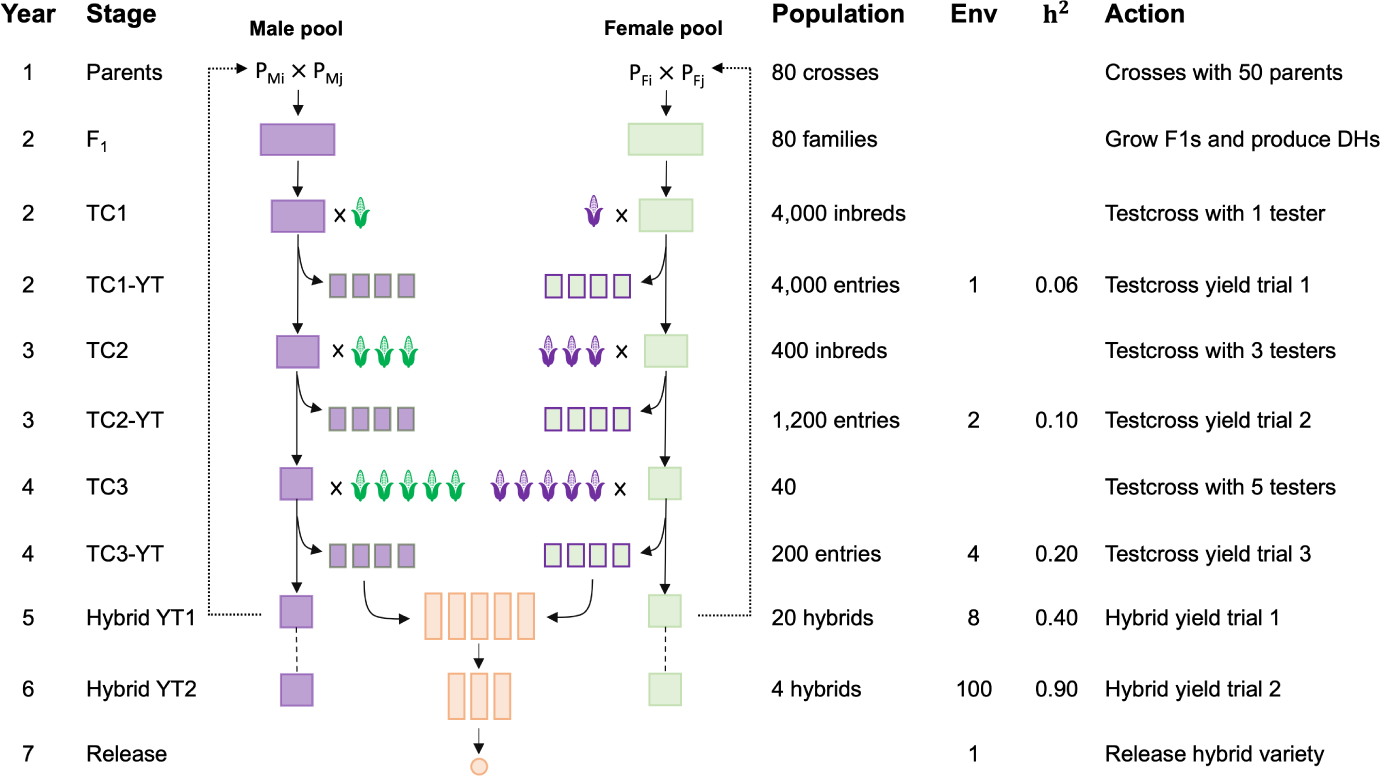
Key features of a hypothetical maize hybrid breeding program following [12]. Presented are key breeding stages (TC - testcross, TC-YT - testcross yield trial, Hybrid YT - hybrid yield trial), each involving different numbers of plant genotypes in a population, environments (Env), heritability, and action taken. The black dotted line indicates the stage where new parents are selected.

### 2.3 Common breeding techniques

In this subsection, we briefly describe several examples of common techniques applied in plant breeding. AlphaSimR scripts (Table A2) demonstrate these techniques independently, but they can also be integrated into breeding program simulations.

*Example 1: Trait introgression*: this technique facilitates the transfer of a desired trait, such as a major QTL allele or a transgene, from a donor parent into the superior genetic background of the recurrent parent [12]. Typically, breeders aim to recover the genome of the recurrent parent while introducing the new gene of interest into the resulting line. This process can be achieved through phenotypic selection but can be expedited with marker-assisted selection. AlphaSimR script (Table A2, Technique 1) demonstrates the backcrossing of an allele at a single locus, representing a trait of introgression, into the background of the recurrent parent with marker-assisted selection.

*Example 2: Speed breeding*; this technique shortens the breeding cycle by manipulating plants under controlled greenhouse conditions to accelerate their growth and development. For example, speed breeding allows up to six generations of wheat plants to be grown annually under ideal conditions, compared to typical two generations in outdoor conditions with shuttle breeding between two regions [14]. Speed breeding can also be applied to accelerate selfing and for rapid cycling with DH technology in two-part breeding programs. AlphaSimR script (Table A2, Technique 2) demonstrates a simulation of four generations per year. Also see its implementation for rapid cycling in a two-part wheat breeding program (Table A1, Program 6).

*Example 3: Genome editing*: this technique allows for precise modification of plant genomes by targeting specific loci (e.g., editing QTL with CRISPR-Cas) to introduce or modify specific traits. AlphaSimR script (Table A2, Technique 3) demonstrates the editing of a single locus in the entire population. *Example 4: Multi-trait selection with selection index*: this approach is commonly used for the simultaneous selection of multiple traits, each with an assigned (economic) weight. AlphaSimR script (Table A2, Technique 4) demonstrates how to simulate traits with different genetic architectures and different correlations, and how to perform näıve or Smith-Hazel index selection. The selection index may be stored in the miscellaneous slot of the population. The miscellaneous slot provides space to store additional information, such as selection index values, for each individual to be used during selection (see Technique 5 in Table A2).

*Example 5: Mating plans*: planned crosses serve multiple purposes in plant breeding, including the generation of novel genetic variation, estimation of genetic variances through experimental designs, formation of heterotic groups, and mapping of major QTLs. AlphaSimR script (Table A2, Technique 6) demonstrates several mating designs, including biparental crosses, testcrosses, half- and full-diallel crosses within both a single and two populations, and more. AlphaSimR script (Table A2, Technique 7) for GWAS simulation is also provided.

*Example 6: Genomic models*: there are a variety of models that can be fit to genomic data. Models can include only additive effects, or can include both additive and dominance effects. Models can also incorporate population specific effects, known as general combining ability and specific combining ability, applicable to hybrid breeding. AlphaSimR script (Table A2, Technique 8) for such different models is provided.

*Example 7: Setting heritability*: there are several ways that heritability can be set in AlphaSimR. AlphaSimR script (Table A2, Technique 9) demonstrated setting narrow-sense heritability, broad-sense heritability, error variance, and number of replications.

These examples highlight a small number of potential applications to demonstrate various breeding techniques, as well as the versatility of combining AlphaSimR functionality with R scripting. We encourage users to consider other potential applications, such as simulating high-throughput phenotypic, omic, and environmental data and analyzing it with advanced statistical and machine learning methods.

## 3 Results

In this section, we demonstrate the key simulation steps outlined in Material and Methods section 2.1 with a wheat breeding program as a case study. We simulate a program with phenotypic selection and a program with genomic selection. Each step is accompanied by a comprehensive description and details specific to the wheat breeding program example. We suggest that readers process the information provided below in parallel with the R Markdown vignette (files LineBreeding.{Rmd,html}). The vignette mirrors steps 3.2 - 3.5 over a single year of the breeding program, including some of the text for completeness. The complete multi-year simulations of the wheat breeding program are available as AlphaSimR scripts.

### 3.1 Outlining the breeding program

Before initiating a breeding program simulation, it is imperative to lay out its key stages and actions, which usually requires gathering essential biological, logistical, statistical, and agronomic details from breeding program managers and specialists. The information gathered in this step is the foundation for translating a breeding program into AlphaSimR scripts. Considerations include:

- Breeding objectives: which key traits are being improved.
- Type of mating: whether crop is selfing, clonal, or cross-pollinating.
- Numbers per breeding stage: how many individual genotypes and how many observational trials.
- Heritabilities: what are the narrow-sense heritabilities for every selection that is performed.
- Type of selection: what kind of selection is performed, such as individual-,family-, or testcross-based.
- Genomic features: information about the crop’s genome and its evolution, possible knowledge about major and minor QTL.
- Breeding population features: information on genetic means, variances, correlations between traits, number of initial parents, or level of inbreeding and heterosis.
- Biological restrictions: what biological constraints exist in the crop such as self-incompatibility mechanism, a limited seed multiplication rate, or the need for seed vernalization.
- Logistical constraints: what logistical constraints exist in the crop or the modelled breeding program, such as the number of generations that can be achieved each year, the number of available testing locations, the number of parental crosses, or the number of candidate genotypes produced.
- Program specifics: information on data generation, statistical models used, and breeding region.

To establish an overall budget for the baseline breeding program, we obtain information on the costs for various actions, such as maintaining plants in the nursery, parent crossing, DH production, genotyping, and phenotyping. With a budget in hand, we can ensure fair comparisons of new strategies under comparable cost constraints.

In Figure 2, we present the key features of a wheat breeding program, including the number of individuals at each stage, heritability of trait, and selection actions. In Table 2 we outline approximate costs for both phenotypic and genomic selection in such breeding programs. Here, we used approximate costs for demonstration purposes. We constrained costs of the genomic program to those of the phenotypic program ($503,500) by reducing the number of DH individuals produced per parental cross from 100 to 89 and skipping the headrow stage. We stress, however, that changing the costs of actions will have a profound impact on the choice of the most optimal strategy. In the Results sub-section 3.7, we also present the outcome of an unconstrained genomic selection program with a total cost of $553,500 that shows the achievable gain at the maximum capacity of the program. Other ways of reducing costs, such as decreasing the extent of field testing or removing particular stages, may also offset the extra costs of implementing a new method. Such changes should be logistically feasible and informed by input from the breeding program manager.

**Table 2.** Cost comparison between the phenotypic breeding program and the cost-constrained genomic selection program (DH - doubled haploid, HDRW - headrow, PYT - preliminary yield trial, AYT - advanced yield trial, EYT - elite yield trial).

| Action | Cost (\$) | Env | Phenotypic | | Genomic | |
| --- | --- | --- | --- | --- | --- | --- |
| | | | # Units | Cost (\$) | # Units | Cost (\$) |
| Cross | 30/cross | / | 100 | 3,000 | 100 | 3,000 |
| Grow F<sub>1</sub>s | 30/plant | / | 100 | 3,000 | 100 | 3,000 |
| Make DHs | 30/plant | / | 10,000 | 300,000 | 8,900 | 267,000 |
| Genotype | 15/plant | / | / | / | 8,900 | *133,400* |
| HDRW | 10/plot | 1 | 10,000 | 100,000 | / | / |
| PYT | 20/plot | 5 | 500 | 50,000 | 500 | 50,000 |
| AYT | 50/plot | 15 | 50 | 37,500 | 50 | 37,500 |
| EYT | 50/plot | 20 | 10 | 10,000 | 10 | 10,000 |
|  |  |  | **Total** | 503,500 | **Total** | 504,500 |

*Note:* Text in italics indicates the additional cost of genotyping in a genomic selection program, offset by the smaller number of DH individuals and skipping the HDRW stage.

### 3.2 Specifying global parameters

Translating a breeding program into AlphaSimR code begins with the definition of simulation parameters based on the information gathered in the previous step. We specify values such as the number of crosses, progeny per cross, and individuals evaluated in various stages as well as genetic features, including variances, the number of QTL, degree of dominance, and demographic history. In Table 3, we provide a list of parameters, definitions, and the values used for simulating the wheat breeding program. We encourage users to use clear and consistent variable naming in AlphaSimR scripts (e.g., using camelCase or PascalCase notation) to improve code readability and debugging. The same applies for the R code style [68].

**Table 3.** Global parameters and definitions with values used for simulating a wheat breeding program (HDRW - headrow, PYT - preliminary yield trial, AYT - advanced yield trial, EYT - elite yield trial, GS - genomic selection).

| Parameter | Definition | Value |
| --- | --- | --- |
| nReps | Number of simulation replications | 10 |
| nBurnin | Number of years in the burn-in phase | 20 |
| nFuture | Number of years in future phase | 20 |
| nQTL | Number of QTL per chromosome | 20 |
| nSnp | Number of SNPs per chromosome | 400* |
| initMeanG | Initial population mean genetic value for yield trait | 0 |
| initVarG | Initial population genetic variance for yield trait | 1 |
| initVarGE | Initial GxE interaction variance for yield trait | 2 |
| varE | Yield trial error variance for yield trait | 4 |
| nParents | Number of parents to start a breeding cycle | 50 |
| newParents | Number of new parents each breeding cycle | 50 |
| nCrosses | Number of crosses among parents to start a breeding cycle | 100 |
| nDH | Number of DH individuals produced per cross | 100/89 |
| famMax | Maximum number of DH individuals per cross to enter PYT | 10 |
| nPYT | Number of entries in PYT | 500 |
| nAYT | Number of entries in AYT | 50 |
| nEYT | Number of entries in EYT | 10 |
| repHDRW | Effective replication in HDRW | 4/9 |
| repPYT | Effective replication in PYT | 1 |
| repAYT | Effective replication in AYT | 4 |
| repEYT | Effective replication in EYT | 8 |
| startTP | Year to start collecting training records for GS | 18* |

*Note:* Values in italics represent the reduced number of DH individuals to offset the cost of implementing genomic selection and \* indicates added parameters for genomic selection program.

Careful selection of global parameter values is crucial to ensure that the resulting properties of a simulated population, such as population structure, level of heterosis or inbreeding, and genetic or phenotypic variation, reflect those of a real breeding program. Parameter values can be chosen based on estimates from prior analyses of real data or long-term trends, trial and error, or practical experience. Deciding on parameter values may require simulating the founder parents and breeding program multiple times until realized population properties resemble expectations. Users must be cautious to avoid biasing parameters to favour a specific scenario in the study.

### 3.3 Simulating genomes and founders

In the next step, we initialize a founder population for simulation, which includes simulating or importing founder haplotypes and specifying trait-related and other features. Many agricultural species have experienced significant and repeated bottlenecks during domestication and selective breeding, resulting in distinct population structure and linkage-disequilibrium patterns. AlphaSimR embeds MaCS software [69] to generate founder genomes through a backward-in-time (coalescent) simulation. The software creates genealogical trees based on recombination and demographic parameters and drops mutations onto the trees, producing whole-chromosome haplotypes that form genomes and genotypes of founder individuals.

In AlphaSimR, we can create founder genomes via three approaches: i) selecting from pre-defined species’ demographic histories, ii) creating custom species histories, or iii) importing externally obtained haplotypes (Table A2, Technique 10 - 11). For quick tests, we can sample founder haplotypes randomly from a Bernoulli distribution or specify a ”generic” species to rapidly generate founders with a somewhat realistic demography for an agricultural species. To create a custom species demography, we need to specify parameters such as the number of chromosomes, loci, mutation and recombination rates, and demographic history in terms of changes in effective population size over time. In our experience, setting the effective population size to an empirically obtained current value in the range of 10 to 100 produces genomes with far fewer segregating sites than observed in real data. Increasing the effective population size over time in the population’s history addresses this issue at least partly [70]. It is important to note that simulating realistic demographic histories is challenging, both due to the lack of knowledge and excessive run times of complex demographic simulations. We have observed that different methods of simulating genomes and their parameters have an impact on the overall performance of breeding programs, but they generally do not impact relative performance of different scenarios. Although these aspects of species evolution and domestication are only partially known, community-curated population genetics models for various species, facilitated by software such as stdpopsim [70], are becoming available and will enhance user convenience and ensure reproducibility. stdpopsim can work with both the backward-in-time simulator msprime [71] and the forward-in-time simulator SLiM [72] with a standard way of specifying demographic models [73], with an R front end [74]. Finally, we can import externally obtained haplotypes, either simulated with other software or derived from observed genotypes, which must be phased into haplotypes. Although some users might assume that importing their genomic data is sufficient to replicate observed phenotypic variation in simulations, this is generally not the case because phased genotypes for SNP markers usually lack QTL information, let alone their effects on traits of interest.

AlphaSimR enables simulations with haploid, diploid, and polyploid species. Some functions even enable work with variable ploidy within a simulation. For allopolyploid species, we recommend specifying their genome as diploid with an increased number of chromosome pairs that show disomic inheritance. For autopolyploid species, we advise specifying their genome as polyploid with the actual number of chromosome groups that show polysomic inheritance. While AlphaSimR’s forward-in-time simulation adequately accommodates variable ploidy, the MaCS call for backward-in-time simulation of autopolyploid genomes requires appropriate modifications [32, 75].

After creating the founder population, we can add trait-related features to it. We can add single or multiple correlated traits or uncorrelated traits with shared or distinct QTL and specify the distribution of QTL effects, means, and variance-covariance matrices for QTL effects (additive, dominance, and epistatic). It is also possible to import a trait with manually specified QTL and their effects (see Technique 10 in Table A2). At this stage, we can also specify sex (if relevant for species), one or more SNP arrays (including QTL or not), tracking pedigree, tracking recombination and segregation events, sex-specific recombination, the frequency of quadrivalent pairing in autopolyploids, or the parameters of the gamma-sprinkling recombination model [76].

In our wheat breeding program, we simulated a founder population with a pre-defined wheat species demographic history. For demonstration purposes, we only consider a single trait that represents grain yield. In Table 4, we summarize the founder genomes and trait parameters used in the simulation.

**Table 4.**
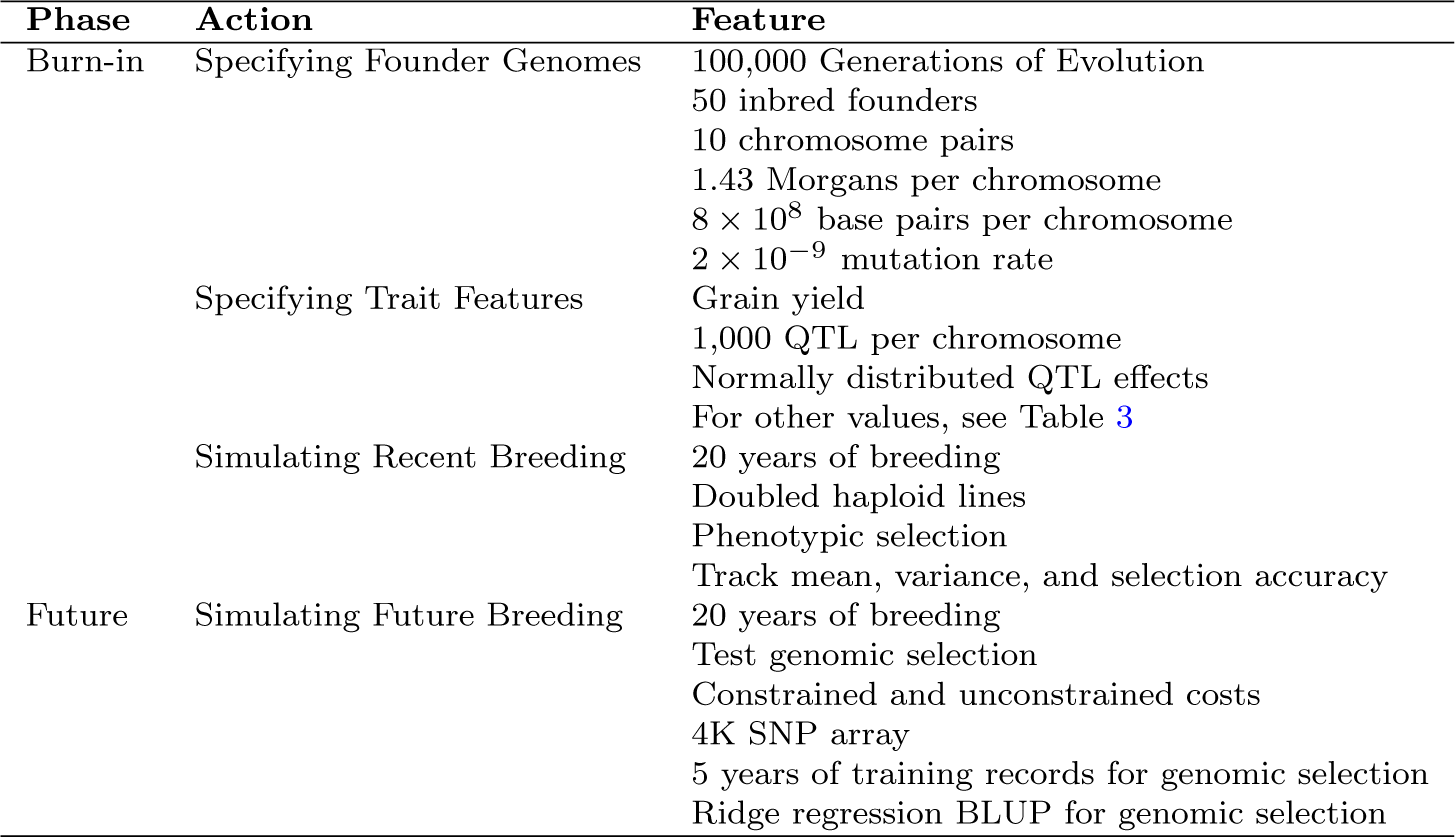
Summary of simulation features. Presented are steps taken in the burn-in and the future phase pertaining to the genome, founders, and trait.

### 3.4 Filling the breeding pipeline

Once the stages of the breeding program are defined and the founders are simulated, the next step is to code the filling of each breeding stage with a distinct population, or cohort, that relates back to its parents. The filling process is necessary to mimic the overlap of different breeding cycles in real breeding programs. In practical terms, this means that, at any given time, a breeding program will consist of multiple cohorts that progress through stages of the breeding pipeline. We initiate the filling by crossing the founder parents and progressing the new populations of genotypes through the stages of our breeding program, saving a cohort at a different breeding stage during each iteration.

Figure 5 illustrates the filling process for our wheat breeding program example. The program has six stages, which means that we need six distinct cohorts. The first cohort (orange) progresses through all stages and is saved in the sixth (final) stage (EYT), making it the oldest cohort. The second cohort (yellow) progresses to the fifth stage (AYT), making it the second oldest cohort. This process repeats six times until all stages are populated with distinct cohorts (see year 0), with the last being the most recent (dark gray). Despite using the constant founder parents, unique cohorts arise due to different crossings, randomness of genetic inheritance, and selections in each cohort. As the burnin phase commences in year 1 (described in the next sub-section), all cohorts, excluding the oldest which has already completed all stages, will simultaneously progress diagonally down the pipeline each year, also generating a new cohort in the earliest stage (e.g. Cohort 7, purple).

**Fig. 5.**
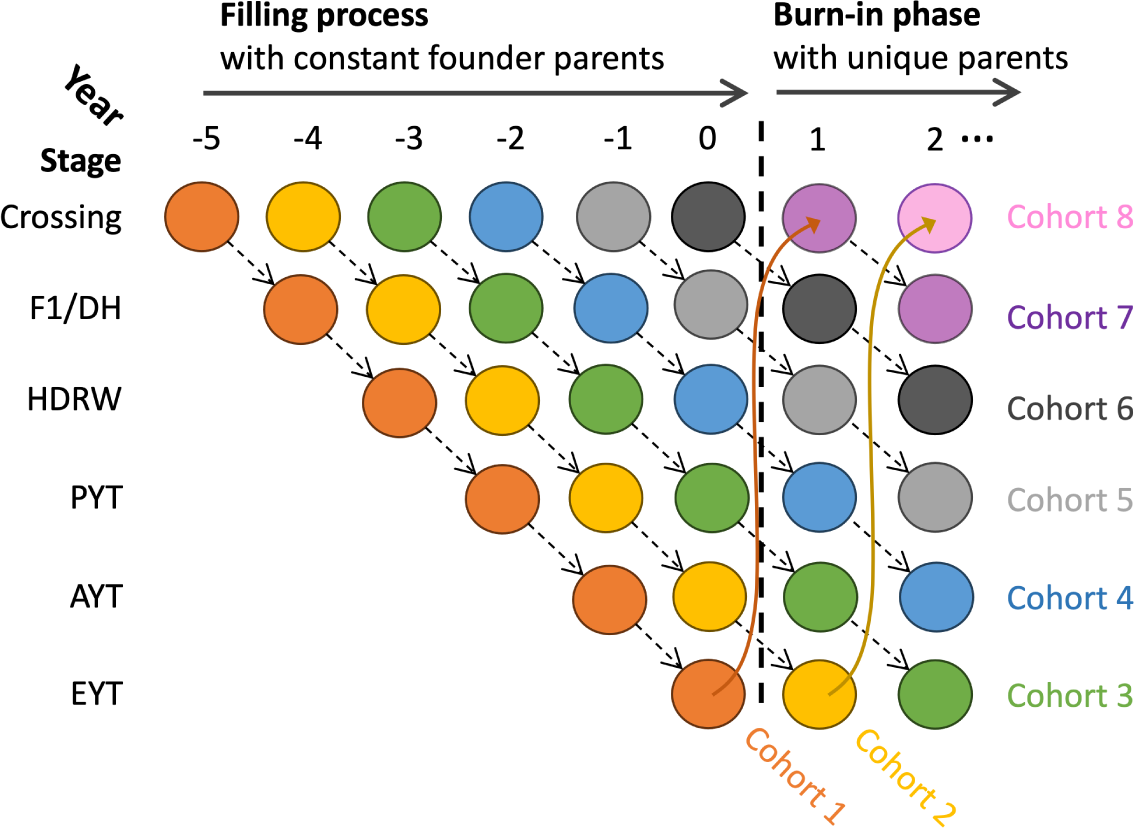
An illustration of filling the breeding pipeline and the beginning of the burn-in phase in the simulated wheat breeding program. *Filling phase*: initiates cohorts of the breeding program using constant founder parents over six iterations. In the first iteration, Cohort 1 (orange) passes to the sixth stage (EYT). In the second iteration, Cohort 2 (yellow) passes to the fifth stage (AYT), and so on. After the last iteration, each stage contains a population of a different cohort. *Burn-in phase*: in the breeding cycles after filling, each cohort progresses diagonally from left to right to the next stage of the breeding program. The cohorts are transitioning through the breeding program simultaneously, and a new cohort is generated each year (e.g., cohorts 7 and 8).

### 3.5 Running the burn-in phase

Before comparing different breeding scenarios, a burn-in phase must be simulated to serve as a common point from which scenarios can be compared (see the next subsection 3.6). This phase is necessary for two reasons: i) representing a historical breeding period preceding the implementation of a new strategy; and relatedly ii) achieving realistic evolution of linkage-disequilibrium across the genome under selection to ensure genetic structure and diversity mimic those of a real breeding program. In our wheat breeding program, the burn-in phase lasts 20 years and uses a phenotypic selection program as the baseline (Figure 2 and Table 4). Following the process shown in Figure 5, each new burn-in year involves i) retiring the oldest cohort that reached the final EYT stage in the previous cycle and selecting the best individuals as parents, ii) creating a new cohort, and iii) progressing cohorts one stage ahead. Throughout the burn-in phase, we collect historical records for training a genomic prediction model in the future phase when genomic selection is introduced and compared to phenotypic selection program.

In the burn-in phase we also begin collecting simulation data and its summary statistics. To facilitate this process, we initialize empty data frames that serve as containers for variables. These data frames may include columns such as the scenario name, replication number, stage, simulation year, specific simulation input parameters, and variables monitored throughout the simulation. Note that AlphaSimR provides many integrated functions for computing common summary statistics, such as mean and variance of genetic or phenotypic values, which is useful for exploring results after simulation. The data collected can be used to report results at specific stages or across all breeding stages over time, as well as to identify potential coding errors. Sometimes, we might wish to store entire populations, which can be achieved by creating list objects, in which each population is stored as a separate list element that can be accessed later in the simulation. When saving whole population objects with large number of individuals, care must be taken regarding computer memory capacity.

### 3.6 Running the future phase with competing scenarios

After the burn-in phase, we start with the evaluation of various breeding scenarios, each implementing a new feature. The choice and properties of these scenarios depend on the purpose of our simulation study. In practice, breeders may use the future phase to inform their decision-making process. For example, determining the optimal crossing design to produce large genetic variation among F_1_s, identifying the ideal number of parents to maximize genetic gain, or optimizing the number of observational trials at a particular stage. A strategic approach to testing this case is to select reasonable as well as extreme values from the parameter space to gain intuition of how the parameters impact outcomes. Typically, varying one parameter at a time is the most effective method to understand various nuances and avoid confounding different factors. In our wheat breeding simulation, we compare phenotypic selection and genomic selection (Table 4). In Figure 6, we present the implementation of genomic selection for our example to: i) reduce the program’s length by advancing individuals directly from the DH stage to the PYT stage, ii) to reduce the program’s generation interval by selecting parents from the earliest stage (DH), as opposed to the EYT stage (Figure 2), and iii) to improve selection accuracy in the DH stage with prediction. We note that this is just one potential implementation of genomic selection and typically several different scenarios would be tested to identify the most optimal one.

**Fig. 6.**
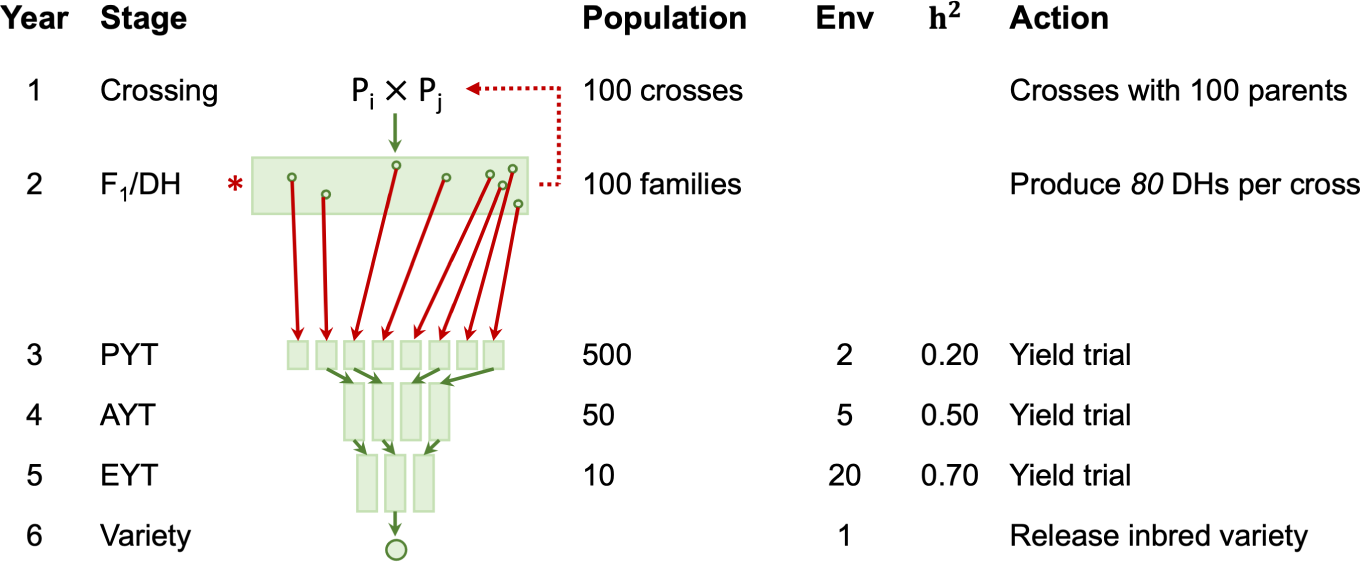
Key features of a hypothetical wheat line breeding program implementing genomic selection. Presented are key breeding stages with different populations (DH - doubled haploid, PYT - preliminary yield trial, AYT - advanced yield trial, EYT - elite yield trial), each involving different numbers of plant genotypes in a population, environments (Env), heritability of yield, and action taken. The actions in red highlight the changes enabled by genomic selection and are described in Results sub-section 3.6.

Evaluating a new feature may require the use of external software. If this software is an R package, then we can integrate code directly into the AlphaSimR script. If the software is not an R package, we need to pause the simulation each time external software is required. We export the dataset in a compatible format for analysis, analyse it, import the software’s output back into R to perform the required task on the target breeding population, and then resume the simulation. This process can be automated using R’s functionality to interact with the operation system (such as system() R function). In our wheat genomic selection simulation, we use an internal AlphaSimR ridge regression function to train a genomic prediction model and to assign estimated breeding values to individuals in a population. Alternatively, we could use other software (e.g., ASReml or blupf90) for this task, which requires creating a dataset in the correct format (see the pedigree-based tea program, Program 8 in Table A1). We advise users to develop custom functions for data transformation and storage within the R environment or on the local disk. Several functions in AlphaSimR can assist with tasks such as extracting SNP or QTL genotype or haplotype matrices and phenotype data from a target population or preparing export data (e.g., for input to plink).

The collection of descriptive and simulation output variables continues into the future phase to allow scenario comparison. In the wheat breeding programs, we track the genetic mean, the genetic variance and the selection accuracy over a 40-year simulation period. The first two measures for both phenotypic selection and genomic selection program are collected from the earliest common DH stage, where the number of individuals and genetic variation are the largest. Selection accuracy, measured as the correlation between estimated (genomic or phenotypic) and true genetic values, is recorded upon advancing individuals from HDRW or DH to the PYT stage for phenotypic and genomic selection, respectively. Note that we collect only a single value for each measure every year for demonstration purposes. In practice, we collect these measures at every stage of the breeding program for diagnostic purposes. We will discuss the results in the next step.

### 3.7 Replication and breeding program comparison

Outcomes of simulated breeding programs are stochastic due to the inherent randomness of genetic inheritance and environmental factors. To capture the distribution of potential outcomes, we need to replicate the entire simulation, including founders, burn-in, and future phases. Recent study highlights that founder simulation alone can significantly impact the outcome of a breeding strategy [57]. Replication serves at least two different purposes: (i) to capture and understand the key sources of variation and their impact on the breeding program; and (ii) to calculate a summary (e.g. mean and variance) of tracked statistics (genetic mean, genetic variance, accuracy, etc.) over multiple simulation outputs, necessary to determine whether there are significant differences between scenarios. Comparison of summary values for the tracked statistics between scenarios is often the goal of simulation studies - to indicate average difference between the scenarios. However, sometimes interest lies in the magnitude of variation between individual replicates to gain insight how an individual instance of a breeding program could deviate from the average. The number of replications will depend on the aims of the study, the level of precision desired, the complexity of a simulation, and required compute resources. Our experience has shown that at least 10 replicates are required to achieve somewhat stable distributions of tracked statistics and their means, but preferably more to increase the precision. Also, to control for variation between burn-in phase replicates, we represent and test future scenarios by centering their starting values to the mean across the burn-in replicates. A good practice to overcome long compute times is to parallelize multiple replicates on a cluster computer. We first run the burn-in phase and save a snapshot of the R environment, including states of variables and populations at the end of the burn-in. Next, parallel R sessions are started by importing these snapshots to begin the future phase for each scenario and run independently from the same starting point (within a replicate), with results saved and collated afterwards. Doing so, each burn-in replicate is used by several scenarios at the same time, and elapsed time of running all scenarios is significantly reduced.

Figure 7 shows the trends of genetic mean, genetic variance, and selection accuracy over the 40-year simulation period for phenotypic and genomic selection in wheat breeding. The results show a clear advantage of genomic selection over phenotypic selection at the expense of rapid loss in genetic variance. The positive difference in genetic mean is due to a shorter breeding program and generation interval, as well as increased selection accuracy in breeding stages where genomic selection was applied. These results are consistent with previous publications listed in Table 1, which interested readers are referred to for detailed discussions. Figure 7 shows trends in genetic gain for individual simulation replicates (light lines) and their mean (solid line) in both the burn-in phase and the future phase. Variability around the mean line not only reflects the inherent stochasticity of the simulation but also highlights the importance of replication. Looking at individual replicate lines in the future phase, one can observe multiple re-rankings among the three scenarios, particularly between the constrained and unconstrained genomic selection programs. Our results suggest that investing an additional $49,000 for genotyping 2,000 more plants in the cost-unconstrained genomic selection scenario can increase genetic gains compared to the cost-constrained genomic selection scenario. A paired t-tests between 10 replicate values in the final year confirm significant differences between all three scenarios. It is worth noting that, for demonstration purposes and for short running times, all our genomic selection scenarios restrict the size of training populations, with only two years worth of data. This factor may have influenced the simulation outcome, further highlighting the importance of clearly deciding on the parameter space of a simulation.

**Fig. 7.**
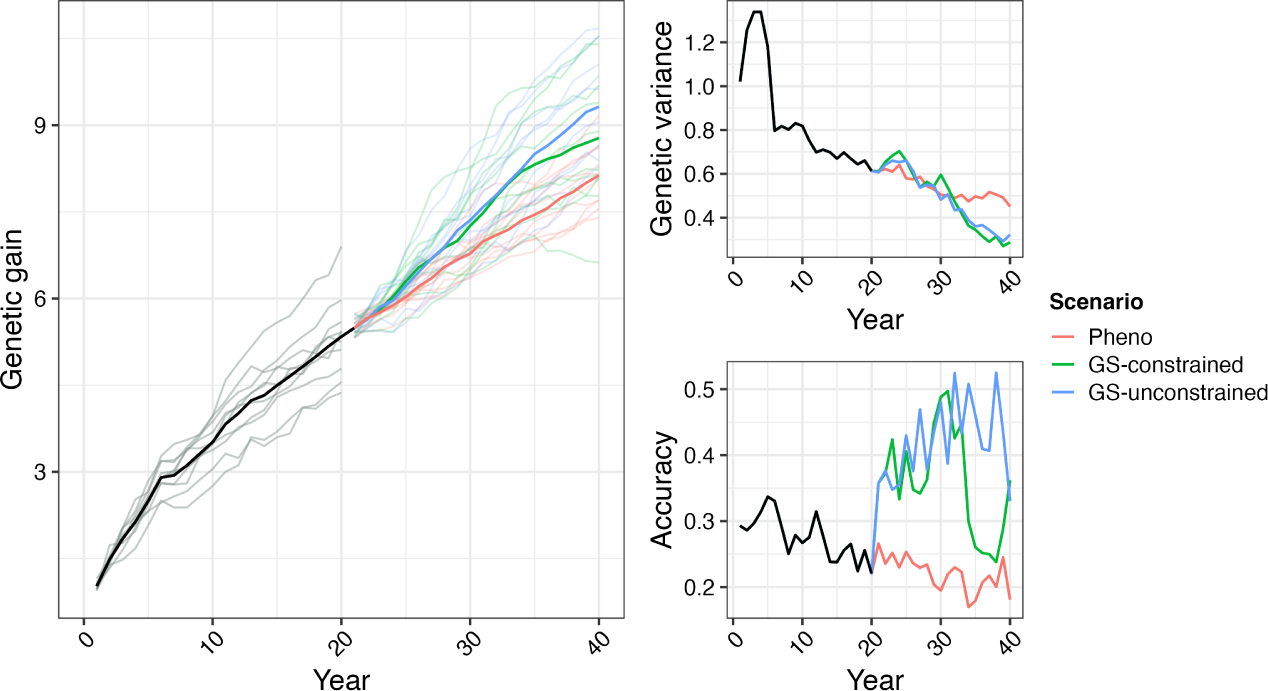
Summary of the wheat breeding program simulation outputs over a 40-year period. Presented in figure are genetic mean (left), genetic variance (top-right), and selection accuracy (bottom-right) for phenotypic breeding program and genomic selection (GS) program with constrained and unconstrained costs. Black line represents the burn-in phase and coloured lines represent scenarios of the future phase. For genetic mean, thick dark lines indicate the average across 10 simulation replicates, and thin light lines indicate individual replicates. For clarity, we omit individual replicates for genetic variance and selection accuracy. Scenarios of the future phase, start at year 20, with the starting value presented as centered on the average of the final burn-in year.

## 4 Discussion

Stochastic simulations have a great potential to improve plant breeding programs to meet the growing food demands of an expanding human population amidst climate change. Previous literature has highlighted their usefulness in refining and restructuring existing breeding programs, optimizing the implementation of emerging technologies, answering basic research questions, and designing new breeding strategies (see some examples in Table 1). However, there remains a lack of literature that provides practical guidance on implementing and deploying such simulations. This paper aims to fill this gap as an introductory guide to stochastic simulation of plant breeding programs using the R package AlphaSimR [24]. Through a detailed walk-through of a wheat breeding program example, we share our past experiences and best practices in building a plant breeding program simulation. Additionally, we provide AlphaSimR scripts for various breeding programs (self-pollinated, clonal, and cross-pollinated plants) and techniques (backcrossing, speed breeding, genomic selection, index selection, etc.) to highlight AlphaSimR’s flexible nature coupled with the plasticity of R scripting. These resources are intended as educational and reference materials that offer a starting point for designing bespoke plant breeding programs or studies of them.

While this paper covers general topics of building a plant breeding simulation, there are several areas of such simulations where development is ongoing or needed. Here, we highlight five areas of development.

- *Genotype by environment (G×E) interaction* complicates selection in plant breeding. Currently, AlphaSimR offers two methods for modeling G*×*E: a formal G*×*E trait that models a simple form of G*×*E driven by a single latent environmental covariate or manual process of modeling G*×*E using multiple correlated traits. A scalable and reproducible framework for simulation of realistic G*×*E interaction is being developed that allows users to specify between-environment correlations across years before the simulation commences [18]. This will enable users to simulate multi-environment field trials of different G*×*E interaction complexities within simulation.
- *Optimizing large parameter space* is one of the inherent challenges of stochastic simulation studies, now that we can easily design and parameterise large number of breeding scenarios. Typically, users can explore only a limited number of parameter sets to find the optimal scenario. This process can be time-consuming and computationally expensive and hence, often only a subset of scenarios is tested. Recently, Bayesian optimization-type approaches have been used to manage the exploration of a large set of possible breeding scenarios [57, 77, 78].
- *Genome simulation* is often overlooked due to the lack of genomic and demographic information. With rapid advances in genomics, the characterization of species’ genomes and demographies of multiple populations is becoming available [70, 73]. With this information, more realistic simulations of the founder genomes will be feasible [71, 72]. In the future, we also envision the possibility of importing comprehensive representations of real data into simulations [79].
- *Multi-population simulation* is important for some species, such as forages, where breeding programs perform group selection rather than individual selection. We are expanding AlphaSimR functionality to work with entries that are themselves populations, as has already been done in an insect breeding simulator [80].
- *Model-specified phenotype simulation* is required to specify custom sources of variation. Currently, AlphaSimR limits the user as to how many fixed terms and random terms with pre-defined distribution can be added to the simulation of a phenotypic value. While there are existing attempts to add additional sources of variation, for example, field trial layouts [81], a more flexible approach is needed for simulating phenotypic values based on user-defined model formulae and user-provided data layout. This will allow users to specify custom fixed and random effects such as location, genotype by location, genotype by location by year, and various linear and non-linear functions associated with environmental covariates. While this can be currently achieved by a combination of AlphaSimR population’s misc slot and R scripting, it requires programming skills.

As these developments continue, plant breeding simulations will become an increasingly powerful tool to accelerate the genetic improvement of crops. The readers are encouraged to actively contribute to this effort by participating in the free online edX course Breeding Programme Modelling with AlphaSimR (https://www.edx.org/course/breeding-programme-modelling-with-alphasimr), consulting the technical documentation (https://cran.r-project.org/web/packages/AlphaSimR/AlphaSimR.pdf), participating in user discussions and issues on GitHub (https://github.com/gaynorr/AlphaSimR), and extending the existing literature outlined in Table 1. Ultimately, every breeding program would benefit from having a corresponding virtual/digital twin as a testing ground for new ideas before incorporating them into practice. By integrating the use of simulation software into plant breeding education and practice, future plant breeders will be equipped with this powerful tool.

## Author Contributions

GG initiated the project. JB and PG wrote the manuscript. RCG, JB, and PG developed R scripts. All authors have reviewed the R scripts and manuscript and approved the final version.

## Supporting information

Scripts

## Acknowledgments ORCID of Authors

Jon Bančič https://orcid.org/0000-0001-7077-7163

Philip Greenspoon https://orcid.org/0000-0001-6284-7248

Chris R. Gaynor https://orcid.org/0000-0003-0558-6656

Gregor Gorjanc https://orcid.org/0000-0001-8008-2787

## Code availability

AlphaSimR scripts for all the breeding programs and techniques presented in this manuscript (Tables A1 and A2) are available in the supplement and at GitHub repository https://github.com/HighlanderLab/jbancic alphasimr plants.

## Appendix A Inventory of AlphaSimR code

This appendix contains an inventory of the AlphaSimR scripts we provide for a variety of breeding programs and techniques - available in the supplement and at GitHub repository https://github.com/HighlanderLab/jbancicsalphasimrplants. In Tables A1 and A2 we indicate the folder and file path for each breeding program and technique.

**Table A1.** List of breeding programs and methods presented in this paper and their file paths.

| Program | Description | Directory Path |
| --- | --- | --- |
|  | Line breeding | 01_LineBreeding/ |
|  | - Phenotypic selection | 01_LineBreeding/01_PhenotypicSelection |
| 1 | - Mass selection | 01_LineBreeding/01_PhenotypicSelection/01_MassSelection/ |
| 2 | - Single seed selection | 01_LineBreeding/01_PhenotypicSelection/02_SingleSeedDescent/ |
| 3 | - Pedigree selection | 01_LineBreeding/01_PhenotypicSelection/03_PedigreeSelection/ |
| 4 | - Doubled-haploid | 01_LineBreeding/01_PhenotypicSelection/04_DoubledHaploid/ |
| 5 | - Genomic selection | 01_LineBreeding/02_GenomicSelection/ |
| 6 | - Two-part genomic selection | 01_LineBreeding/03_TwoPartGS/ |
|  | Clonal breeding | 02_ClonalBreeding/ |
| 7 | - Phenotypic selection | 02_ClonalBreeding/01_PhenotypicSelection/ |
| 8 | - Pedigree selection | 02_ClonalBreeding/02_PedigreeSelection/ |
| 9 | - Genomic selection | 02_ClonalBreeding/03_GenomicSelection/ |
|  | Hybrid breeding | 03_HybridBreeding/ |
| 10 | - Phenotypic selection | 03_HybridBreeding/01_PhenotypicSelection/ |
| 11 | - Genomic selection | 03_HybridBreeding/02_GenomicSelection/ |
| 12 | - Two-part genomic selection | 03_HybridBreeding/03_TwoPartGS/ |

**Table A2.** List of breeding techniques presented in this paper and their file paths.

| Technique | Description | File Path |
| --- | --- | --- |
| 1 | Trait introgression | 04_Techniques/traitIntrogression.R |
| 2 | Speed breeding | 04_Techniques/speedBreeding.R |
| 3 | Genome editing | 04_Techniques/genomeEditing.R |
| 4 | Multiple traits | 04_Techniques/multipleTraits.R |
| 5 | Miscellaneous slot | 04_Techniques/miscellaneousSlot.R |
| 6 | Mating plans | 04_Techniques/matingPlans.R |
| 7 | GWAS | 04_Techniques/simulateGWAS.R |
| 8 | Genomic models | 04_Techniques/genomicModels.R |
| 9 | Set heritability | 04_Techniques/setHeritability.R |
| 10 | Import external haplotypes | 04_Techniques/importExternalHaplo.R |
| 11 | Specify demography | 04_Techniques/specifyDemography.R |

