## Supplementary material for "Plant breeding simulations with AlphaSimR": Scripts: AdvanceYear.R

```
# Advance year

# Advance breeding program by 1 year
# Works backwards through pipeline to avoid copying data

# Stage 3
# Release variety

# Stage 2
F1 <- setPheno(F1, varE = varE)
selected <- selectInd(F1,nParents)
output$accSel[year] <- cor(F1@gv,F1@pheno)

# Stage 1
F1 <- randCross(Parents, nCrosses)
```
