## Supplementary material for "Plant breeding simulations with AlphaSimR": Scripts: CreateParents.R

```
# Create founders

# Generate initial haplotypes
founderPop = runMacs(nInd     = nParents,
                     segSites = nQtl + nSnp,
                     inbred   = TRUE,
                     species  = "WHEAT")
SP = SimParam$new(founderPop)

# Add SNP chip
SP$restrSegSites(nQtl, nSnp)
if (nSnp > 0) {
  SP$addSnpChip(nSnp)
}

# Add traits: trait represents yield
SP$addTraitAG(nQtlPerChr = nQtl,
              mean       = initMeanG,
              var        = initVarG)

# Collect pedigree
SP$setTrackPed(TRUE)

# Create founder parents
Parents = newPop(founderPop)

rm(founderPop)
```
