## Supplementary material for "Plant breeding simulations with AlphaSimR": Scripts: ExtraFunctions.R

```
# This function is used to calculate between family accuracies
#' @param pop an object of Pop-class
accuracy_family <- function(pop){
  mother <- pop@mother
  father <- pop@father
  df <- data.frame(
    mother = mother,
    father = father
  )
  df <- unique(df)
  families <- vector(mode = "list", length = nrow(df))
  for (i in 1:nrow(df)) {
    mother_i <- df$mother[i]
    father_i <- df$father[i]
    families_i <- pop@mother == mother_i & pop@father == father_i
    tmp <- pop[families_i]
    families[[i]] <- tmp
    # PG: Do we want group level heritability so there is measurement error?
  }
  phenotypes <- unlist(lapply(families, meanP))
  gvs <- unlist(lapply(families,meanG))
  # PG: sometimes the gvs are all identical throwing an NA for the correlation.
  cor(gvs,phenotypes)
}
```
