## Supplementary material for "Plant breeding simulations with AlphaSimR": Scripts: FillPipeline.R

```
# Fill breeding pipeline

# Set initial yield trials with unique individuals
for(cohort in 1:2){
  cat("  FillPipeline stage:",cohort,"of 2\n")
  if(cohort < 3){
    #Stage 1
    F1 <- randCross(Parents, nCrosses)
  }
  if(cohort < 2){
    #Stage 2
    F1 <- setPheno(F1, varE = varE)
    selected <- selectInd(F1,nParents)
  }
  if(cohort < 1){
    ##Stage 3
    #Release variety
  }
}
```
