## Supplementary material for "Plant breeding simulations with AlphaSimR": Scripts: GlobalParameters.R

```
# Global Parameters

# ---- Number of simulation replications and breeding cycles ----

nReps   = 1 # Number of simulation replicates
nBurnin = 20 # Number of years in burnin phase
nFuture = 20 # Number of years in future phase
nCycles = nBurnin + nFuture

# ---- Genome simulation ----

nQtl = 1000 # Number of QTL per chromosome
nSnp = 0    # Number of SNP per chromosome

# ---- Initial parents mean and variance ----

initMeanG  = 1
initVarG   = 1
varE = 1

# ---- Breeding program details ----

nParents = 50  # Number of parents to start a breeding cycle
nCrosses = 100 # Number of crosses per year
```
