## Supplementary material for "Plant breeding simulations with AlphaSimR": Scripts: AdvanceYear.R

```
# Advance year

# Advance breeding program by 1 year
# Works backwards through pipeline to avoid copying data

# Stage 11
# Release variety

# Stage 10
EYT = selectInd(AYT, nEYT)
EYT = setPheno(EYT, varE = varE, reps = repEYT)

# Stage 9
AYT = selectInd(PYT, nAYT)
AYT = setPheno(AYT, varE = varE, reps = repAYT)

# Stage 8
output$accSel[year] = cor(HDRW@gv, HDRW@pheno)
PYT = selectWithinFam(HDRW, famMax)
PYT = selectInd(PYT, nPYT)
PYT = setPheno(PYT, varE = varE, reps = repPYT)

# Stage 7
HDRW = setPheno(S4, varE = varE, reps = repHDRW)

# Stage 6
# Assume all seed germinates
# Take single seed from each plant
# Lines almost fully inbred
S4 = self(S3)

# Stage 5
# Assume all seed germinates
# Take single seed from each plant
S3 = self(S2)

# Stage 4
# Assume all seed germinates
# Take single seed from each plant
S2 = self(S1)

# Stage 3
# Take single seed from each plant
S1 = self(S0)

# Stage 2
# Create segregating population
S0 = self(F1, nProgeny)

# Stage 1
F1 = randCross(Parents, nCrosses)
```
