## Supplementary material for "Plant breeding simulations with AlphaSimR": Scripts: CreateParents.R

```
# Create founders

# Generate initial haplotypes
founderPop = runMacs(nInd     = nParents,
                     segSites = nQtl + nSnp,
                     inbred   = TRUE,
                     species  = "WHEAT")
SP = SimParam$new(founderPop)

# Add SNP chip
SP$restrSegSites(nQtl, nSnp)
if (nSnp > 0) {
  SP$addSnpChip(nSnp)
}

# Add traits: trait represents yield
SP$addTraitAG(nQtlPerChr = nQtl,
              mean       = initMeanG,
              var        = initVarG,
              varEnv     = initVarEnv,
              varGxE     = initVarGE)

# Collect pedigree to conduct the Single-Hill method
SP$setTrackPed(TRUE)

# Create founder parents
Parents = newPop(founderPop)

# Add phenotype reflecting evaluation in EYT
Parents = setPheno(Parents, varE = varE, reps = repEYT)
rm(founderPop)
```
