## Supplementary material for "Plant breeding simulations with AlphaSimR": Scripts: FillPipeline.R

```
# Fill breeding pipeline

# Set initial yield trials with unique individuals
for(cohort in 1:11){
  cat("  FillPipeline stage:",cohort,"of 11\n")
  if(cohort < 11){
    # Stage 1
    F1 = randCross(Parents, nCrosses)
  }
  if(cohort < 10){
    # Stage 2
    S0 = self(F1, nProgeny)
  }
  if(cohort < 9){
    # Stage 3
    S1 = self(S0)
  }
  if(cohort < 8){
    # Stage 4
    S2 = self(S1)
  }
  if(cohort < 7){
    # Stage 5
    S3 = self(S2)
  }
  if(cohort < 6){
    # Stage 6
    S4 = self(S3)
  }
  if(cohort < 5){
    # Stage 7
    HDRW = setPheno(S4, varE = varE, reps = repHDRW)
  }
  if(cohort < 4){
    # Stage 8
    PYT = selectWithinFam(HDRW, famMax)
    PYT = selectInd(PYT, nPYT)
    PYT = setPheno(PYT, varE = varE, reps = repPYT)
  }
  if(cohort < 3){
    # Stage 9
    AYT = selectInd(PYT, nAYT)
    AYT = setPheno(AYT, varE = varE, reps = repAYT)
  }
  if(cohort < 2){
    # Stage 10
    EYT = selectInd(AYT, nEYT)
    EYT = setPheno(EYT, varE = varE, reps = repEYT)
  }
  if(cohort < 1){
    # Stage 11
    # Release variety
  }
}
```
