## Supplementary material for "Plant breeding simulations with AlphaSimR": Scripts: GlobalParameters.R

```
# Global Parameters

# ---- Number of simulation replications and breeding cycles ----
nReps   = 1 # Number of simulation replicates
nBurnin = 20 # Number of years in burnin phase
nFuture = 20 # Number of years in future phase
nCycles = nBurnin + nFuture

# ---- Genome simulation ----
nQtl = 1000 # Number of QTL per chromosome
nSnp = 0    # Number of SNP per chromosome

# ---- Initial parents mean and variance ----
initMeanG  = 1
initVarG   = 1
initVarEnv = 1e-6 # Virtually zero for consistency with 2-Part paper
initVarGE  = 2
varE       = 4 # Yield trial error variance, bushels per acre
               # Relates to error variance for an entry mean

# ---- Breeding program details ----
nParents = 50  # Number of parents to start a breeding cycle
nCrosses = 100 # Number of crosses per year
nProgeny = 10  # Selfed pregeny produced per cross
famMax   = 10  # The maximum number of selfed lines per cross to enter PYT
nPYT     = 500 # Entries per preliminary yield trial
nAYT     = 50  # Entries per advanced yield trial
nEYT     = 10  # Entries per elite yield trial

# Effective replication of yield trials
repHDRW  = 4/9 #h2 = 0.1
repPYT   = 1   #h2 = 0.2
repAYT   = 4   #h2 = 0.5
repEYT   = 8   #h2 = 0.7
```
