## Supplementary material for "Plant breeding simulations with AlphaSimR": Scripts: 00RUNME.R

```
# Script name: Phenotypic line breeding program with pedigree selection
#
# Authors: Jon Bancic, Philip Greenspoon, Chris Gaynor, Gregor Gorjanc
#
# Date Created: 2023-01-23

# ---- Clean environment and load packages ----
rm(list = ls())
# install.packages(pkgs = "AlphaSimR")
library(package = "AlphaSimR")
source(file = "ExtraFunctions.R")

# ---- Load global parameters ----
source(file = "GlobalParameters.R")
scenarioName = "LinePheno_pedigree"

# ---- Create list to store results from reps ----
results = list()

for(REP in 1:nReps) {
  cat("Working on REP:", REP,"\n")

  # ---- Create a data frame to track key parameters ----
  output = data.frame(year     = 1:nCycles,
                      rep      = rep(REP, nCycles),
                      scenario = rep(scenarioName, nCycles),
                      meanG    = numeric(nCycles),
                      varG     = numeric(nCycles),
                      accSel   = numeric(nCycles))

  # ---- Create initial parents ----
  source(file = "CreateParents.R")

  # ---- Fill breeding pipeline with unique individuals from initial parents ----
  source(file = "FillPipeline.R")

  # ---- Burn-in phase ----
  for(year in 1:nBurnin) {
    cat("  Working on burn-in year:",year,"\n")
    source(file = "UpdateParents.R") # Pick parents
    source(file = "AdvanceYear.R")   # Advances yield trials by a year
    # Report results
    output$meanG[year] = meanG(mergePops(F2))
    output$varG[year]  = varG(mergePops(F2))
  }

  # ---- Future phase ----
  for(year in (nBurnin+1):(nBurnin+nFuture)) {
    cat("  Working on future year:",year,"\n")
    source(file = "UpdateParents.R") # Pick parents
    source(file = "AdvanceYear.R")   # Advances yield trials by a year
    ## Report results
    output$meanG[year] = meanG(mergePops(F2))
    output$varG[year]  = varG(mergePops(F2))
  }

  ## Save results from current replicate
  results = append(results, list(output))
}

# Save results
saveRDS(results, file = paste0(scenarioName,".rds"))
```
