## Supplementary material for "Plant breeding simulations with AlphaSimR": Scripts: AdvanceYear.R

```
# Advance year

# Advance breeding program by 1 year
# Works backwards through pipeline to avoid copying data

# Stage 10
# Release variety

# Stage 9
EYT = selectInd(AYT, nEYT)
EYT = setPheno(EYT, varE = varE, reps = repEYT)

# Stage 8
AYT = selectInd(PYT, nAYT)
AYT = setPheno(AYT, varE = varE, reps = repAYT)

# Stage 7
output$accSel[year] = accuracy_family(F6)
PYT = selectInd(F6, nPYT)
PYT = setPheno(PYT, varE = varE, reps = repPYT)

# Stage 6
F6 = vector("list",nCrosses) #Selected plants from each cross
for(i in 1:nCrosses){ # Loop over crosses
  n = nInd(F5[[i]]) # Number of rows per cross
  F6lines = vector("list",n) # Rows in crosses
  F6pheno = numeric(n) # Row phenotypes
  for(j in 1:n){
    F6lines[[j]] = F5[[i]][j] # No selfing due to deriving lines
    F6_j = self(F5[[i]][j],plantsPerRow)
    F6pheno[j] = meanP(F6_j)
  }
  #  Select "nRowF6" F6 rows per cross
  take = order(F6pheno,decreasing=TRUE)[1:nRowF6]
  F6lines = F6lines[take]
  # Derive new lines from rows
  F6[[i]] = mergePops(F6lines)
}
F6 = mergePops(F6)
F6 = setPheno(F6, varE = varE, reps = repF6)

# Stage 5
#  Grow selected plants in F5 rows
F5 = vector("list",nCrosses) # Selected plants from each cross
for(i in 1:nCrosses){ # Loop over crosses
  n = nInd(F4[[i]]) # Number of rows per cross
  F5rows = vector("list",n) # Rows in crosses
  F5pheno = numeric(n) # Row phenotypes
  for(j in 1:n){
    F5rows[[j]] = self(F4[[i]][j],plantsPerRow)
    F5pheno[j] = meanP(F5rows[[j]])
  }
  # Select "nSelF5" F5 rows per cross
  take = order(F5pheno,decreasing=TRUE)[1:nRowF5]
  F5rows = F5rows[take]
  #  Select "nSelF5" plants per F5 row
  for(j in 1:nRowF5){
    F5rows[[j]] = setPheno(F5rows[[j]], varE = varE, reps = 1)
    F5rows[[j]] = selectInd(F5rows[[j]],nSelF5)
  }
  F5[[i]] = mergePops(F5rows)
}

# Stage 4
#  Grow selected plants in F4 rows
F4 = vector("list",nCrosses) # Selected plants from each cross
for(i in 1:nCrosses){ # Loop over crosses
  n = nInd(F3[[i]]) # Number of rows per cross
  F4rows = vector("list",n) # Rows in crosses
  F4pheno = numeric(n) # Row phenotypes
  for(j in 1:n){
    F4rows[[j]] = self(F3[[i]][j],plantsPerRow)
    F4pheno[j] = meanP(F4rows[[j]])
  }
  # Select "nRowF4" F4 rows per cross
  take = order(F4pheno,decreasing=TRUE)[1:nRowF4]
  F4rows = F4rows[take]
  # Select "nSelF4" plants per F4 row
  for(j in 1:nRowF4){
    F4rows[[j]] = setPheno(F4rows[[j]], varE = varE, reps = 1)
    F4rows[[j]] = selectInd(F4rows[[j]],nSelF4)
  }
  F4[[i]] = mergePops(F4rows)
}

# Stage 3
F3 = vector("list",nCrosses) # Selected plants from each cross
for(i in 1:nCrosses){ # Loop over crosses
  n = nInd(F2[[i]]) # Number of rows per cross
  F3rows = vector("list",n) # Rows in crosses
  F3pheno = numeric(n) # Row phenotypes
  for(j in 1:n){
    F3rows[[j]] = self(F2[[i]][j],plantsPerRow)
    F3pheno[j] = meanP(F3rows[[j]])
  }
  # Select "nRowF3" F3 rows per cross
  take = order(F3pheno,decreasing=TRUE)[1:nRowF3]
  F3rows = F3rows[take]
  # Select "nSelF3" plants per selected F3 row
  for(j in 1:nRowF3){
    F3rows[[j]] = setPheno(F3rows[[j]], varE = varE, reps = 1)
    F3rows[[j]] = selectInd(F3rows[[j]],nSelF3)
  }
  F3[[i]] = mergePops(F3rows)
}

# Stage 2
F2 = vector("list",nCrosses) # Keep crosses seperate
for(i in 1:nCrosses){ # Loop over crosses
  F2_i = self(F1[i], nProgeny = nF2)
  F2_i = setPheno(F2_i, varE = varE, reps = 1)
  F2[[i]] = selectInd(F2_i, nInd = nSelF2)
}

# Stage 1
F1 = randCross(Parents, nCrosses)
```
