## Supplementary material for "Plant breeding simulations with AlphaSimR": Scripts: GlobalParameters.R

```
# Global Parameters

# ---- Number of simulation replications and breeding cycles ----
nReps   = 1   # Number of simulation replicates
nBurnin = 20  # Number of years in burnin phase
nFuture = 20  # Number of years in future phase
nCycles = nBurnin + nFuture

# ---- Genome simulation ----
nQtl = 1000 # Number of QTL per chromosome
nSnp = 0    # Number of SNP per chromosome
nChromosomes = 1 #Number of chromosomes

# ---- Initial parents mean and variance ----
initMeanG  = 1
initVarG   = 1
initVarEnv = 1e-6 # Virtually zero for consistency with 2-Part paper
initVarGE  = 2
varE       = 4 # Yield trial error variance, bushels per acre
               # Relates to error variance for an entry mean
## h2 = 0.05

# ---- Breeding program details ----
nParents = 20 # Number of parents to start a breeding cycle
nCrosses = 40 # Number of crosses per year

# Number of progeny per selfed individual in each stage
nF2 = 100
plantsPerRow = 20

# Number of individuals to select in each stage
nPYT = 100 # Entries per preliminary yield trial
nAYT = 50  # Entries per advanced yield trial
nEYT = 10  # Entries per elite yield trial
nSelF2 = 10
nSelF3 = 4
nSelF4 = 4
nSelF5 = 4

# Number of rows to select in each stage
nRowF3 = 10
nRowF4 = 10
nRowF5 = 10
nRowF6 = 4

# Effective replication of yield trials
repF6  = 4/9 # h2 = 0.1
repPYT = 1   #h2 = 0.2
repAYT = 4   #h2 = 0.5
repEYT = 8   #h2 = 0.7
```
