## Supplementary material for "Plant breeding simulations with AlphaSimR": Scripts: 00RUNME.R

```
# Script name: Genomic selection wheat line breeding program
#
# Authors: Initially developed by Chris Gaynor; exanded/polished for this
# publication by Jon Bancic, Philip Greenspoon, Chris Gaynor, Gregor Gorjanc
#
# Date Created: 2023-01-23
#
# Applies GS to advance individuals from DH to make PYT as well as to select new
# parents from DH stage.

# ---- Clean environment and load packages ----
rm(list = ls())
# install.packages(pkgs = "AlphaSimR")
library(package = "AlphaSimR")

# ---- Load global parameters ----
source(file = "GlobalParameters.R")
scenarioName = "LineGS"

# ---- Create list to store results from reps ----
results = list()

for(REP in 1:nReps){
  cat("Working on REP:", REP,"\n")

  # ---- Create a data frame to track key parameters ----
  output = data.frame(year     = 1:nCycles,
                      rep      = rep(REP, nCycles),
                      scenario = "",
                      meanG = numeric(nCycles),
                      varG  = numeric(nCycles),
                      acc_sel = numeric(nCycles))
  # ---- Create initial parents ----
  source(file = "CreateParents.R")

  # ---- Fill breeding pipeline with unique individuals from initial parents ----
  source(file = "FillPipeline.R")

  # ---- Simulate year effects ----
  P = runif(nCycles)

  # ---- Burn-in phase: Phenotypic selection program ----
  cat("--> Working on Burn-in \n")
  for(year in 1:nBurnin) {
    cat(" Working on burn-in year:",year,"\n")
    source(file = "UpdateParents.R") # Pick new parents
    source(file = "AdvanceYear.R")   # Advance yield trials by a year
    source(file = "StoreTrainPop.R") # Store training population
    # Report results
    output$meanG[year] = meanG(DH)
    output$varG[year]  = varG(DH)
  }

  # ---- Future phase: Genomic selection program ----
  cat("--> Working on Genomic line breeding program \n")
  for(year in (nBurnin+1):(nBurnin+nFuture)) {
    cat(" Working on future year:",year,"\n")
    source(file = "RunGSModels.R")      # Run genomic model
    source(file = "UpdateParents_GS.R") # Pick new parents
    source(file = "AdvanceYear_GS.R")   # Advance yield trials by a year
    source(file = "StoreTrainPop.R")    # Store training population
    # Report results
    output$meanG[year] = meanG(DH)
    output$varG[year]  = varG(DH)
  }

  # Save results from current replicate
  results = append(results, list(output))
}

# Save results
saveRDS(results, file = paste0(scenarioName,".rds"))
```
