## Supplementary material for "Plant breeding simulations with AlphaSimR": Scripts: AdvanceYear_GS.R

```
# Advance year

# Advance breeding program by 1 year
# Works backwards through pipeline to avoid copying data

# Stage 6
# Release variety

# Stage 5
EYT = selectInd(AYT, nEYT)
EYT = setPheno(EYT, varE = varE, reps = repEYT)

# Stage 4
AYT = selectInd(PYT, nAYT)
AYT = setPheno(AYT, varE = varE, reps = repAYT)

# Stage 3 - apply genomic selection
# NOTE: HDRW removed because phenotyping not needed
DH = setEBV(DH, gsModel)
output$accSel[year] = cor(DH@gv, DH@ebv)
PYT = selectWithinFam(DH, famMax,use = "ebv")
PYT = selectInd(PYT, nPYT, use="ebv")
PYT = setPheno(PYT, varE = varE, reps = repPYT)

# Stage 2
DH = makeDH(F1, nDH)

# Stage 1
F1 = randCross(Parents, nCrosses)
```
