## Supplementary material for "Plant breeding simulations with AlphaSimR": Scripts: StoreTrainPop.R

```
# Store training population

if (year == startTP){
  cat("  Start collecting training population \n")
  TrainPop = c(PYT, EYT, AYT)
}

if (year > startTP & year < nBurnin+1){
  cat("  Collecting training population \n")
  TrainPop = c(TrainPop,
               PYT, EYT, AYT)
}

if (year > nBurnin){
  cat("  Maintaining training population \n")
  TrainPop = c(TrainPop[-c(1:c(PYT, EYT, AYT)@nInd)],
               PYT, EYT, AYT)
}
```
