## Supplementary material for "Plant breeding simulations with AlphaSimR": Scripts: UpdateParents_GS.R

```
# Update parents

# Replace 10 oldest inbred parents with 10 new genomically-predicted
# inbreds from DH stage

# Predict ebv of DHs
DH = setEBV(DH, gsModel)

# Select 10 new parents based on EBVs
newParents = selectInd(DH, 10, use = "ebv")

# Replace 10 oldest inbred parents with 10 new inbreds from DH stage
Parents   = c(Parents[11:nParents], newParents)
```
