## Supplementary material for "Plant breeding simulations with AlphaSimR": Scripts: 00RUNME.R

```
# Script name: Two-part wheat line breeding program
#
# Authors: Initially developed by Chris Gaynor; exanded/polished for this
# publication by Jon Bancic, Philip Greenspoon, Chris Gaynor, Gregor Gorjanc
#
# Date Created: 2023-01-23
#
# Uses two-part strategy with rapid cycling of parents in population improvement
# and conventional breeding for product development. Applies GS to advance
# individuals from DH to make PYT as well as in population improvement.

# ---- Clean environment and load packages ----
rm(list = ls())
# install.packages(pkgs = "AlphaSimR")
library(package = "AlphaSimR")

# ---- Load global parameters ----
source(file = "GlobalParameters.R")
scenarioName = "LineGSTP"

# ---- Create list to store results from reps ----
results = list()
results_accPI = list()

for(REP in 1:nReps){
  cat("Working on REP:", REP,"\n")

  # ---- Create a data frame to track key parameters ----
  output = data.frame(year     = 1:nCycles,
                      rep      = rep(REP, nCycles),
                      scenario = "",
                      meanG = numeric(nCycles),
                      varG  = numeric(nCycles),
                      accSel  = numeric(nCycles))

  # ---- Create initial parents ----
  source(file = "CreateParents.R")

  # ---- Fill breeding pipeline with unique individuals from initial parents ----
  source(file = "FillPipeline.R")

  # ---- Simulate year effects ----
  P = runif(nCycles)

  # ---- Burn-in phase: Phenotypic selection program ----
  cat("--> Working on Burn-in \n")
  for(year in 1:nBurnin) {
    cat(" Working on burn-in year:",year,"\n")
    source(file = "UpdateParents.R") # Pick new parents
    source(file = "AdvanceYear.R")   # Advance yield trials by a year
    source(file = "StoreTrainPop.R") # Store training population
    # Report results
    output$meanG[year] = meanG(DH)
    output$varG[year]  = varG(DH)
  }

  # ---- Future phase: Genomic selection program ----
  cat("--> Working on Two-part line breeding program \n")
  # New parameters for population improvement
  nCyclesPI = 2    # Number of rapid cycles per year
  nParents  = 50   # Number of parents
  nCrossPI  = 100  # Number of crosses per cycle
  nF1PI = 100      # Number of F1-PI to advance to PD
  # Create a data frame to track selection accuracy in PI
  accPI = data.frame(accPI = numeric(nFuture*nCyclesPI))

  for(year in (nBurnin+1):(nBurnin+nFuture)) {
    cat(" Working on future year:",year,"\n")
    source(file = "RunGSModels.R")      # Run genomic model
    source(file = "AdvanceYear_GSTP.R")   # Advance yield trials by a year
    source(file = "StoreTrainPop.R")    # Store training population
    # Report results
    output$meanG[year] = meanG(DH)
    output$varG[year]  = varG(DH)
  }

  # Save results from current replicate
  results = append(results, list(output))
  results_accPI = append(results_accPI, list(accPI))
}

# Save results
saveRDS(results, file = paste0(scenarioName,".rds"))
saveRDS(results_accPI, file = paste0(scenarioName,"_accPI.rds"))
```
