## Supplementary material for "Plant breeding simulations with AlphaSimR": Scripts: AdvanceYear_GSTP.R

```
# Advance year

# Advance breeding program by 1 year
# Works backwards through pipeline to avoid copying data

# Stage 6
# Release variety

# Stage 5
EYT = selectInd(AYT, nEYT)
EYT = setPheno(EYT, varE = varE, reps = repEYT)

# Stage 4
AYT = selectInd(PYT, nAYT)
AYT = setPheno(AYT, varE = varE, reps = repAYT)

# Stage 3 - apply genomic selection
# NOTE: HDRW removed because phenotyping not needed
DH = setEBV(DH, gsModel)
output$accSel[year] = cor(DH@gv, DH@ebv)
PYT = selectWithinFam(DH, famMax,use = "ebv")
PYT = selectInd(PYT, nPYT, use="ebv")
PYT = setPheno(PYT, varE = varE, reps = repPYT)

# Stage 2
DH = makeDH(F1, nDH)

# Stage 1
# Run population improvement
count = ifelse((year == nBurnin+1), 1, count + nCyclesPI)
for(cycle in 1:nCyclesPI){
  cat("   Population improvement cycle", cycle, "/", nCyclesPI,"\n")
  if(cycle == 1){
    if (year == (nBurnin + 1)) {
      # Create F1s by crossing parents from Burn-in
      Parents = randCross(Parents, nCrossPI)
    }
    # 1. Select best F1s using GS
    Parents = setEBV(Parents, gsModel)
    # Report selection accuracy
    accPI$accPI[count] = cor(Parents@gv, Parents@ebv)
    # F1s to advance to product development
    F1 = selectInd(Parents, nF1PI, use = "ebv")
    # F1s to advance to next cycle as new parents
    Parents = selectInd(Parents, nParents, use = "ebv")

    # 2. Make parental crosses
    Parents = randCross(Parents, nCrossPI)
  } else {
    # 1. Select best F1s using GS
    Parents = setEBV(Parents, gsModel)
    # Report selection accuracy
    accPI$accPI[count+cycle-1] = cor(Parents@gv, Parents@ebv)
    # F1s to advance to next cycle as new parents
    Parents = selectInd(Parents, nParents, use = "ebv")

    # 2. Make parental crosses
    Parents = randCross(Parents, nCrossPI)
  }
}
```
