## Supplementary material for "Plant breeding simulations with AlphaSimR": Scripts: ANALYZERESULTS.R

```
# install.packages(pkgs = "dplyr")
library(package = "dplyr")

# Read in results
df <- bind_rows(readRDS("LineGSTP.rds"))
df2 <- bind_rows(readRDS("LineGSTP_accPI.rds"))

# Plot results
png("Results.png", height = 800, width = 300)
par(mfrow=c(4,1))

# Genetic Gain
plot(-19:20,rowMeans(matrix(df$meanG,ncol = max(df$rep))),type="l",
     main="Genetic gain",xlab="Year",ylab="Yield")

# Variance
plot(-19:20,rowMeans(matrix(df$varG,ncol = max(df$rep))),type="l",
     main="Genetic variance",xlab="Year",ylab="Variance")

# Selection accuracy
plot(-19:20,rowMeans(matrix(df$accSel,ncol = max(df$rep))),type="l",
     main="Selection accuracy in Product Development",xlab="Year",ylab="Correlation")

# Selection accuracy in population improvement
plot(1:40,rowMeans(matrix(df2$accPI,ncol = max(df$rep))),type="l",
     main="Selection accuracy in Population Improvement",xlab="Year",ylab="Correlation",xaxt = "n")
axis(1, at = seq(0,40,10), labels = seq(0,20,5))
abline(v = seq(1,41,2),col="gray80",lty=2)

dev.off()
```
