## Supplementary material for "Plant breeding simulations with AlphaSimR": Scripts: PaperPlot.R

```
# This script summarizes results from RUNME_all.R script
# and produces Figure 6 in the manuscript.

# ---- Clean environment and load packages ----

rm(list = ls())
# install.packages(pkgs = c("dplyr", "readr", "reshape2", "ggplot2", "ggpubr"))
library(package = "dplyr")
library(package = "readr")
library(package = "reshape2")
library(package = "ggplot2")
library(package = "ggpubr")

# ---- Specify ggplot theme options ----
theme_optns <- theme_bw(base_size = 16, base_family = "sans") +
  theme(panel.background = element_blank(),
        # legend.title = element_blank(),
        legend.title = element_text(size = 14, face = "bold"),
        legend.text  = element_text(size = 14),
        legend.position = "right",
        plot.title = element_text(size = 15, face = "bold"),
        axis.text  = element_text(size = 14,colour = "black"),
        axis.title = element_text(size = 18),
        # axis.title.x = element_blank(),
        axis.text.x  = element_text(hjust = .9, angle = 45),
        strip.text   = element_text(face = "bold", size = 18, colour = 'black'))


# ---- Read in and organise results ----

nReps = 10
df.pheno <- read_csv("Results_LinePheno.csv")
df.GS    <- read_csv("Results_LineGS_const.csv")
df.GSun  <- read_csv("Results_LineGS_unconst.csv")
temp <- rbind(df.pheno,df.GS,df.GSun)
rm(df.pheno,df.GS,df.GSun)

# Organise data
colnames(temp)[c(1,3)] <- c("Year","Scenario")
temp <- melt(temp,
             id.vars = c("Year", "rep", "Scenario"),
             measure.vars = c("meanG","varG","accSel"))


# Rename Scenarios
# temp[temp$Year < 21,]$Scenario = "Burn-in"
temp$Scenario <- recode_factor(temp$Scenario,
                               `Burn-in`        = "Pheno burn-in",
                               `LinePheno`      = "Pheno",
                               `LineGS_const`   = "GS-constrained",
                               `LineGS_unconst` = "GS-unconstrained")
temp0 <- temp # save for later t-tests

# ---- Plot results ----

#### Plot genetic gain of each replicate in the burnin phase ####

temp$paste <- factor(paste0(temp$rep,temp$Scenario))
# Create a mean line
df.mean <- temp %>%
  group_by(Year, Scenario, variable) %>%
  summarise(value = mean(value)) %>%
  filter((variable %in% c("meanG"))) %>%
  droplevels
df.mean$paste <- factor(paste0(df.mean$Scenario))
(a1 <- temp %>%
    filter((variable %in% c("meanG"))) %>%
    filter(Year < 21) %>%
    droplevels() %>% 
    ggplot(aes(x = Year, y = value, group = paste)) +
    geom_line(aes(color = Scenario), linewidth = 0.6, alpha = 0.2) +
    geom_line(data = df.mean[df.mean$Year < 22,], aes(x = Year, y = value), linewidth = 0.8) +
    # ylim(-0.1,5) +
    xlim(0,40) +
    # ggtitle("Burn-in phase") +
    ylab("Genetic gain") + 
    theme_optns +
    guides(color = guide_legend(override.aes = list(linewidth = 1, alpha = 1))))

#### Plot genetic gain of each replicate in the future phase ####

# First center data to year 20
mean_burnin20 <- df.mean$value[df.mean$Year == 20]
for(i in 1:nReps){
  mean = temp$value[temp$rep == i & temp$Year == 20 & temp$variable == "meanG"]
  center = temp$value[temp$rep == i & temp$variable == "meanG"] - mean
  temp$value[temp$rep == i & temp$variable == "meanG"] = center + mean_burnin20
}
# Plot centered future genetic gain for separate reps
df.center <- temp %>% 
  filter((variable %in% c("meanG"))) %>%
  filter(Year > 20) %>%
  droplevels()
(a2 <- a1 + 
    geom_line(data = df.center, 
              aes(x = Year, y = value, color = Scenario), linewidth = 0.6, alpha = 0.2) +
    geom_line(data = df.mean[df.mean$Year > 20,], aes(colour = paste), linewidth = 0.8)) 

##-- Calculate means for variance and accuracy 
temp <- temp %>%
  group_by(Year, Scenario, variable) %>%
  summarise(value = mean(value)) %>%
  droplevels
# Black line for burnin
temp2 <- temp %>%
  filter((Scenario %in% c("Pheno"))) %>%
  filter(Year < 21) %>%
  droplevels()

# Genetic variance
(b <- temp %>%
    filter((variable %in% c("varG"))) %>%
    droplevels() %>% 
    ggplot(aes(x = Year, y = value)) +
    geom_line(aes(color = Scenario), linewidth = 0.8) +
    geom_line(data = temp2[temp2$variable == "varG",],
              aes(x = Year, y = value), color = "black", linewidth = 0.8) +
    ylab("Genetic variance") + 
    theme_optns)

# Selection accuracy
(c <- temp %>%
    filter((variable %in% c("accSel"))) %>%
    droplevels() %>% 
    ggplot(aes(x = Year, y = value)) +
    geom_line(aes(color = Scenario), linewidth = 0.8) +
    geom_line(data = temp2[temp2$variable == "accSel",],
              aes(x = Year, y = value), color = "black", linewidth = 0.8) +
    ylab("Accuracy") + 
    theme_optns) 

#### Merge plots and save ####

(p1 <- ggarrange(a2, nrow = 1, ncol = 1, 
                 common.legend = T, legend = "none", align = "hv"))
(p2 <- ggarrange(b, c, nrow = 2, ncol = 1, 
                 common.legend = T, legend = "right", align = "hv"))
(p <- ggarrange(p1, p2, nrow = 1, ncol = 2, 
                common.legend = T, widths = c(1,1)))
ggsave(plot = p, filename ="06_Figure.png", width = 4.5, height = 2.5, scale = 2.5)


# ---- Pair-wise comparison test ----

# Comparison constrained GS to Pheno
t.test(temp0[with(temp0,Scenario == "Pheno" & Year == 40 & variable == "meanG"),]$value,
       temp0[with(temp0,Scenario == "GS-constrained" & Year == 40 & variable == "meanG"),]$value,
       paired = T)[3] # 0.03960644

# Comparison unconstrained GS to Pheno
t.test(temp0[with(temp0,Scenario == "Pheno" & Year == 40 & variable == "meanG"),]$value,
       temp0[with(temp0,Scenario == "GS-unconstrained" & Year == 40 & variable == "meanG"),]$value,
       paired = T)[3] # 0.0001225286

# Comparison constrained GS to unconstrained GS
t.test(temp0[with(temp0,Scenario == "GS-constrained" & Year == 40 & variable == "meanG"),]$value,
       temp0[with(temp0,Scenario == "GS-unconstrained" & Year == 40 & variable == "meanG"),]$value,
       paired = T)[3] # 0.7701437
```
