## Supplementary material for "Plant breeding simulations with AlphaSimR": Scripts: RUNME_all.R

```
# Script name: Run wheat breeding programs
#
# Authors: Jon Bancic, Philip Greenspoon, Chris Gaynor, Gregor Gorjanc
#
# Date Created: 2023-01-23
#
# This script runs both phenotypic and genomic selection program one after
# another.

# ---- Clean environment and load packages ----
rm(list = ls())
# install.packages(pkgs = "AlphaSimR")
library(package = "AlphaSimR")

# ---- Number of simulation replications ----
nReps = 10

for(REP in 1:nReps){
  cat("Working on REP:", REP,"\n")

  # ---- Load global parameters ----
  source(file = "02_GenomicSelection/GlobalParameters.R")

  # ---- Create initial parents ----
  source(file = "02_GenomicSelection/CreateParents.R")

  # ---- Fill breeding pipeline with unique individuals from initial parents ----
  source(file = "02_GenomicSelection/FillPipeline.R")

  # ---- Create a data frame to track key parameters----
  output = data.frame(year     = 1:nCycles,
                      rep      = rep(REP, nCycles),
                      scenario = "",
                      meanG  = numeric(nCycles),
                      varG   = numeric(nCycles),
                      accSel = numeric(nCycles))

  # ---- Burn-in phase: Phenotypic selection program ----
  # NOTE: some scripts below are for phenoypic selection, but saved in genomic
  # selection folder!
  cat("--> Working on Burn-in with Phenotypic selection program \n")
  for(year in 1:nBurnin) {
    cat(" Working on burn-in year:",year,"\n")
    source(file = "02_GenomicSelection/UpdateParents.R") # Pick new parents
    source(file = "02_GenomicSelection/AdvanceYear.R")   # Advance yield trials by a year
    source(file = "02_GenomicSelection/StoreTrainPop.R") # Store training population
    # Report results
    output$meanG[year] = meanG(DH)
    output$varG[year]  = varG(DH)
  }
  # Save burn-in to load later for subsequent scenarios
  save.image("tmp.RData")

  # ---- Future phase: Phenotypic selection program ----
  cat("--> Working on Phenotypic selection program \n")
  for(year in (nBurnin+1):(nBurnin+nFuture)) {
    cat(" Working on future year:",year,"\n")
    source(file = "02_GenomicSelection/UpdateParents.R") # Pick new parents
    source(file = "02_GenomicSelection/AdvanceYear.R")   # Advance yield trials by a year
    source(file = "02_GenomicSelection/StoreTrainPop.R") # Store training population
    # Report results
    output$meanG[year] = meanG(DH)
    output$varG[year]  = varG(DH)
  }
  # Save results
  cat(" Saving results \n")
  Scenario  <- output$scenario <- "LinePheno"
  file.name <- paste0("Results_",Scenario,".csv")
  write.table(output, file.name, sep = ",",
              col.names = !file.exists(file.name), row.names = F, append = T)

  # ---- Future phase: Genomic selection program with unconstrained costs ----
  cat("--> Working on cost-unconstrained Genomic selection program \n")
  load("tmp.RData")
  for(year in (nBurnin+1):(nBurnin+nFuture)) {
    cat(" Working on future year:",year,"\n")
    source(file = "02_GenomicSelection/RunGSModels.R")      # Run genomic model
    source(file = "02_GenomicSelection/UpdateParents_GS.R") # Pick new parents
    source(file = "02_GenomicSelection/AdvanceYear_GS.R")   # Advance yield trials by a year
    source(file = "02_GenomicSelection/StoreTrainPop.R")    # Store training population
    # Report results
    output$meanG[year] = meanG(DH)
    output$varG[year]  = varG(DH)
  }
  # Save results
  cat(" Saving results \n")
  Scenario  <- output$scenario <- "LineGS_unconst"
  file.name <- paste0("Results_",Scenario,".csv")
  write.table(output, file.name, sep = ",",
              col.names = !file.exists(file.name), row.names = F, append = T)

  # ---- Future phase: Genomic selection program with constrained costs ----
  cat("--> Working on cost-constrained Genomic selection program \n")
  load("tmp.RData")
  # Reduce number of DHs produced per cross
  # to offset costs of genotyping
  nDH <- 75

  for(year in (nBurnin+1):(nBurnin+nFuture)) {
    cat(" Working on future year:",year,"\n")
    source(file = "02_GenomicSelection/RunGSModels.R")      # Run genomic model
    source(file = "02_GenomicSelection/UpdateParents_GS.R") # Pick new parents
    source(file = "02_GenomicSelection/AdvanceYear_GS.R")   # Advance yield trials by a year
    source(file = "02_GenomicSelection/StoreTrainPop.R")    # Store training population
    # Report results
    output$meanG[year] = meanG(DH)
    output$varG[year]  = varG(DH)
  }
  # Save results
  cat(" Saving results \n")
  Scenario  <- output$scenario <- "LineGS_const"
  file.name <- paste0("Results_",Scenario,".csv")
  write.table(output, file.name, sep = ",",
              col.names = !file.exists(file.name), row.names = F, append = T)

  # ---- Future phase: Two-Part genomic selection program ----
  cat("--> Working on Two-part genomic selection program \n")
  load("tmp.RData")

  # New parameters for population improvement
  nCyclesPI = 2    # Number of rapid cycles per year
  nParents  = 50   # Number of parents
  nCrossPI  = 100  # Number of crosses per cycle
  nF1PI = 100      # Number of F1-PI to advance to PD
  # Create a data frame to track selection accuracy in PI
  accPI = data.frame(accPI = numeric(nFuture*nCyclesPI))

  for(year in (nBurnin+1):(nBurnin+nFuture)) {
    cat(" Working on future year:",year,"\n")
    source(file = "03_TwoPartGS/RunGSModels.R")      # Run genomic model
    source(file = "03_TwoPartGS/AdvanceYear_GSTP.R")   # Advance yield trials by a year
    source(file = "03_TwoPartGS/StoreTrainPop.R")    # Store training population
    # Report results
    output$meanG[year] = meanG(DH)
    output$varG[year]  = varG(DH)
  }

  # Save results
  cat(" Saving results \n")
  Scenario  <- output$scenario <- "LineGSTP"
  file.name <- paste0("Results_",Scenario,".csv")
  write.table(output, file.name, sep = ",",
              col.names = !file.exists(file.name), row.names = F, append = T)
}

# Remove temporary file
file.remove("tmp.RData")

# Summarize results using PaperPlot.R script
```
