## Supplementary material for "Plant breeding simulations with AlphaSimR": Scripts: AdvanceYear.R

```
# Advance year

# Advance breeding program by 1 year
# Works backwards through pipeline to avoid copying data

# Stage 16
ECT6 = setPheno(ECT5, varE = VarE, reps = repECT, p = P[year])

# Stage 15
ECT5 = ECT4

# Stage 14
ECT4 = ECT3

# Stage 13
ECT3 = ECT2

# Stage 12
ECT2 = ECT1

# Stage 11
ECT1 = selectInd(ACT5, nInd = nClonesECT, use = "pheno")

# Stage 10
ACT5 = setPheno(ACT4, varE = VarE, reps = repACT, p = P[year])

# Stage 9
ACT4 = ACT3

# Stage 8
ACT3 = ACT2

# Stage 7
ACT2 = ACT1

# Stage 6
output$accSel[year] = cor(gv(HPT3), pheno(HPT3)) # accuracy based on 2000 inds
ACT1 = selectInd(HPT3, nInd = nClonesACT, use = "pheno")

# Stage 5
HPT3 = setPheno(HPT2, varE = VarE, reps = repHPT, p = P[year])

# Stage 4
HPT2 = HPT1

# Stage 3
HPT1 = Seedlings

# Stage 2
Seedlings = setPheno(F1, varE = VarE, reps = repHPT, p = P[year])

# Stage 1
# Crossing block
F1 = randCross(Parents, nCrosses = nCrosses, nProgeny = nProgeny)
```
