## Supplementary material for "Plant breeding simulations with AlphaSimR": Scripts: ANALYZERESULTS.R

```
# install.packages(pkgs = "dplyr")
library(package = "dplyr")

# Read in results
df <- bind_rows(readRDS("ClonalPheno.rds"))

# Plot results
png("Results.png", height = 600, width = 300)
par(mfrow=c(3,1))

# Inbred Genetic Gain
plot(1:80,rowMeans(matrix(df$meanG,ncol = max(df$rep))),type="l",
     main="Genetic gain",xlab="Year",ylab="Yield")

# Inbred Variance
plot(1:80,rowMeans(matrix(df$varG,ncol = max(df$rep))),type="l",
     main="Genetic variance",xlab="Year",ylab="Variance")

# Selection accuracy
plot(1:80,rowMeans(matrix(df$accSel,ncol = max(df$rep))),type="l",
     main="Selection accuracy",xlab="Year",ylab="Correlation")
dev.off()
```
