## Supplementary material for "Plant breeding simulations with AlphaSimR": Scripts: CreateParents.R

```
# Create founders

# Create founder population
founderPop = runMacs2(nInd     = nParents,
                      nChr     = nChr,
                      segSites = nQtl+nSnp,
                      genLen   = genLen,
                      mutRate  = mutRate)

# Set simulation parameters
SP = SimParam$new(founderPop)

# Add SNP chip
SP$restrSegSites(nQtl,nSnp)
if (nSnp > 0) {
  SP$addSnpChip(nSnp)
}

# Add traits: trait represents yield
SP$addTraitADG(nQtlPerChr = nQtl,
               mean       = initMeanG,
               var        = initVarG,
               varGxE     = initVarGE)

# Collect pedigree
SP$setTrackPed(TRUE)

# Create founder parents
Parents = newPop(founderPop)

# Set a phenotype to founder parents
Parents = setPheno(Parents, varE = VarE, reps = repECT)
rm(founderPop)
```
