## Supplementary material for "Plant breeding simulations with AlphaSimR": Scripts: FillPipeline.R

```
# Fill breeding pipeline

# Set initial yield trials with unique individuals

# Sample year effects
P = runif(16)

# Breeding program
for(cohort in 1:16) {
  cat("  FillPipeline stage:",cohort,"of 16\n")
  # Stage 1 Crossing block
  F1 = randCross(Parents, nCrosses = nCrosses, nProgeny = nProgeny)
  if(cohort < 16){
    # Stage 2 Germinate the seedlings in the nursery
    Seedlings = setPheno(F1, varE = VarE, reps = repHPT, p = P[cohort])
  }
  if(cohort < 15){
    # Stage 3 Plant in the seedlings in the field as HPT and record yields
    HPT1 = Seedlings
  }
  if(cohort < 14){
    # Stage 4 Record the HPT yields
    HPT2 = HPT1
  }
  if(cohort < 13){
    # Stage 5 Record the HPT yields
    HPT3 = setPheno(HPT2, varE = VarE, reps = repHPT, p = P[cohort+3L])
  }
  if(cohort < 12){
    # Stage 6 Select 500 superior individuals and plant as advanced clonal trials (ACT)
    ACT1 = selectInd(HPT3, nInd = nClonesACT, use = "pheno")
  }
  if(cohort < 11){
    # Stage 7 Record ACT yields
    ACT2 = ACT1
  }
  if(cohort < 10){
    # Stage 8 Record ACT yields
    ACT3 = ACT2
  }
  if(cohort < 9){
    # Stage 9 Record ACT yields
    ACT4 = ACT3
  }
  if(cohort < 8){
    # Stage 10 Record ACT yields
    ACT5 = setPheno(ACT4, varE = VarE, reps = repACT, p = P[cohort+8L])
  }
  if(cohort < 7){
    # Stage 11 Select 30 superior individuals and plant as elite clonal trials (ECT)
    ECT1 = selectInd(ACT5, nInd = nClonesECT, use = "pheno")
  }
  if(cohort < 6){
    # Stage 12 Record ECT yields
    ECT2 = ECT1
  }
  if(cohort < 5){
    # Stage 13 Record ECT yields
    ECT3 = ECT2
  }
  if(cohort < 4){
    # Stage 14 Record ECT yields
    ECT4 = ECT3
  }
  if(cohort < 3){
    # Stage 15 Record ECT yields
    ECT5 = ECT4
  }
  if(cohort < 2){
    # Stage 16 Record ECT yields
    ECT6 = setPheno(ECT5, varE = VarE, reps = repECT, p = P[cohort+14L])
  }
}
```
