## Supplementary material for "Plant breeding simulations with AlphaSimR": Scripts: GlobalParameters.R

```
# Global Parameters

# ---- Number of simulation replications and breeding cycles ----
nReps   = 1       # Number of simulation replicates
nBurnin = 40      # Number of years in burnin phase
nFuture = 40      # Number of years in future phase
startRecords = 35 # Year when training and pedigree record collecting begins
nCycles = nBurnin + nFuture

# ---- Genome simulation ----
nChr    = 15      # Number of chromosomes
nQtl    = 160     # Number of QTL per chromosome: 15 chr x 160 QTL = 2400 QTLs
nSnp    = 600     # Simulate SNP chip with 9000 markers
genLen  = 1       # Genetic length
PhyLen  = 1e+08   # Physical length
mutRate = 2.5e-08 # Mutation rate

# ---- Initial parents mean and variance ----
initMeanG = 2500    # Phenotypic mean
initVarG  = 150000  # Genetic variance
initVarGE = 150000  # Genotype-by-year interaction variance
VarE      = 2800000 # Single variance

# ---- Breeding program details ----
nParents   = 20   # Number of parents (and founders)
nCrosses   = 100  # Number of crosses
nProgeny   = 20   # Number of progenies per cross
nClonesACT = 500  # Number of individuals selected at ACT stage
nClonesECT = 40   # Number of individuals selected at ECT stage

# Effective replication of yield trials
repHPT     = 1   # h2 = 0.05
repACT     = 15  # h2 = 0.45
repECT     = 50  # h2 = 0.65
```
