## Supplementary material for "Plant breeding simulations with AlphaSimR": Scripts: AdvanceYear_Pedigree.R

```
# Advance year

# Advance breeding program by 1 year
# Works backwards through pipeline to avoid copying data

# Stage 13
ECT6 = setPheno(ECT5, varE = VarE, reps = repECT, p = P[year])

# Stage 12
ECT5 = ECT4

# Stage 11
ECT4 = ECT3

# Stage 10
ECT3 = ECT2

# Stage 9
ECT2 = ECT1

# Stage 8
ECT1 = selectInd(ACT5, nInd = nClonesECT, use = "pheno")

# Stage 7
ACT5 = setPheno(ACT4, varE = VarE, reps = repACT, p = P[year])

# Stage 6
ACT4 = ACT3

# Stage 5
ACT3 = ACT2

# Stage 4
ACT2 = ACT1

# Stage 3
# Use pedigree estimated breeding values to select seedlings for further evaluation
Seedlings@ebv       = as.matrix(tail(EBV, Seedlings@nInd))
output$accSel[year] = cor(gv(Seedlings), ebv(Seedlings))
ACT1 = selectInd(Seedlings, nInd = nClonesACT, use = "ebv")

# Stage 2
Seedlings = setPheno(F1, varE = VarE, reps = repHPT, p = P[year])

# Stage 1
# Crossing block
F1 = randCross(Parents, nCrosses = nCrosses, nProgeny = nProgeny)
```
