## Supplementary material for "Plant breeding simulations with AlphaSimR": Scripts: RunModel_Pedigree.R

```
# Run pedigree BLUP model using ASReml

# Pedigree BLUP is used to predict breeding values of Seedlings in
# order to skip HPT stages

# Prepare prediction dataset for seedlings
ped_temp = rbind(Pedigree,
                 data.frame(Ind   = c(Seedlings@id),
                            Sire  = c(Seedlings@father),
                            Dam   = c(Seedlings@mother),
                            Year  = year,
                            Stage = rep("Seedlings",Seedlings@nInd),
                            Pheno = NA))

# Construct numerator relationship matrix A
ped_temp$Ind  = as.factor(ped_temp$Ind)
ped_temp$Dam  = as.factor(ped_temp$Dam)
ped_temp$Sire = as.factor(ped_temp$Sire)
A = ainverse(ped_temp[,1:3])

# Run model
ped_temp$Year[is.na(ped_temp$Year)] = 0
ped_temp$Year  = as.factor(ped_temp$Year)
ped_temp$Stage = as.factor(ped_temp$Stage)

asreml.options(trace=FALSE)
pedModel <- asreml(fixed = Pheno ~ 1 + Year,
                   random = ~ vm(Ind, A),
                   # residual = ~ dsum(~id(units) | Year),
                   residual = ~ units,
                   na.action = na.method(y='include'),
                   data = ped_temp)

# Loop to ensure model converges
while (pedModel$converge != TRUE) {
  pedModel <- update.asreml(pedModel)
}
rm(ped_temp)

# Assign estimated breeding values to Seedling
EBV = pedModel$coef$random
```
