## Supplementary material for "Plant breeding simulations with AlphaSimR": Scripts: StoreTrainPop.R

```
# Save pedigree records and training population for GS from year 35

ACT5@fixEff <- as.integer(rep(year,nInd(ACT5)))
ECT6@fixEff <- as.integer(rep(year,nInd(ECT6)))

if(year == startRecords) {
  trainPop = ECT6
  Pedigree = data.frame(Ind   = c(ACT5@id),
                        Sire  = c(ACT5@father),
                        Dam   = c(ACT5@mother),
                        Year  = year,
                        Stage = c(rep("ACT5",ACT5@nInd)),
                        Pheno = c(ACT5@pheno))
}

if (year > startRecords & year < nBurnin+1) {
  trainPop = c(trainPop,ECT6)
  Pedigree = rbind(Pedigree,
                   data.frame(Ind   = c(ACT5@id),
                              Sire  = c(ACT5@father),
                              Dam   = c(ACT5@mother),
                              Year  = year,
                              Stage = c(rep("ACT5",ACT5@nInd)),
                              Pheno = c(ACT5@pheno)))
}

if (year > nBurnin) {
  # Update training population (set to keep 6 years worth of records)
  remove   = table(trainPop@fixEff)[1]            # remove oldest records
  trainPop = c(trainPop[-c(1:remove)], ECT6)      # add new records

  # Update pedigree with new records from ACT stage
  Pedigree = rbind(Pedigree,
                   data.frame(Ind   = c(ACT5@id),
                              Sire  = c(ACT5@father),
                              Dam   = c(ACT5@mother),
                              Year  = year,
                              Stage = c(rep("ACT",ACT5@nInd)),
                              Pheno = c(ACT5@pheno)))
}
```
