## Supplementary material for "Plant breeding simulations with AlphaSimR": Scripts: 00RUNME.R

```
# Script name: Phenotypic selection hybrid maize breeding program
#
# Authors: Initially developed by Chris Gaynor; exanded/polished for this
# publication by Jon Bancic, Philip Greenspoon, Chris Gaynor, Gregor Gorjanc
#
# Date Created: 2023-01-23

# ---- Clean environment and load packages ----
rm(list = ls())
# install.packages(pkgs = "AlphaSimR")
library(package = "AlphaSimR")

# ---- Load global parameters ----
source(file = "GlobalParameters.R")
scenarioName = "HybridPheno"

# ---- Create list to store results from reps ----
results = list()

for(REP in 1:nReps){
  cat("Working on REP:", REP,"\n")

  # ---- Create a data frame to track key parameters ----
  output = data.frame(year     = 1:nCycles,
                      rep      = rep(REP, nCycles),
                      scenario = scenarioName,
                      meanG_inbred = numeric(nCycles),
                      varG_inbred  = numeric(nCycles),
                      meanG_hybrid = numeric(nCycles),
                      varG_hybrid  = numeric(nCycles),
                      acc_sel = numeric(nCycles),
                      cor = numeric(nCycles))

  # ---- Create initial parents and set testers ----
  source(file = "CreateParents.R")

  # ---- Fill breeding pipeline with unique individuals from initial parents ----
  source(file = "FillPipeline.R")

  # ---- Simulate year effects ----
  P = runif(nCycles)

  # ---- Burn-in phase ----
  cat("--> Working on Burn-in \n")
  for(year in 1:nBurnin) {
    cat(" Working on burnin year:",year,"\n")
    source(file = "UpdateParents.R") # Pick new parents
    source(file = "UpdateTesters.R") # Pick new testers
    source(file = "AdvanceYear.R")   # Advance yield trials by a year
    source(file = "StoreTrainPop.R") # Store training population
    # Report results
    output$meanG_inbred[year] = (meanG(MaleInbredYT3) + meanG(FemaleInbredYT3))/2
    output$varG_inbred[year]  = (varG(MaleInbredYT3) + varG(FemaleInbredYT3))/2
    tmp = hybridCross(FemaleInbredYT3, MaleInbredYT3, returnHybridPop=TRUE)
    output$meanG_hybrid[year] = meanG(tmp)
    output$varG_hybrid[year]  = varG(tmp)
    tmp = calcGCA(tmp,use="gv")
    output$cor[year] = cor(c(tmp$GCAf[,2],tmp$GCAm[,2]),
                           c(FemaleInbredYT3@gv[,1],MaleInbredYT3@gv[,1]))
  }

  # ---- Future phase: Phenotypic program ----
  cat("--> Working on Phenotypic hybrid program \n")
  output$scenario <- "Pheno"
  for(year in (nBurnin+1):(nBurnin+nFuture)) {
    cat(" Working on future year:",year,"\n")
    source(file = "UpdateParents.R") # Pick new parents
    source(file = "UpdateTesters.R") # Pick new testers
    source(file = "AdvanceYear.R")   # Advance yield trials by a year
    # Report results
    output$meanG_inbred[year] = (meanG(MaleInbredYT3) + meanG(FemaleInbredYT3))/2
    output$varG_inbred[year]  = (varG(MaleInbredYT3) + varG(FemaleInbredYT3))/2
    tmp = hybridCross(FemaleInbredYT3, MaleInbredYT3, returnHybridPop=TRUE)
    output$meanG_hybrid[year] = meanG(tmp)
    output$varG_hybrid[year]  = varG(tmp)
    tmp = calcGCA(tmp,use="gv")
    output$cor[year] = cor(c(tmp$GCAf[,2],tmp$GCAm[,2]),
                           c(FemaleInbredYT3@gv[,1],MaleInbredYT3@gv[,1]))
  }

  # Save results from current replicate
  results = append(results, list(output))
}

# Save results
saveRDS(results, file = paste0(scenarioName,".rds"))
```
