## Supplementary material for "Plant breeding simulations with AlphaSimR": Scripts: ANALYZERESULTS.R

```
# install.packages(pkgs = "dplyr")
library(package = "dplyr")

# Read in results
df <- bind_rows(readRDS("HybridPheno.rds"))

# Plot results
png("Results.png", height = 800, width = 600)
par(mfrow=c(3,2))

# Inbred Genetic Gain
plot(-19:20,rowMeans(matrix(df$meanG_inbred,ncol = max(df$rep))),type="l",
     main="Inbred genetic gain",xlab="Year",ylab="Yield")

# Hybrid Genetic Gain
plot(-19:20,rowMeans(matrix(df$meanG_hybrid,ncol = max(df$rep))),type="l",
     main="Hybrid genetic gain",xlab="Year",ylab="Yield")

# Inbred Variance
plot(-19:20,rowMeans(matrix(df$varG_inbred,ncol = max(df$rep))),type="l",
     main="Inbred genetic variance",xlab="Year",ylab="Variance")

# Hybrid Variance
plot(-19:20,rowMeans(matrix(df$varG_hybrid,ncol = max(df$rep))),type="l",
     main="Hybrid genetic variance",xlab="Year",ylab="Correlation")

# Selection accuracy
plot(-19:20,rowMeans(matrix(df$acc_sel,ncol = max(df$rep))),type="l",
     main="Selection accuracy",xlab="Year",ylab="Correlation")

# Correlation
plot(-19:20,rowMeans(matrix(df$cor,ncol = max(df$rep))),type="l",
     main="Inbred vs. hybrid yield cor.",xlab="Year",ylab="Variance")
dev.off()
```
