## Supplementary material for "Plant breeding simulations with AlphaSimR": Scripts: CreateParents.R

```
# Create founders

cat("Making initial parents \n")

# Create founder population
founderPop = runMacs(nInd = nParents*2,
                     nChr = nChr,
                     segSites = nQtl+nSnp,
                     inbred = TRUE,
                     split = nGenSplit,
                     species = "MAIZE")

# Set simulation parameters
SP = SimParam$new(founderPop)

# Add SNP chip
SP$restrSegSites(nQtl,nSnp)
if (nSnp > 0) {
  SP$addSnpChip(nSnp)
}

# Add traits: trait represents yield
SP$addTraitADG(nQtlPerChr = nQtl,
               mean   = initMeanG,
               var    = initVarG,
               meanDD = MeanDD,
               varDD  = VarDD,
               varGxE = initVarGE)
# Set permanent yield trial error variance
SP$setVarE(varE = VarE)

# Split heterotic pools to form initial parents
FemaleParents = newPop(founderPop[1:nParents])
MaleParents   = newPop(founderPop[(nParents+1):(nParents*2)])

# Set hybrid parents for later yield trials
MaleElite   = selectInd(MaleParents, nElite, use = "gv")
FemaleElite = selectInd(FemaleParents, nElite, use = "gv")

# Reverse order to keep best parent in longer
MaleElite = MaleElite[nElite:1]
FemaleElite = FemaleElite[nElite:1]

# Set initial testers for YT1 and YT2
# Requires nTesters to be smaller than nElite
MaleTester1   = MaleElite[1:nTester1]
FemaleTester1 = FemaleElite[1:nTester1]
MaleTester2   = MaleElite[1:nTester2]
FemaleTester2 = FemaleElite[1:nTester2]
rm(founderPop)
```
