## Supplementary material for "Plant breeding simulations with AlphaSimR": Scripts: FillPipeline.R

```
# Fill breeding pipeline

#Set initial yield trials with unique individuals

P = runif(6) # p-values for GxY effect

for(cohort in 1:6){
  cat("FillPipeline year:", cohort, "of 6\n")

  # Stage 1
  MaleF1   = randCross(MaleParents, nCrosses)
  FemaleF1 = randCross(FemaleParents, nCrosses)

  # Stage 2
  if(cohort<6){
    p = P[6-cohort]

    MaleDH   = makeDH(MaleF1, nDH)
    FemaleDH = makeDH(FemaleF1, nDH)

    MaleYT1   = setPhenoGCA(MaleDH, FemaleTester1,
                            reps = repYT1, inbred = T, p = p)
    FemaleYT1 = setPhenoGCA(FemaleDH, MaleTester1,
                            reps = repYT1, inbred = T, p = p)
  }

  # Stage 3
  if(cohort<5){
    p = P[5-cohort]

    MaleYT2   = selectInd(MaleYT1, nInbred2)
    FemaleYT2 = selectInd(FemaleYT1, nInbred2)

    MaleYT2   = setPhenoGCA(MaleYT2, FemaleTester2,
                            reps = repYT2, inbred = T, p = p)
    FemaleYT2 = setPhenoGCA(FemaleYT2, MaleTester2,
                            reps = repYT2, inbred = T, p = p)
  }

  # Stage 4
  if(cohort<4){
    p = P[4-cohort]

    MaleInbredYT3   = selectInd(MaleYT2, nInbred3)
    FemaleInbredYT3 = selectInd(FemaleYT2, nInbred3)

    MaleHybridYT3   = hybridCross(MaleInbredYT3, FemaleElite)
    FemaleHybridYT3 = hybridCross(FemaleInbredYT3, MaleElite)

    MaleHybridYT3   = setPheno(MaleHybridYT3,
                               reps = repYT3, p = p)
    FemaleHybridYT3 = setPheno(FemaleHybridYT3,
                               reps = repYT3, p = p)
  }

  # Stage 5
  if(cohort<3){
    p = P[3-cohort]

    MaleHybridYT4   = selectInd(MaleHybridYT3, nYT4)
    FemaleHybridYT4 = selectInd(FemaleHybridYT3, nYT4)

    MaleHybridYT4   = setPheno(MaleHybridYT4,
                               reps = repYT4, p = p)
    FemaleHybridYT4 = setPheno(FemaleHybridYT4,
                               reps = repYT4, p = p)

    MaleInbredYT4 =
      MaleInbredYT3[MaleInbredYT3@id%in%MaleHybridYT4@mother]
    FemaleInbredYT4 =
      FemaleInbredYT3[FemaleInbredYT3@id%in%FemaleHybridYT4@mother]
  }

  # Stage 6
  if(cohort<2){
    p = P[2-cohort]

    MaleHybridYT5   = selectInd(MaleHybridYT4, nYT5)
    FemaleHybridYT5 = selectInd(FemaleHybridYT4, nYT5)

    MaleHybridYT5   = setPheno(MaleHybridYT5,
                               reps = repYT5, p = p)
    FemaleHybridYT5 = setPheno(FemaleHybridYT5,
                               reps = repYT5, p = p)

    MaleInbredYT5 =
      MaleInbredYT4[MaleInbredYT4@id%in%MaleHybridYT5@mother]
    FemaleInbredYT5 =
      FemaleInbredYT4[FemaleInbredYT4@id%in%FemaleHybridYT5@mother]
  }

  # Stage 7, release
}
```
