## Supplementary material for "Plant breeding simulations with AlphaSimR": Scripts: GlobalParameters.R

```
# Global Parameters

# ---- Number of simulation replications and breeding cycles ----
nReps   = 1    # Number of simulation replicates
nBurnin = 20   # Number of years in burnin phase
nFuture = 20   # Number of years in future phase
nCycles = nBurnin + nFuture
startTP = 16   # Year to start training population

# ---- Genome simulation ----
nChr = 15       # Number of chromosomes
nQtl = 300      # Number of QTL per chromosome
nSnp = 400      # Number of SNP per chromosome
nGenSplit = 100 # Heterotic pool split

# ---- Initial inbred parents mean and variance ----
initMeanG = 70 # bushels per acre
initVarG  = 20 # bushels per acre
# Degree of dominance
MeanDD = 0.92   # mean
VarDD  = 0.3    # variance
# Error variances
initVarGE = 40   # Genotype-by-year interaction
VarE = 270  # Yield trial error variance, bushels per acre
            # Relates to error variance for an entry mean

# ---- Breeding program details ----
nParents = 50  # Number of parents to start a breeding cycle
nCrosses = 80  # Number of crosses per year
famMax   = 15  # The maximum number of DH lines per cross
nDH      = 50  # DH lines produced per cross

# Effective replication of yield trials
repYT1 = 1    #h2 = 0.06
repYT2 = 2    #h2 = 0.11
repYT3 = 4    #h2 = 0.20
repYT4 = 8    #h2 = 0.34
repYT5 = 100  #h2 = 0.86

# --- Selection on GCA ----
# Number of inbreds per heterotic pool per stage
nInbred1 = nCrosses*nDH # Do not change
nInbred2 = 400
nInbred3 = 40

# Number of testers per heterotic pool per stage
# Values must be smaller than nElite
nTester1 = 1
nTester2 = 3

# Yield trial entries
nYT1 = nInbred1*nTester1 # Do not change
nYT2 = nInbred2*nTester2 # Do not change

# ---- Selection on SCA ----

# Elite parents per heterotic pool
nElite = 5

# Elite YT size
nYT3 = nInbred3*nElite # Do not change
nYT4 = 20
nYT5 = 4
```
