## Supplementary material for "Plant breeding simulations with AlphaSimR": Scripts: StoreTrainPop.R

```
# Store training population of the past 3 years

if (year == startTP){
  cat("  Start collecting training population \n")
  MaleTrainPop = c(MaleInbredYT3,MaleInbredYT4,MaleInbredYT5)
  FemaleTrainPop = c(FemaleInbredYT3,FemaleInbredYT4,FemaleInbredYT5)
}

if (year > startTP & year < nBurnin+1){
  cat("  Collecting training population \n")
  MaleTrainPop = c(MaleTrainPop,
                   MaleInbredYT3,MaleInbredYT4,MaleInbredYT5)
  FemaleTrainPop = c(FemaleTrainPop,
                     FemaleInbredYT3,FemaleInbredYT4,FemaleInbredYT5)
}

if (year > nBurnin){
  cat("  Maintaining training population \n")
  MaleTrainPop = c(MaleTrainPop[-c(1:c(MaleInbredYT3,MaleInbredYT4,MaleInbredYT5)@nInd)],
                   MaleInbredYT3,MaleInbredYT4,MaleInbredYT5)
  FemaleTrainPop = c(FemaleTrainPop[-c(1:c(FemaleInbredYT3,FemaleInbredYT4,FemaleInbredYT5)@nInd)],
                     FemaleInbredYT3,FemaleInbredYT4,FemaleInbredYT5)
}
```
