## Supplementary material for "Plant breeding simulations with AlphaSimR": Scripts: UpdateParents.R

```
# Update parents

# Replace 10 oldest inbred parents with 10 new inbreds from YT4 stage
MaleParents   = c(MaleParents[11:nParents], selectInd(MaleInbredYT4,10))
FemaleParents = c(FemaleParents[11:nParents], selectInd(FemaleInbredYT4,10))
```
