## Supplementary material for "Plant breeding simulations with AlphaSimR": Scripts: UpdateResults.R

```
# Track performance of parental per se performance
inbredMean[year] = (meanG(MaleInbredYT3)+meanG(FemaleInbredYT3))/2
inbredVar[year] = (varG(MaleInbredYT3)+varG(FemaleInbredYT3))/2

# Track performance of parental hybrid performance
tmp = hybridCross(FemaleInbredYT3,MaleInbredYT3,
                  returnHybridPop=TRUE) #Only use with DH parents
hybridMean[year] = meanG(tmp)
hybridVar[year] = varG(tmp)

# Track per se-GCA correlation
tmp = calcGCA(tmp,use="gv")
hybridCorr[year] = cor(c(tmp$GCAf[,2],tmp$GCAm[,2]),
                       c(FemaleInbredYT3@gv[,1],MaleInbredYT3@gv[,1]))
rm(tmp)
```
