## Supplementary material for "Plant breeding simulations with AlphaSimR": Scripts: UpdateTesters.R

```
# Update testers

# Replace oldest hybrid parent with parent of best hybrid from YT6
bestMaleInbred = MaleHybridYT5@mother[
  order(MaleHybridYT5@pheno[,1],decreasing=TRUE)[1]
  ]
MaleElite = c(MaleElite[-1],MaleInbredYT5[bestMaleInbred])

bestFemaleInbred = FemaleHybridYT5@mother[
  order(FemaleHybridYT5@pheno,decreasing=TRUE)[1]
  ]
FemaleElite = c(FemaleElite[-1],FemaleInbredYT5[bestFemaleInbred])

# Update testers
MaleTester1 = MaleElite[1:nTester1]
FemaleTester1 = FemaleElite[1:nTester1]

MaleTester2 = MaleElite[1:nTester2]
FemaleTester2 = FemaleElite[1:nTester2]
```
