## Supplementary material for "Plant breeding simulations with AlphaSimR": Scripts: AdvanceYear_GS.R

```
# Advance year

cat("  Advancing year \n")
# Advance breeding program by 1 year
# Works backwards through pipeline to avoid copying data

# Stage 7
# Release hybrid

# Stage 6
MaleHybridYT5 = selectInd(MaleHybridYT4, nYT5)
FemaleHybridYT5 = selectInd(FemaleHybridYT4, nYT5)

MaleHybridYT5 = setPheno(MaleHybridYT5, reps = repYT5, p = p)
FemaleHybridYT5 = setPheno(FemaleHybridYT5, reps = repYT5, p = p)

MaleInbredYT5 =
  MaleInbredYT4[MaleInbredYT4@id%in%MaleHybridYT5@mother]
FemaleInbredYT5 =
  FemaleInbredYT4[FemaleInbredYT4@id%in%FemaleHybridYT5@mother]

# Stage 5
MaleHybridYT4 = selectInd(MaleHybridYT3, nYT4)
FemaleHybridYT4 = selectInd(FemaleHybridYT3, nYT4)

MaleHybridYT4 = setPheno(MaleHybridYT4, reps = repYT4, p = p)
FemaleHybridYT4 = setPheno(FemaleHybridYT4, reps = repYT4, p = p)

MaleInbredYT4 =
  MaleInbredYT3[MaleInbredYT3@id%in%MaleHybridYT4@mother]
FemaleInbredYT4 =
  FemaleInbredYT3[FemaleInbredYT3@id%in%FemaleHybridYT4@mother]

# Stage 4
MaleInbredYT3 = selectInd(MaleYT2, nInbred3)
FemaleInbredYT3 = selectInd(FemaleYT2, nInbred3)

MaleHybridYT3 = hybridCross(MaleInbredYT3, FemaleElite)
FemaleHybridYT3 = hybridCross(FemaleInbredYT3, MaleElite)

MaleHybridYT3 = setPheno(MaleHybridYT3, reps = repYT3, p = p)
FemaleHybridYT3 = setPheno(FemaleHybridYT3, reps = repYT3, p = p)

# Stage 3 - apply genomic selection
# Predict GCA
if (exists("gsModel")) {
  MaleYT1 = setEBV(MaleYT1, gsModel)
  FemaleYT1 = setEBV(FemaleYT1, gsModel)
} else {
  MaleYT1 = setEBV(MaleYT1, gsModelM)
  FemaleYT1 = setEBV(FemaleYT1, gsModelF)
}

# Report selection accuracy
output$acc_sel[year]  = c((cor(MaleYT1@ebv,MaleYT1@gv) +
                             cor(FemaleYT1@ebv,FemaleYT1@gv))/2)
# Select using EBVs
MaleYT2 = selectInd(MaleYT1, nInbred2, use = "ebv")
FemaleYT2 = selectInd(FemaleYT1, nInbred2, use = "ebv")

MaleYT2 = setPhenoGCA(MaleYT2, FemaleTester2, reps = repYT2, inbred = T, p = p)
FemaleYT2 = setPhenoGCA(FemaleYT2, MaleTester2, reps = repYT2, inbred = T, p = p)

# Stage 2 - apply genomic selection
MaleDH = makeDH(MaleF1, nDH)
FemaleDH = makeDH(FemaleF1, nDH)

# Predict GCA of DHs
if (exists("gsModel")) {
  MaleDH = setEBV(MaleDH, gsModel)
  FemaleDH = setEBV(FemaleDH, gsModel)
} else {
  MaleDH = setEBV(MaleDH, gsModelM)
  FemaleDH = setEBV(FemaleDH, gsModelF)
}
# Select using EBVs
MaleDH = selectInd(selectWithinFam(MaleDH, famMax, use = "ebv"), nInbred2, use = "ebv")
FemaleDH = selectInd(selectWithinFam(FemaleDH, famMax, use = "ebv"), nInbred2, use = "ebv")

MaleYT1 = setPhenoGCA(MaleDH, FemaleTester1, reps = repYT1, inbred = T, p = p)
FemaleYT1 = setPhenoGCA(FemaleDH, MaleTester1, reps = repYT1, inbred = T, p = p)

# Stage 1
MaleF1 = randCross(MaleParents, nCrosses)
FemaleF1 = randCross(FemaleParents, nCrosses)
```
