## Supplementary material for "Plant breeding simulations with AlphaSimR": Scripts: RunGSModels.R

```
# Run genomic models

# # Option 1: Single pool additive GCA model (in code)
# cat("  Running GS model\n")
# gsModelM = RRBLUP(MaleTrainPop)
# gsModelF = RRBLUP(FemaleTrainPop)

# Option 2: Hybrid additive GCA model (in code)
cat("  Running GS model\n")
gsModel = RRBLUP(HybTrainPop)

# Option 3: Hybrid Additive + Dominance Model (GCA + SCA)
# NOTE: Not implemented in the code!
# Example for asigning EBVs
# MaleParents = setEBV(MaleParents, gsModel, value = "bv", targetPop = FemaleParents)
# FemaleParent = setEBV(FemaleParent, gsModel, value = "bv", targetPop = MaleParents)

# Option 4: Hybrid Pool Specific Additive Model (GCAm + GCAf)
# NOTE: Not implemented in the code!
# gsModel = RRBLUP_GCA(HybTrainPop)
# Example for asigning EBVs
# MaleParents = setEBV(MaleParents, gsModel, value = "male")
# FemaleParent = setEBV(FemaleParent, gsModel, value = "female")

# Option 5: Hybrid Pool Specific Additive + Dominance Model (GCAm + GCAf + SCA)
# NOTE: Not implemented in the code!
# gsModel = RRBLUP_SCA(HybTrainPop)
# Example for asigning EBVs
# MaleParents = setEBV(MaleParents, gsModel, value = "male", targetPop = FemaleParents)
# FemaleParent = setEBV(FemaleParent, gsModel, value = "female", targetPop = MaleParents)
```
