## Supplementary material for "Plant breeding simulations with AlphaSimR": Scripts: UpdateParents_GS.R

```
# Update parents

# Replace 10 oldest inbred parents with 10 new genomically-predicted
# inbreds from DH stage

# Predict GCA of DHs
if (exists("gsModel")) {
  MaleDH = setEBV(MaleDH, gsModel)
  FemaleDH = setEBV(FemaleDH, gsModel)
} else {
  MaleDH = setEBV(MaleDH, gsModelM)
  FemaleDH = setEBV(FemaleDH, gsModelF)
}

# Select 10 new parents based on EBVs
newMaleParents = selectInd(MaleDH, 10, use = "ebv")
newFemaleParents = selectInd(FemaleDH, 10, use = "ebv")

# Replace 10 oldest inbred parents with 10 new inbreds from DH stage
MaleParents   = c(MaleParents[11:nParents], newMaleParents)
FemaleParents = c(FemaleParents[11:nParents], newFemaleParents)
```
