## Supplementary material for "Plant breeding simulations with AlphaSimR": Scripts: functions.R

```
# This function calculates Fst among two AlphaSimR populations
calcFst <- function(pop1, pop2) {
  # Pop 1 expected heterozygosity
  M = pullQtlGeno(pop1)
  p1 = colMeans(M)/2
  He_pop1 = mean(2*p1*(1-p1))
  # Pop 2 expected heterozygosity
  M  = pullQtlGeno(pop2)
  p2 = colMeans(M)/2
  He_pop2 = mean(2*p2*(1-p2))
  # Total pop expected heterozygosity
  M  = pullQtlGeno(pop)
  p = colMeans(M)/2
  He_tot = mean(2*p*(1-p))
  # Fst
  Hs = (He_pop1*pop1@nInd+He_pop2*pop2@nInd)/pop@nInd
  return(data.frame("Fst" = (He_tot-Hs)/He_tot))
}

# This function calculates heterozygosity and inbreeding
calcHet <- function(pop) {
  geno = pullQtlGeno(pop)
  Het = mean(rowMeans(1-abs(geno-1)))
  Inb = 1 - Het
  return(data.frame(Het, Inb))
}

# This function creates maximum avoidance mating plan
maxAvoidPlan = function(nInd, nProgeny = 1L){
  crossPlan = matrix(1:nInd, ncol=2, byrow=TRUE)
  tmp = c(seq(1, nInd, by=2),
          seq(2, nInd, by=2))
  crossPlan = cbind(rep(tmp[crossPlan[,1]],
                        each=nProgeny),
                    rep(tmp[crossPlan[,2]],
                        each=nProgeny))
  return(crossPlan)
}

# This function creates circular mating plan
circularPlan = function(nInd, nProgeny = 1) {
  crossPlan = rep(1:nInd, each = 2)
  crossPlan = c(crossPlan[length(crossPlan)], crossPlan[-(length(crossPlan))])
  crossPlan = matrix(crossPlan,
                     ncol = 2, byrow = TRUE)
  crossPlan = crossPlan[rep(1:nrow(crossPlan), each = nProgeny),]
  return(crossPlan)
}

# This function calculates heterosis
calcHeterosis <- function(popA, popB, hybPop) {
  inbMean = (meanG(popA) + meanG(popB))/2
  hybMean = meanG(hybPop)
  heterosis = meanG(hybPop) - (meanG(popA) + meanG(popB))/2
  perHeterosis = (hybMean-inbMean)/inbMean*100
  return(data.frame("Midparent value" = inbMean,
                    "Hybrid value" = hybMean,
                    "Heterosis" = heterosis,
                    "Percent heterosis" = perHeterosis))
}
```
