## Supplementary material for "Plant breeding simulations with AlphaSimR": Scripts: genomeEditing.R

```
# Script name: Genome editing in AlphaSimR
#
# Authors: Jon Bancic, Philip Greenspoon, Chris Gaynor, Gregor Gorjanc
#
# Date Created: 2023-01-23
#
# This script demonstrates editing of a single locus in AlphaSimR.

# ---- Clean environment and load packages ----

rm(list = ls())
# install.packages(pkgs = "AlphaSimR")
library(package = "AlphaSimR")

# ---- Setup simulation ----

# Generate founder haplotypes
founderPop = runMacs(nInd     = 100,
                     segSites = 1000,
                     inbred   = TRUE,
                     species  = "WHEAT")

# Set simulation parameters
SP = SimParam$new(founderPop)

# Create additive trait with 10 QTLs
SP$addTraitAG(nQtlPerChr = 10,
              mean = 1,
              var  = 1)

# Create population
pop = newPop(founderPop)

# Check genetic values
meanG(pop)

# Get QTL haplotypes
genMap = pullQtlHaplo(pop)

# ---- Option 1: Edit single locus in an individual ----

# To change a single locus you can do the following, which changes
# the first homologue in individual 1 to its alternative allele
genMap["1_1",1] = (genMap["1_1", 1] + 1) %% 2

# Or you can manually edit the first QTL on the first homologue to be 1
for(i in 1:pop@nInd) {
  genMap[paste0(i,"_1"), colnames(genMap)[1]] = 1
}
# Re-assign haplotypes
pop2 = setMarkerHaplo(pop, genMap)
# Check genetic mean
meanG(pop2)

# ---- Option 2: Edit single locus in the entire population ----

# Edit QTL with largest additive effect using internal function
pop3 = editGenomeTopQtl(pop, ind = 100, nQtl = 1)
# Check genetic mean
meanG(pop3)
```
