## Supplementary material for "Plant breeding simulations with AlphaSimR": Scripts: genomicModels.R

```
# Script name: Genomic models in AlphaSimR
#
# Authors: Jon Bancic, Philip Greenspoon, Chris Gaynor, Gregor Gorjanc
#
# Date Created: 2023-01-23
#
# This script demonstrates the implementation of different
# genomic models in AlphaSimR. Models are demonstrated 
# using high and low density SNP chip.

# ---- Clean environment and load packages ----

rm(list = ls())
# install.packages(pkgs = "AlphaSimR")
library(package = "AlphaSimR")
source(file = "functions.R")

# ---- Setup simulation ----

# Create founder haplotypes
founderPop = runMacs(
  nInd     = 60,
  segSites = 1000,
  nChr     = 10,
  split    = 100,
  inbred   = TRUE,
  species  = "MAIZE"
)

# Set simulation parameters
SP = SimParam$new(founderPop)

# Ensure QTLs and SNPs do not overlap
SP$restrSegSites(overlap = FALSE)

# Add trait with dominance
SP$addTraitAD(
  nQtlPerChr = 200,
  mean   = 5,
  var    = 1,
  meanDD = 0.92,
  varDD  = 0.2
)

# Add two SNP chips
SP$addSnpChip(nSnpPerChr = 300, name = "LowDensity")
SP$addSnpChip(nSnpPerChr = 800, name = "HighDensity")

# Check that there is no overlap between SNPs and QTLs
snpMap = getSnpMap(snpChip = 1)
qtlMap = getQtlMap(trait = 1)
sum(snpMap$id %in% qtlMap$id)
rm(snpMap,qtlMap)

# Create new population
pop = newPop(founderPop)

# Create two populations (e.g. two heterotic pools)
female = pop[1:30]
male = pop[31:60]

# Create hybrid population
pop = hybridCross(female, male)

# Set phenotypes with a narrow-sense heritability of 0.3
pop = setPheno(pop, h2 = 0.3)

# Store model results
df = data.frame(matrix(NA, 3, 5))
colnames(df) = c("Add_L", "Add_H", "AddDom", "GCA", "SCA")
rownames(df) = c("pop", "female", "male")

# ---- RR-BLUP with additve effects only ----

# Run model with low density chip
gsModel_A1 = RRBLUP(pop, snpChip = 1) # specify chip
# Check accuracy for entire population and
# separate populations
pop = setEBV(pop, gsModel_A1)
df[1, 1] = cor(pop@gv, pop@ebv)
# Females
female = setEBV(female, gsModel_A1)
df[2, 1] = cor(female@gv, female@ebv)
# Males
male = setEBV(male, gsModel_A1)
df[3, 1] = cor(male@gv, male@ebv)

# Run model with high denisty chip
gsModel_A2 = RRBLUP(pop, snpChip = 2) # specify chip
# Check accuracy for entire population and
# separate populations
pop = setEBV(pop, gsModel_A2)
df[1, 2] = cor(pop@gv, pop@ebv)
# Females
female = setEBV(female, gsModel_A2)
df[2, 2] = cor(female@gv, female@ebv)
# Males
male = setEBV(male, gsModel_A2)
df[3, 2] = cor(male@gv, male@ebv)
round(df, 3)

# ---- RR-BLUP with additive and dominance effects ----

# Run model with low density chip
gsModel_AD = RRBLUP_D(pop, snpChip = 1)
# Check accuracy for entire population and
# separate populations
pop = setEBV(pop, gsModel_AD)
df[1, 3] = cor(pop@gv, pop@ebv)
# Females
female = setEBV(female, gsModel_AD)
df[2, 3] = cor(female@gv, female@ebv)
# Males
male = setEBV(male, gsModel_AD)
df[3, 3] = cor(male@gv, male@ebv)
round(df, 3)

# Equivalent to...
addEff <- gsModel_AD@gv[[1]]@addEff
domEff <- gsModel_AD@gv[[1]]@domEff
M <- pullSnpGeno(pop)
ebv <- M %*% addEff + (1-abs(M-1)) %*% domEff
cor(pop@ebv, ebv)

# ---- RR-BLUP with population specific additive (GCA) effects ----

# Run model with low density chip
gsModel_GCA = RRBLUP_GCA(pop, snpChip = 1)
# Check accuracy for entire population and
# separate populations
pop = setEBV(pop, gsModel_GCA)
df[1, 4] = cor(pop@gv, pop@ebv)
# Females
female = setPhenoGCA(female, male, use = "gv", inbred = TRUE)
female = setEBV(female, gsModel_GCA, value = "female")
df[2, 4] = cor(female@gv, female@ebv)
# Males
male = setPhenoGCA(male, female, use = "gv", inbred = TRUE)
male = setEBV(male, gsModel_GCA, value = "male")
df[3, 4] = cor(male@gv, male@ebv)
round(df, 3)

# ---- RR-BLUP with population specific GCA and SCA ----
# GCA - additive effects & SCA - dominance

# Run model with low denisty chip
gsModel_SCA = RRBLUP_SCA(pop, snpChip = 2)
# Check accuracy for entire population and
# separate populations
pop = setEBV(pop, gsModel_SCA)
df[1, 5] = cor(pop@gv, pop@ebv)
# Females
female = setEBV(female, gsModel_SCA,
                value = "female", targetPop = male)
df[2, 5] = cor(female@gv, female@ebv)
# Males
male = setEBV(male, gsModel_SCA,
              value = "male", targetPop = female)
df[3, 5] = cor(male@gv, male@ebv)


# ---- Compare results ----

round(df, 3)
```
