## Supplementary material for "Plant breeding simulations with AlphaSimR": Scripts: importExternalHaplo.R

```
# Script name: Importing External Haplotypes in AlphaSimR
#
# Authors: Initially developed by Chris Gaynor; exanded/polished for this
# publication by Jon Bancic, Philip Greenspoon, Chris Gaynor, Gregor Gorjanc
#
# Date Created: 2023-01-23
#
# This script demonstrates the process of importing external 
# marker data into AlphaSimR for haplotype analysis.

# ---- Clean environment and load packages ----

rm(list = ls())
# install.packages(pkgs = "AlphaSimR")
library(package = "AlphaSimR")

# ---- Step 1: Load in founder haplotypes ----
# This requires a genetic map and phased genotypes

# Load the genetic map in the below format
# Format: Marker name, Chromosome, Position (in Morgans)
# Modeling 10 chromosomes with 10 loci each
# Loci are equally space along 1 Morgan chromosomes
genMap = data.frame(
  marker     = paste0("x", 1:100),
  chromosome = rep(1:10, each = 10),
  position   = rep(seq(from = 0, to = 1, length.out = 10), times = 10)
)

# Load the haplotypes in the below format
# Modeling 10 individuals
# There will need to be 20 rows (# of individuals times 2, since each has 2 haplotypes)
# Each individual will have 100 loci based on the map above
# Generating just random 0s and 1s
haplo = matrix(
  data = sample(0:1, size = 2 * 10 * 100, replace = TRUE),
  nrow = 2 * 10,
  ncol = 100
)

# Assign marker names to the haplotypes
# These can be in any order, because the software will order them based
# on the genetic map automatically
colnames(haplo) = genMap$marker

# Load a pedigree in the below format (optional)
# Pedigree will be for just the 10 individuals represented in the haplotypes
ped = data.frame(id = letters[1:10],
                 mother = rep(0, 10),
                 father = rep(0, 10))

# Create the founder population
# Uses the data structures loaded above
founderPop = importHaplo(
  haplo  = haplo,
  genMap = genMap,
  ploidy = 2,
  ped    = ped
)

# ---- Step 2: Set simulation parameters ----
# Initialize parameters with founder haplotypes

SP = SimParam$new(founderPop)

# Load your own QTL effects (optional)
# This is useful if you want to model known QTL, estimated effects
# from a genomic prediction model, or you just want to create your
# own distribution for effects. The marker names most match the names
# given in the map above. You can model additive effects, dominance
# effects, and an intercept.
qtlEffects = data.frame(marker = c("x1", "x11"),
                        aditiveEffect = c(1, -1))

# Import into SimParam
SP$importTrait(
  markerNames = qtlEffects$marker,
  addEff = qtlEffects$aditiveEffect,
  name = "Your_Trait"
)

# ---- Step 3: Create a population from the founder haplotypes ----

pop = newPop(founderPop)

# The population now works like any other AlphaSimR population
gv(pop)
getPed(pop)
```
