## Supplementary material for "Plant breeding simulations with AlphaSimR": Scripts: matingPlans.R

```
# Script name: Simulation of mating designs in AlphaSimR
#
# Authors: Jon Bancic, Philip Greenspoon, Chris Gaynor, Gregor Gorjanc
#
# Date Created: 2023-01-23
#
# This script provides practical examples of specifying various 
# mating designs within AlphaSimR.

# ---- Clean environment and load packages ----

rm(list = ls())
# install.packages(pkgs = c("AlphaSimR", "ade4", "ggplot2")
library(package = "AlphaSimR")
library(package = "ade4")
library(package = "ggplot2")
source(file = "functions.R")

# ---- Setup simulation ----

founderPop = runMacs(
  nInd     = 100,
  segSites = 1000,
  inbred   = TRUE,
  species  = "MAIZE"
)

# Set simulation parameters
SP = SimParam$new(founderPop)

# Add trait with dominance
SP$addTraitAD(
  nQtlPerChr = 1000,
  mean   = 5,
  var    = 1,
  meanDD = 0.92,
  varDD  = 0.2
)

# Create a single population
pop = newPop(founderPop)

# Set phenotypes with a narrow-sense heritability of 0.5
pop = setPheno(pop, h2 = 0.5)

# Create two populations and testers
popA = pop[1:10]
popB = pop[11:20]
testerA = popA[1:2]
testerB = popB[1:2]

# ---- Crosses within a single population ----

#### Random biparental/paired cross ####
newPop = randCross(pop, nCrosses = 10, nProgeny = 100)
table(newPop@father, newPop@mother) # visualize
# Compare genetic mean and variance
meanG(pop)
meanG(newPop)
varG(pop)
varG(newPop)

#### Full diallel with reciprocal crosses and selfs (n x n) ####
crossPlan = expand.grid(1:nInd(popA), 1:nInd(popA))
newPop = makeCross2(popA, popA, crossPlan, nProgeny = 10)
table(newPop@father, newPop@mother) # visualize
# Compare genetic mean and variance
meanG(pop)
meanG(newPop)
varG(pop)
varG(newPop)

#### Half diallel without reciprocal crosses and selfs (n(n-1)/2) ####
crossPlan = expand.grid(1:5, 6:10)
newPop = makeCross2(popA, popA, crossPlan, nProgeny = 40)
table(newPop@father, newPop@mother) # visualize
# Compare genetic mean and variance
meanG(pop)
meanG(newPop)
varG(pop)
varG(newPop)

#### Specified crosses  (example: Topcross) ####
crossPlan = cbind(1, 2:20)
newPop = makeCross(pop, crossPlan, nProgeny = 50)
table(newPop@father, newPop@mother) # visualize
# Compare genetic mean and variance
meanG(pop)
meanG(newPop)
varG(pop)
varG(newPop)

#### Specified crosses with maximum avoidance ####
crossPlan = maxAvoidPlan(nInd = 20, nProgeny = 100)
newPop = makeCross(pop, crossPlan)
table(newPop@father, newPop@mother) # visualize
# Compare genetic mean and variance
meanG(pop)
meanG(newPop)
varG(pop)
varG(newPop)

#### Develop Multiparent Generation Intercross (MAGIC) population ####
newPop = pop[1:8]
# 2-way crosses
crossPlan = cbind(c(1, 3, 5, 7),
                  c(2, 4, 6, 8))
newPop = makeCross(pop, crossPlan)
table(newPop@father, newPop@mother) # visualize
# 4-way crosses
crossPlan = cbind(c(1, 3),
                  c(2, 4))
newPop = makeCross(newPop, crossPlan)
table(newPop@father, newPop@mother) # visualize
# 8-way crosses
crossPlan = cbind(1, 2)
newPop = makeCross(newPop, crossPlan, nProgeny = 1000)
table(newPop@father, newPop@mother) # visualize
# Create inbred lines from the final cross
for (i in 1:6) {
  newPop = self(newPop, nProgeny = 1)
}
newPop = setPheno(newPop, h2 = 0.5)
# Check inbreeding
1 - mean(rowMeans(1 - abs(pullQtlGeno(pop) - 1)))
# Compare genetic mean and variance
meanG(pop[1:8])
meanG(newPop)
varG(pop[1:8])
varG(newPop)
varP(pop[1:8])
varP(newPop)

# visualize MAGIC with PCA
geno = pullQtlGeno(c(pop[1:8], newPop))
PCA  = dudi.pca(
  df = geno,
  center = T,
  scale = F,
  scannf = FALSE,
  nf = 5
)
(VAF = 100 * PCA$eig[1:5] / sum(PCA$eig)) # variance explained
df.PCA = data.frame(
  "Pop" = c(rep("Parents", 8), rep("MAGIC", 1000)),
  "PC1" = PCA$l1$RS1,
  "PC2" = PCA$l1$RS2
)
# Plot
ggplot(df.PCA, aes(x = PC1, y = PC2)) +
  geom_point(aes(colour = factor(Pop))) +
  ggtitle("Population structure") +
  xlab(paste("Pcomp1: ", round(VAF[1], 2), "%", sep = "")) +
  ylab(paste("Pcomp2: ", round(VAF[2], 2), "%", sep = ""))

# ---- Crosses among two populations ----

#### Testcross/Topcross/North Carolina 1 ####
crossPlan = expand.grid(1:nInd(popA), 1:nInd(testerB))
newPop = makeCross2(popA, popB, crossPlan, nProgeny = 1)
table(newPop@father, newPop@mother) # visualize
# Using internal AlphaSimR function
newPop = hybridCross(females = popA, males =  testerB)
table(newPop@father, newPop@mother) # visualize
# Check heterosis
calcHeterosis(popA, popB, newPop)

#### Factorial/North Carolina 2 (half diallel) ####
crossPlan = expand.grid(1:nInd(popA), 1:nInd(popB))
# with(crossPlan, table(Var1,Var2))
newPop = makeCross2(popA, popB, crossPlan, nProgeny = 1)
table(newPop@father, newPop@mother) # visualize
# Check heterosis
calcHeterosis(popA, popB, newPop)

#### Specified full diallel between two populations ####
crossPlan = expand.grid(1:nInd(popA), 1:nInd(popB))
with(crossPlan, table(Var1, Var2))
newPop1 = makeCross2(popA, popA, crossPlan, nProgeny = 1)
newPop2 = makeCross2(popB, popB, crossPlan, nProgeny = 1)
newPop = c(newPop1, newPop2)
table(newPop@father, newPop@mother) # visualize
# Check heterosis
calcHeterosis(popA, popB, newPop)
```
