## Supplementary material for "Plant breeding simulations with AlphaSimR": Scripts: miscellaneousSlot.R

```
# Script name: Use of miscellaneous slot in AlphaSimR
#
# Authors: Jon Bancic, Philip Greenspoon, Chris Gaynor, Gregor Gorjanc
#
# Date Created: 2023-01-23
#
# This script demonstrates how to use the miscellaneous slot in
# AlphaSimR populations. Here, the slot is designed to be used as part
# of selection criteria.
#
# WARNING: Organisation of the misc slot has changed with AlphaSimR
# version 1.5.4. Prior to this version, misc slot was a list of length
# equal to the number of individuals (nInd). Since version 1.5.4, misc
# slot is a list of length equal to the number of nodes, with each node

# ---- Clean environment and load packages ----

rm(list = ls())
# install.packages(pkgs = "AlphaSimR")
(AlphaSimRVersion = packageVersion(pkg = "AlphaSimR"))
library(package = "AlphaSimR")

# ---- Setup simulation ----

# Create founder haplotypes
founderPop = runMacs(
  nInd     = 10,
  nChr     = 1,
  segSites = 100,
  inbred   = TRUE,
  species  = "GENERIC"
)

# ---- Add two correlated additive traits with same genetic architecture ----

# Set simulation parameters
SP = SimParam$new(founderPop)

# Specify correlation between traits
traitCor = matrix(c(1.0, 0.5,
                    0.5, 1.0), ncol = 2, byrow = T)
traitCor

# Create two traits
SP$addTraitA(
  nQtlPerChr = 100,
  mean = c(0, 0),
  var  = c(1, 1),
  corA = traitCor
)

# Create population
pop = newPop(founderPop)
pop = setPheno(pop, h2 = c(0.5, 0.5))

# ---- Assign selection index values to the miscellaneous slot ----

# Set weights for each trait
weights = c(1, 0.5)

# Naive selection index
selIndex = c(pheno(pop) %*% weights)

# Assign index values to miscellaneous slot
if (AlphaSimRVersion < "1.5.4") {
  pop = setMisc(pop, "selIndex", selIndex)
} else {
  pop@misc$selIndex = selIndex
}

# Function to obtain nice table of miscellaneous slot values
if (AlphaSimRVersion < "1.5.4") {
  getSelIndex <- function(pop) {
    sapply(getMisc(pop, "selIndex"), FUN = function(z) z)
  }
} else {
  getSelIndex <- function(pop) {
    pop@misc$selIndex
  }
}

# Check miscellaneous slot
getSelIndex(pop)

# Order individuals using selection index stored in miscellaneous slot
popOrd = pop[order(getSelIndex(pop), decreasing = TRUE)]
data.frame(id = popOrd@id,
           selIndex = getSelIndex(popOrd))
```
