## Supplementary material for "Plant breeding simulations with AlphaSimR": Scripts: multipleTraits.R

```
# Script name: Simulation of multiple traits in AlphaSimR
#
# Authors: Jon Bancic, Philip Greenspoon, Chris Gaynor, Gregor Gorjanc
#
# Date Created: 2023-01-23
#
# This script first demonstrates how to simulate multiplt traits with
# AlphaSimR:
# i) two correlated traits with the same genetic architecture and
#   fully shared QTL and
# ii) two traits with different genetic architecture and no shared
#   QTLs.
#
# The script also demonstrates how to perform naive or Hazel-Smith
# index selection with pre-assigned weights
#
# WARNING: Simulating traits with no shared QTL had a bug that was
# fixed in AlphaSimR version 1.5.3.

# ---- Clean environment and load packages ----

rm(list = ls())
# install.packages(pkgs = "AlphaSimR")
packageDescription(pkg = "AlphaSimR")$Version
library(package = "AlphaSimR")

# ---- Setup simulation ----

# Create founder haplotypes
founderPop = runMacs(
  nInd     = 100,
  nChr     = 1,
  segSites = 500,
  inbred   = TRUE,
  species  = "WHEAT"
)

# ---- Add two correlated additive traits with fully shared QTL ----

# Set simulation parameters
SP = SimParam$new(founderPop)

# Specify correlation between traits
traitCor = matrix(c(1.0, 0.5,
                    0.5, 1.0), ncol = 2, byrow = TRUE)
traitCor

# Create two traits
SP$addTraitA(
  nQtlPerChr = 100,
  mean = c(0, 0),
  var  = c(1, 1),
  corA = traitCor
)

# Create population
pop = newPop(founderPop)

# Check genetic values and mean
gv(pop)
meanG(pop)
# Check correlation between traits
cov2cor(varG(pop))

# ---- Add two traits with different genetic architectures and no shared QTLs ----

# Bug fixed in AlphaSimR 1.5.3 - check currently installed version
packageDescription(pkg = "AlphaSimR")$Version

# Set simulation parameters
SP = SimParam$new(founderPop)

# Add first quantitative trait with dominance
SP$addTraitAD(
  nQtlPerChr = 100,
  mean   = 0,
  var    = 1,
  meanDD = 0.9,
  varDD  = 0.1
)

# Prevent QTL from first trait being reused
qtlMap = getQtlMap(trait = 1)
SP$restrSegSites(excludeQtl = qtlMap$id)

# Add second additive oligogenic trait
SP$addTraitA(nQtlPerChr = 2,
             mean = 0,
             var  = 1)

# Create population
pop = newPop(founderPop)

# Check that no QTLs are in common
qtlMap2 = getQtlMap(trait = 2)
sum(qtlMap$id %in% qtlMap2$id)

# Check genetic values and mean
gv(pop)
meanG(pop)
# Check correlation between traits
cov2cor(varG(pop))

# ---- Apply selection index ----

# Set weights for each trait
weights = c(1, 0.5)

# Set phenotype
pop = setPheno(pop, h2 = c(0.5, 0.5))

#### Option 1: Apply naive index ####

# Naive selection index
naiveIndex = pheno(pop) %*% weights

# Select 10 best individuals using index function
pop2 = selectInd(pop, nInd = 10,
                 trait = selIndex, # index function
                 b = weights)      # provide weights
#Alternatively, manually select 10 best individuals using index values
pop2 = pop[order(naiveIndex, decreasing = TRUE)][1:10]

#### Option 2: Apply Smith-Hazel index ####

# Calculate Smith-Hazel weights with known variances
b = smithHazel(weights, varG = varG(pop), varP = varP(pop))

# Select 10 best individuals using index function
pop3 = selectInd(pop, nInd = 10,
                 trait = selIndex, # index function
                 b = b)            # provide weights
# Alternatively, manually select 10 best individuals using index values
hazelIndex = selIndex(Y = pop@pheno, b = b)
pop3 = pop[order(hazelIndex, decreasing = TRUE)][1:10]

# Check correlation between naive and Smith-Hazel index
cor(naiveIndex, hazelIndex)
```
