## Supplementary material for "Plant breeding simulations with AlphaSimR": Scripts: setHeritability.R

```
# Script name: Setting heritability in AlphaSimR
#
# Authors: Jon Bancic, Philip Greenspoon, Chris Gaynor, Gregor Gorjanc
#
# Date Created: 2023-01-23
#
# This script demonstrates three ways for setting error variance
# of a phenotype. This is shown using a trait with additive
# and dominance effects.

# ---- Clean environment and load packages ----

rm(list = ls())
# install.packages(pkgs = "AlphaSimR")
library(package = "AlphaSimR")

# ---- Setup simulation ----

# Create founder haplotypes
founderPop = quickHaplo(nInd = 1000,
                        nChr = 1,
                        segSites = 1000)

# Set simulation parameters
SP = SimParam$new(founderPop)

# Add an additve + dominance trait
SP$addTraitAD(
  nQtlPerChr = 1000,
  mean   = 0,
  var    = 1,
  meanDD = 0.92,
  varDD  = 0.2
)

# Create population
pop = newPop(founderPop)

# Obtain true additive and total genetic variance from SP object
# These relate to the founder population and will change with selection
varA = SP$varA[1]
varG = SP$varG[1]

# NOTES:
# Users should choose only one of three options to set phenotype.
# Option 1 and 2 use variances that relate to the founder population
# We assume that heritability of the trait is 0.5.

# ---- Option 1: By assigning the narrow sense heritability ----

pop = setPheno(pop, h2 = 0.5)
# Check heritability
cor(pop@pheno, pop@gv)^2

# Equivalent to manual setting of phenotype as
varE  = varA / 0.5 - varG  # obtain error variance given h2 and varA
error = rnorm(pop@nInd, sd = sqrt(varE))
pheno = pop@gv + error
# Check heritability
cor(pheno, pop@gv)^2

# ---- Option 2: By assigning the broad-sense heritability ----

pop = setPheno(pop, H2 = 0.5)
# Check heritability
cor(pop@pheno, pop@gv)^2

# Equivalent to manual setting of phenotype as
varE  = varG / 0.5 - varG  # obtain error variance given h2 and varG
error = rnorm(pop@nInd, sd = sqrt(varE))
pheno = pop@gv + error
# Check heritability
cor(pheno, pop@gv)^2

# ---- Option 3: By assigning error variance and number of replications ----

pop = setPheno(pop, varE = 1, reps = 1)
# Check heritability
cor(pop@pheno, pop@gv)^2

# Increasing the number of replications will result in higher h2
pop = setPheno(pop, varE = 1, reps = 10)
# Check heritability
cor(pop@pheno, pop@gv)^2

# Equivalent to manual setting of phenotype as
reps  = 10
varE  = 1
error = rnorm(pop@nInd, sd = sqrt(varE))
pheno = pop@gv + error / sqrt(reps)
# Check heritability
cor(pheno, pop@gv)^2
```
