## Supplementary material for "Plant breeding simulations with AlphaSimR": Scripts: simulateGWAS.R

```
# Script name: Performing association study with simulated population
#
# Authors: Jon Bancic, Philip Greenspoon, Chris Gaynor, Gregor Gorjanc
#
# Date Created: 2023-01-23
#
# This script demonstrates how to simulate a population for a GWAS 
# study. The simulation starts of by creating a single homogeneous 
# population which then undergoes a few rounds of selection 
# to induce stratification.

# ---- Clean environment and load packages ----

rm(list = ls())
# install.packages(pkgs = c("AlphaSimR", "ade4", "ggplot2", "rrBLUP", "qqman")
library(package = "AlphaSimR")
library(package = "ade4")
library(package = "ggplot2")
library(package = "rrBLUP")
library(package = "qqman")

# ---- Setup simulation ----

# Generate founder haplotypes
founderPop = runMacs(
  nInd     = 1000,
  nChr     = 10,
  segSites = 300,
  inbred   = FALSE,
  # split    = 100, # alternative way to split population
  species = "GENERIC"
)

# Set simulation parameters
SP = SimParam$new(founderPop)

# Force QTLs and SNPs to overlap
SP$restrSegSites(overlap = T)

# Create additive trait with 5 QTLs per chromosome
SP$addTraitAG(nQtlPerChr = 5,
              mean = 0,
              var  = 1)

# Add SNP-chip with 300 SNPs per chromosome
SP$addSnpChip(nSnpPerChr = 300)

# Check that QTLs and SNPs overlap
sum(getQtlMap(trait = 1)$id %in% getSnpMap(1)$id)

# ---- Create two subpopulations ----

pop  = newPop(founderPop)
popA = pop[1:500]
popB = pop[501:1000]
# Few rounds of crossing and selection
# Alternatively, set split in runMaCS call and skip this
for (i in 1:3) {
  popA = selectCross(popA, nInd = 50, nCrosses = 100, nProgeny = 5, use = "gv")
  popB = selectCross(popB, nInd = 100, nCrosses = 500, use = "gv")
}
pop = c(popA, popB)
pop = setPheno(pop, h2 = 0.4)

# ---- Visualise two sub-populations with PCA ----

geno = pullQtlGeno(pop)
PCA  = dudi.pca(df = geno, center = TRUE, scale = FALSE, scannf = FALSE, nf = 5)
(VAF = 100 * PCA$eig[1:5] / sum(PCA$eig)) # variance explained
df.PCA = data.frame(
  "Pop" = c(rep("Pop1", popA@nInd), rep("Pop2", popB@nInd)),
  "PC1" = PCA$l1$RS1,
  "PC2" = PCA$l1$RS2
)

# Plot
ggplot(df.PCA, aes(x = PC1, y = PC2)) +
  geom_point(aes(colour = factor(Pop))) +
  ggtitle("Population structure") +
  xlab(paste("Pcomp1: ", round(VAF[1], 2), "%", sep = "")) +
  ylab(paste("Pcomp2: ", round(VAF[2], 2), "%", sep = ""))

# ---- Run GWAS ----

# Prepare data
pheno  = data.frame(id     = 1:pop@nInd,
                    pheno  = pop@pheno,
                    subPop = factor(c(
                      rep("Pop1", popA@nInd),
                      rep("Pop2", popB@nInd)
                    )))
geno   = pullSnpGeno(pop)
geno   = data.frame(
  snp = colnames(geno),
  chr = rep(1:10, each = 300),
  pos = rep(1:300, 10),
  t(geno - 1)
)
colnames(geno)[-c(1:3)] = 1:pop@nInd

# Run three GWAS models
model1 = GWAS(pheno[, -3], geno, plot = F)
model2 = GWAS(pheno[, -3], geno, n.PC = 3, plot = F)
model3 = GWAS(pheno,       geno, fixed = "subPop", plot = F)

# Obtain p values
model1$Trait1 = 10 ^ (-model1$Trait1)
model2$Trait1 = 10 ^ (-model2$Trait1)
model3$Trait1 = 10 ^ (-model3$Trait1)

# Get true positions of QTLs
qtl = as.vector(getQtlMap(trait = 1)$id)

# Check Manhattan plot
par(mfrow = c(3, 1))
manhattan(model1, chr = "chr", bp = "pos", p = "Trait1", snp = "snp", highlight = qtl,
  main = "Marker")
manhattan(model2, chr = "chr", bp = "pos", p = "Trait1", snp = "snp", highlight = qtl,
  main = "Marker + principal components")
manhattan(model3, chr = "chr", bp = "pos", p = "Trait1", snp = "snp", highlight = qtl,
  main = "Marker + subpopulation factor")

# Check QQ-plot
par(mfrow = c(3, 1))
qq(model1$Trait1, main = "Marker")
qq(model2$Trait1, main = "Marker + principal components")
qq(model3$Trait1, main = "Marker + subpopulation factor")
```
