## Supplementary material for "Plant breeding simulations with AlphaSimR": Scripts: specifyDemography.R

```
# Script name: Specifying species demography in AlphaSimR
#
# Authors: Jon Bancic, Philip Greenspoon, Chris Gaynor, Gregor Gorjanc
#
# Date Created: 2023-01-23
#
# This script demonstrates how to specify your own demographic history
# when using MaCS simulator of founders' genomes with AlphaSimR.
# Parameters are chosen hypothetically and would normally be obtained
# from literature.

# ---- Clean environment and load packages ----

rm(list = ls())
# install.packages(pkgs = "AlphaSimR")
library(package = "AlphaSimR")

# Load in function to calculate Fst statistic

# ---- Example 1: Specify simple demographic history ----

manualCommand = runMacs2(
  nInd     = 100,
  nChr     = 10,
  segSites = 1000,
  Ne       = 100,
  bp       = 1e+08,
  genLen   = 1,
  mutRate  = 3e-08,
  histNe   = c(500, 1500, 6000,  12e3, 1e5),
  histGen  = c(100, 1000, 10000, 1e5,  1e6),
  ploidy   = 2L,
  inbred   = T,
  returnCommand = TRUE
)
manualCommand

# Create founder haplotypes with desired demography
founderPop = runMacs(
  nInd     = 100,
  nChr     = 10,
  segSites = 1000,
  inbred   = TRUE,
  manualGenLen  = 1,
  manualCommand = manualCommand
)

# Set simulation parameters
SP = SimParam$new(founderPop)

# Create founder population
pop = newPop(founderPop)

# ---- Example 2: Specify demographic history of wheat as in AlphaSimR ----

manualCommand = runMacs2(
  nInd     = 100,
  nChr     = 10,
  segSites = 1000,
  Ne       = 50,
  bp       = 8e+08,
  genLen   = 1.43,
  mutRate  = 2e-09,
  histNe   = c(50, 100, 200, 300, 400, 500, 600, 700, 800, 900, 1e3, 2e3, 3e3, 4e3, 5e3, 6e3, 7e3, 8e3, 9e3, 10e3, 12e3, 16e3, 20e3, 24e3, 28e3, 32e3),
  histGen  = c( 5,  10,  20,  30,  40,  50,  60,  70,  80,  90, 100, 200, 400, 600, 800, 1e3, 2e3, 4e3, 6e3,  8e3, 10e3, 20e3, 40e3, 60e3, 80e3, 10e4),
  ploidy   = 2L,
  inbred   = TRUE,
  # split    = 100, # optional time of population split
  returnCommand = TRUE
)
manualCommand

# Create founder haplotypes with desired demography
founderPop = runMacs(
  nInd     = 100,
  nChr     = 10,
  segSites = 1000,
  inbred   = TRUE,
  manualGenLen  = 1,
  manualCommand = manualCommand
)

# The above call is equivalent to pre-defined "WHEAT" species call in AlphaSimR
# founderPop = runMacs(nInd     = 100,
#                      nChr     = 10,
#                      segSites = 1000,
#                      Ne       = 50,
#                      ploidy   = 2L,
#                      inbred   = TRUE,
#                      # split  = 100, # optional time of population split
#                      species  = "WHEAT")

# Set simulation parameters
SP = SimParam$new(founderPop)
SP$addTraitA(1000)

# Create founder population
pop = newPop(founderPop)
pop1 = pop[1:50]   # first half as Population 1
pop2 = pop[51:100] # second half as Population 2
calcFst(pop1, pop2) # Fst statistic ~ 0 since no split is specified

# ---- Example 3: Modify a MaCS call ----

# Simple MaCS call
manualCommand = runMacs2(
  nInd     = 100,
  nChr     = 10,
  segSites = 1000,
  Ne       = 100,
  bp       = 1e+08,
  genLen   = 1,
  mutRate  = 3e-08,
  histNe   = c(500, 1500,  6000, 12e3, 1e5),
  histGen  = c(100, 1000, 10000,  1e5, 1e6),
  ploidy   = 2L,
  inbred   = TRUE,
  returnCommand = TRUE
)
manualCommand

# This function creates a call for population split in which each
# population has a different effective population size
#' @param Ne effective population size
#' @param histNe effective population size at the time of split
#' @param split historic population split in terms of generations ago
calcMacs <- function(Ne, histNe, split) {
  if (length(histNe) != 2) {
    "Works only for two subpopulatons"
  }
  return = paste0(" -ej ", split / (4 * Ne),
                  " -n 1 ", c(histNe / Ne)[1],
                  " -n 2 ", c(histNe / Ne)[2])
  return(return)
}
newCommand = calcMacs(Ne = 100,
                      histNe = c(100, 50),
                      split = 50)
newCommand

# Merge commands
paste0(manualCommand, newCommand)
```
