## Supplementary material for "Plant breeding simulations with AlphaSimR": Scripts: speedBreeding.R

```
# Script name: Speed breeding
#
# Authors: Jon Bancic, Philip Greenspoon, Chris Gaynor, Gregor Gorjanc
#
# Date Created: 2023-01-23
#
# This script demonstrates speed breeding of plants in a glasshouse
# using AlphaSimR.

# ---- Clean environment and load packages ----

rm(list = ls())
# install.packages(pkgs = "AlphaSimR")
library(package = "AlphaSimR")

# ---- Setup simulation ----

# Generate founder haplotypes
founderPop = runMacs(
  nInd     = 100,
  segSites = 1000,
  inbred   = TRUE,
  species  = "WHEAT"
)

# Set simulation parameters
SP = SimParam$new(founderPop)

# Create additive trait with 10 QTLs
SP$addTraitAG(nQtlPerChr = 10,
              mean = 0,
              var  = 1)

# Create population
pop = newPop(founderPop)

# ---- Example 1: 20 years of breeding without speed breeding ----

pop2 = pop
mean = vector()
mean[1] = meanG(pop2)
# Each year make 100 crosses, grow individuals in field trials,
# and advance 50
for (year in 1:10) {
  pop2 = setPheno(pop2, h2 = 0.5)
  pop2 = selectCross(pop2, nInd = 50, nCrosses = 100)
  mean[year + 1] = meanG(pop2)
}

# ---- Example 2: 10 years of breeding with 4 cycles of speed breeding ----

pop3  = pop
mean2 = vector()
mean2[1] = meanG(pop3)
for (year in 1:10) {
  for (speedCyc in 1:4) {
    # Grow individuals in greenhouse and advance 50
    # Assume accuracy of selection 0.3
    pop3 = setPheno(pop3, h2 = 0.3)
    pop3 = selectCross(pop3, nInd = 50, nCrosses = 100)
  }
  # Grow individuals in field trials and advance 50
  pop3 = setPheno(pop3, h2 = 0.5)
  pop3 = selectInd(pop3, nInd = 50)
  mean2[year + 1] = meanG(pop3)
}

# ---- Compare two breeding scenarios ----

require(ggplot2)
df <- data.frame(
  Year     = 0:10,
  Scenario = c(rep("Conv", 11), rep("Speed", 11)),
  Pheno    = c(mean, mean2)
)

ggplot(df, aes(x = Year, y = Pheno)) +
  geom_line(aes(color = Scenario), linewidth = 0.8)
```
