## Supplementary material for "Plant breeding simulations with AlphaSimR": Scripts: traitIntrogression.R

```
# Script name: Speed breeding
#
# Authors: Initially developed by Chris Gaynor; exanded/polished for this
# publication by Jon Bancic, Philip Greenspoon, Chris Gaynor, Gregor Gorjanc
#
# Date Created: 2023-01-23
#
# This script models a simple trait introgression scheme using AlphaSimR.
# 
# The scheme models a species with 2 chromosomes having a genetic length
# of 1 Morgan.
# 
# Genetic markers spaced 10 cM apart will be used to track IBD between an
# inbred donor line and an inbred recurrent parent. The recurrent parent
# will have the '1' allele at all loci and the donor parent will have the 
# '0' allele.
# 
# A single locus representing a trait being introgressed will be added
# to the middle of the first chromosome. The donor parent will have the 
# '1' allele at this locus and the recurrent parent will have the '0' allele.
# 
# The simulation will model three rounds of backcrossing followed by selfing 
# to obtain homozygous lines for the trait being introgressed. Only plants 
# containing the trait in the heterozygous state are advanced after
# backcrossing.

# ---- Clean environment and load packages ----

rm(list = ls())
# install.packages(pkgs = "AlphaSimR")
library(package = AlphaSimR)

# ---- Setup simulation ----

# Create a genetic map with 2 chromosomes.
# Each chromosome has a genetic length of 1 with markers
# spaced 10 cM apart (11 markers per chromosome).
genMap = data.frame(
  markerName = paste0("M", 1:(2 * 11)),
  chromosome = rep(1:2, each = 11),
  position = rep(seq(from = 0, to = 1, by = 0.1), times = 2)
)

# Create genotypes for parents
geno = rbind(rep(2, 2 * 11),
             rep(0, 2 * 11))
colnames(geno) = genMap$markerName

# Create pedigree with just IDs
# RP stands for recurrent parent
ped = data.frame(
  id = c("RP", "Donor"),
  mother = c(0, 0),
  father = c(0, 0)
)

# Create initial founder population
founderPop = importInbredGeno(geno = geno,
                              genMap = genMap,
                              ped = ped)

# Add the trait being introgressed
# The RP has genotype 0 (c(0,0) haplotypes)
# The donor has genotype 2 (c(1,1) haplotypes)
founderPop = addSegSite(
  founderPop,
  siteName = "Trait",
  chr = 1,
  mapPos = 0.5,
  haplo = matrix(c(0, 0,
                   1, 1), ncol = 1)
)

# Initialize the simulation parameters
SP = SimParam$new(founderPop)

# Exclude "Trait" as an eligible SNP marker
SP$restrSegSites(excludeSnp = "Trait")

# Add a SNP chip for the remaining sites
SP$addSnpChip(nSnpPerChr = 11)

# ---- Parents and Introgression via backcrossing ----

# Create separate population for the recurrent and donor parents
RP = newPop(founderPop[1])
Donor = newPop(founderPop[2])

# Create an F1 plant, only one needed because parents are fully inbred
F1 = randCross2(RP, Donor, nCrosses = 1)

# Create BC1F1 by backcrossing to the recurrent parent
BC1F1 = randCross2(F1, RP, nCrosses = 1000)

# Select BC1F1 for presence of trait
take = pullMarkerGeno(BC1F1, "Trait") == 1
BC1F1 = BC1F1[take]

# Create BC2F1 generation and select for trait
BC2F1 = randCross2(BC1F1, RP, nCrosses = 1000)
take = pullMarkerGeno(BC2F1, "Trait") == 1
BC2F1 = BC2F1[take]

# Create BC3F1 generation and select for trait
BC3F1 = randCross2(BC2F1, RP, nCrosses = 1000)
take = pullMarkerGeno(BC3F1, "Trait") == 1
BC3F1 = BC3F1[take]

# Create BC3F2 generation with selfing
BC3F2 = self(BC3F1)

# Select progeny that are homozygous for the trait
take = pullMarkerGeno(BC3F2, "Trait") == 2
BC3F2 = BC3F2[take]

# ---- IBD analysis ----

# Pull SNP genotypes and divide by 2 to show IBD
IBD = pullSnpGeno(BC3F2) / 2

# Show IBD distribution as percentage by individuals
IBD_ind = rowMeans(IBD) * 100
hist(IBD_ind, xlab = "Percent IBD", main = "Distribution of IBD in BC3F2")

# Show IBD distribution by marker
# Could be made much prettier using ggplot2
# Note that M6 co-localizes with the trait, so it will
# have zero IBD with the recurrent parent.
boxplot(IBD, main = "Distribution of IBD by marker in BC3F2")
```
