## Supplementary material for "Plant breeding simulations with AlphaSimR": Scripts: LineBreeding.html

Line breeding program


### Line breeding program

#### 2023-12-07

#### Introduction

In this vignette we present AlphaSimR code for a line breeding
program using DH technology, as would be used in wheat breeding.
Sections of this vignette mirror subsections 3.2 – 3.5 in the Results
section of the main text, and we suggest the reader refer to it while
reading these sections of the main text. For simplicity, in this
vignette we only display a single year of the burn-in for a single
replicate. For code of the full multi-year and multi-replicate line
breeding program, refer to the AlphaSimR code found in the supplementary
material and on GitHub
`01_LineBreeding/01_PhenotypicSelection/04_DoubledHaploid`
(start with `00RUNME.R`. Also, the supplementary material and
GitHub contain simulation code for numerous other breeding programs.

#### Preliminaries

First we clear our workspace, install required packages if they are
not yet installed, and load required packages.

```
rm(list = ls())
# install.packages(pkgs = "AlphaSimR")
library(package = "AlphaSimR")
```

```
## Loading required package: R6
```

#### Specifying global parameters

We define global parameters that are used throughout a simulation.
Such parameter values can be chosen based on estimates from prior
analyses of real data or long-term trends, trial and error, or practical
experience. For definitions of the parameters see Table 3 in the main
text. Note that in the script files, this step is included with
`source(file = "GlobalParameters.R")`, but here, for
illustrative purposes, we include all code into a single continuous
script. We also define a scenario name so we can track multiple possible
scenarios.

```
# ---- Number of simulation replications and breeding cycles ----
nReps   = 1
nBurnin = 1
nFuture = 0
nCycles = nBurnin + nFuture

# ---- Genome simulation ----
nQtl = 1000
nSnp = 0

# ---- Initial parents mean and variance ----
initMeanG  = 1
initVarG   = 1
initVarEnv = 1e-6 
initVarGE  = 2
varE       = 4

# ---- Breeding program details ----
nParents = 50
nCrosses = 100
nDH      = 100
famMax   = 10
nPYT     = 500
nAYT     = 50
nEYT     = 10

# Effective replication of yield trials
repHDRW  = 4/9
repPYT   = 1
repAYT   = 4
repEYT   = 8

scenarioName = "LinePheno_DH"
```

We create a list to store results from replications.

```
results = list()
```

First we always run just a single replicate of our simulation. Once
we have completed all the steps of the simulation and the simulation
runs as expected, we can run multiple replicates. Note that in the
script files we run through a `for` loop of the multiple
replicates, but for illustrative purposes here, we only have one
replicate.

```
REP = 1
```

We create a data frame to track key parameters.

```
output = data.frame(year     = 1:nCycles,
                    rep      = rep(REP, nCycles),
                    scenario = rep(scenarioName, nCycles),
                    meanG    = numeric(nCycles),
                    varG     = numeric(nCycles),
                    accSel   = numeric(nCycles))
```

#### Simulating genomes and founders

We simulate founder genomes and create initial parents. AlphaSimR
embeds MaCS software to generate founder genomes through a
backward-in-time (coalescent) simulation. Here we use a pre-defined
demographic history for wheat. Note that in the script files this step
is included with `source(file = "CreateParents.R")`.

First we create a founder population, by generating initial
haplotypes using `runMacs()`; and make the global simulation
parameter (`SP`) object.

```
founderPop = runMacs(nInd     = nParents, 
                     segSites = nQtl + nSnp,
                     inbred   = TRUE, 
                     species  = "WHEAT")
SP = SimParam$new(founderPop)
```

We can include a SNP chip if desired (relevant for genomic selection
strategies explained elsewhere).

```
SP$restrSegSites(nQtl, nSnp)
if (nSnp > 0) {
  SP$addSnpChip(nSnp)
}
```

We add a trait, representing yield, and specify the mean, and
variance of QTL effects, as well as the environmental variance and
variance due to genotype-by-environment interactions.

```
SP$addTraitAG(nQtlPerChr = nQtl,
              mean       = initMeanG,
              var        = initVarG,
              varEnv     = initVarEnv,
              varGxE     = initVarGE)
```

We opt to track pedigree information.

```
SP$setTrackPed(TRUE)
```

We create founder parents and set a phenotype, reflecting evaluation
in EYT stage (see the main text).

```
Parents = newPop(founderPop)
Parents = setPheno(Parents, varE = varE, reps = repEYT)
```

Finally we clear the founderPop object to free some computer
memory.

```
rm(founderPop)
```

#### Filling the breeding pipeline

We fill the breeding pipeline with distinct populations for each
stage. The filling process is necessary to capture the overlap of
different breeding cycles in real breeding programs, meaning that at any
given time, a breeding program will have populations in all stages of
the breeding program. We achieve this by running the founder parents
through the crosses and selection steps of our breeding program, saving
a population at a different breeding stage each time.

Our wheat breeding program consists of six breeding stages and hence,
we need to generate six distinct breeding populations through this
filling process. The first round runs through all stages, saving the
population in the final (sixth) stage. The second round runs through the
stages up to the fifth stage, saving the population at that stage. This
process is repeated until all stages have been populated. Despite using
the same founder parents, distinct populations are obtained because each
round involves unique crossings, random process of inheritance, and
selections. We will update parents after the filling process.

Note that in the script files this step is included with
`source(file = "FillPipeline.R")`.

```
# Set initial yield trials with unique individuals
for (cohort in 1:7) {
  cat("  Fill pipeline stage:", cohort, "of 7\n")
  if (cohort < 7) {
    # Stage 1
    F1 = randCross(Parents, nCrosses)
  }
  if (cohort < 6) {
    # Stage 2
    DH = makeDH(F1, nDH)
  }
  if (cohort < 5) {
    # Stage 3
    HDRW = setPheno(DH, varE = varE, reps = repHDRW)
  }
  if (cohort < 4) {
    # Stage 4
    PYT = selectWithinFam(HDRW, famMax)
    PYT = selectInd(PYT, nPYT)
    PYT = setPheno(PYT, varE = varE, reps = repPYT)
  }
  if (cohort < 3) {
    # Stage 5
    AYT = selectInd(PYT, nAYT)
    AYT = setPheno(AYT, varE = varE, reps = repAYT)
  }
  if (cohort < 2) {
    # Stage 6
    EYT = selectInd(AYT, nEYT)
    EYT = setPheno(EYT, varE = varE, reps = repEYT)
  }
  if (cohort < 1) {
    # Stage 7
  }
}
```

```
##   Fill pipeline stage: 1 of 7
##   Fill pipeline stage: 2 of 7
##   Fill pipeline stage: 3 of 7
##   Fill pipeline stage: 4 of 7
##   Fill pipeline stage: 5 of 7
##   Fill pipeline stage: 6 of 7
##   Fill pipeline stage: 7 of 7
```

We will now explain each of the operations performed in filling the
pipeline, which are seen again in the Running the Burn-in phase
section.

In “Stage 1”, `F1 = randCross(Parents, nCrosses)`,
performs `nCrosses` random crosses of the parents to produce
`nCrosses` F1s.

In “Stage 2” `DH = makeDH(F1, nDH)` uses doubled-haploid
technology to make `nDH` double haploids per each F1. By
definition, these DH individuals are homozygous at all loci.

In “Stage 3”
`HDRW = setPheno(DH, varE = varE, reps = repHDRW)`
sets/simulates a phenotype for the DH lines with a certain error
variance and replication, which captures a certain heritability. The
phenotype will be used as the basis for breeder’s selection.

In “Stage 4” `PYT = selectWithinFam(HDRW, famMax)`
performs selection within families such that no more than
`famMax` DH lines per family can be carried forward;
`PYT = selectInd(PYT, nPYT)` performs selection among all
individuals in the Preliminary Yield Trial choosing the best
`nPYT`; and
`PYT = setPheno(PYT, varE = varE, reps = repPYT)`
sets/simulates a phenotype of the remaining individuals.

In “Stage 5” `AYT = selectInd(PYT, nAYT)` performs
selection in the Preliminary Yield Trial; and
`AYT = setPheno(AYT, varE = varE, reps = repAYT)`
sets/simulates a phenotype for the remaining individuals.

In “Stage 6” `EYT = selectInd(AYT, nEYT)` performs
selection in the Advanced Yield Trial; and
`EYT = setPheno(EYT, varE = varE, reps = repEYT)` sets a
phenotype for the remaining individuals in the first year of Elite Yield
Trials.

#### Running the burn-in phase

Now we will go through a single year of the breeding program’s
burn-in phase. In a real simulation, we would run multiple years of
burn-in followed by multiple years of future breeding.

```
year = 1
```

##### Update Parents

First we update our parents by replacing the 10 oldest parents with
10 new parents from EYT stage. Note that in the script files this step
is included with `source(file = "UpdateParents.R")`.

```
Parents = c(Parents[11:nParents], EYT)
```

##### Advance Year

Then we advance a year, going through all the crossing and selection
steps. In one year, each stage of the breeding program is updated,
reflecting the parallel breeding pipelines that are progressing. Note
that in the script files this step is included with
`source(file = "AdvanceYear.R")`.

```
# Stage 7
# Release variety

# Stage 6
EYT = selectInd(AYT, nEYT)
EYT = setPheno(EYT, varE = varE, reps = repEYT)

# Stage 5
AYT = selectInd(PYT, nAYT)
AYT = setPheno(AYT, varE = varE, reps = repAYT)

# Stage 4
PYT = selectWithinFam(HDRW, famMax)
PYT = selectInd(PYT, nPYT)
PYT = setPheno(PYT, varE = varE, reps = repPYT)

# Stage 3
HDRW = setPheno(DH, varE = varE, reps = repHDRW)

# Stage 2
DH = makeDH(F1, nDH)

# Stage 1
F1 = randCross(Parents, nCrosses)
```

##### Backward vs forward pipeline pass

An important note! We work backward through the pipeline to avoid
copying a population before performing operations on it. This is an
important point because failing to do so is a common source of errors in
simulating breeding programs with AlphaSimR. To explain further,
consider what would happen if we went forward through the pipeline
instead.

```
# Stage 1
F1 = randCross(Parents, nCrosses)

# Stage 2
DH = makeDH(F1, nDH)
```

While the above might look reasonable and is syntactically valid R
and AlphaSimR code, note that by making a new F1, we have erased the F1
from the previous year, which we require to make the DH for this year.
The above code effectively fast-tracks the crossing of parents to create
F1 and creation of DH from the F1 in one time step. If we continue in
this way, we would fast-track cohorts through all the stage of the
breeding pipeline in one time step, which is biologically and
logistically impossible.

A correct way to work forward through the pipeline would be to
duplicate populations at each stage, as in the following, but this is
inefficient both in terms of computer memory and code presentation.

```
# Stage 1
F1_saved_for_DH = F1
F1 = randCross(Parents, nCrosses)

# Stage 2
DH_saved_for_HDRW = DH
DH = makeDH(F1_saved_for_DH, nDH)
```

We avoid this needles duplication of populations (with a forward pass
through the pipeline) by running backward through the pipeline, as shown
above in previous code chunks.

##### Saving simulation summaries

Finally, for each year we save results in the `output`
data frame. Note that results could here invilve complete simulated
populations, associated data, or just data summaries as shown here.
Here, we save mean of genetic values and variance of genetic values for
individuals in the DH stage, while for accuracy of selection we
calculate it at the HDRW stage where first phenotypes are available.
Note that heritability of these phenotypes is very low, representing
breeder’s visual selection.

```
output$meanG[year]  = meanG(DH)
output$varG[year]   = varG(DH)
output$accSel[year] = cor(HDRW@pheno, HDRW@gv)
```

#### Wrap-up

The above completed simulation of a single year of the burn-in of a
wheat breeding program for a single replicate. Explore additional
material available in R scripts with multi-year, multi-rep, and
multi-scenario examples.
