## Supplementary material for "Plant breeding simulations with AlphaSimR": Scripts: README.md

# Plant breeding simulations with AlphaSimR

## Introduction

This repository contains R scripts that demonstrate simulations of plant breeding programs and techniques using AlphaSimR https://cran.r-project.org/package=AlphaSimR. These scripts and the process of simulating plant breeding programs are described in:

Bancic et al. (2023) bioRxiv TODO

    @article{BancicEtAl2023,
      title = {Plant breeding simulations with AlphaSimR},
      author = {Bančič, Jon and Greenspoon, Philip and Gaynor, R Chris and Gorjanc, Gregor},
      journal = {bioRxiv},
      year = {2023},
      doi = {TODO}
    }

If you have no prior experience with AlphaSimR, we suggest you complete the free online course "Breeding Programme Modelling with AlphaSimR" at https://www.edx.org/course/breeding-programme-modelling-with-alphasimr. The course is continually open, so that you can enrol when it's convenient for you.

## Repository contents

  * `README.md` is this file.

  * `LineBreeding.Rmd` (and its output `LineBreeding.html`) - Introductory vignette describing the logic of provided R scripts. 

  * `jbancic_alphasimr_plants.Rproj` is the RStudio/Posit project file (to set the working directory etc.).

  * `01_LineBreeding` folder contains scripts for simulating a line breeding program - showing a wheat example that can be adapted to other selfing species. The scripts show several phenotypic selection strategies (mass selection, single seed descent, pedigree selection, and doubled haploids) and genomic selection strategies (conventional and two-part).

  * `02_ClonalBreeding` folder contains scripts for simulating a clonal breeding program - showing a tea example that can be adapted to other clonal species. The scripts show phenotypic, pedigree, or genomic selection strategies.

  * `03_HybridBreeding` folder contains scripts for simulating a hybrid breeding program - showing a maize example that can be adapted to other outcrossing species. The scripts show a phenotypic selection strategy and two genomic selection strategies (conventional and two-part).

  * `04_Techiques` folder contains scripts for simulating plant breeding programs using various breeding and genetic techniques. The scripts show:
  
    * Trait introgression (`traitIntrogression.R`)
  
    * Speed breeding (`speedBreeding.R`)
  
    * Genome editing (`genomeEditing.R`)
    
    * Multi-trait selection with selection index (`multipleTraits.R`)
    
    * Miscellaneous slot to store user-defined information (`miscellaneousSlot.R`)
    
    * Mating plans (`matingPlans.R`)
    
    * GWAS (`simulateGWAS.R`)
    
    * Import external haplotypes (`importExternalHaplo.R`)
    
    * Specifying demography for simulating founder genomes (`specifyDemography.R`)
    
    * Genomic prediction & selection models (`genomicModels.R`)
    
    * Setting heritability (`setHeritability.R`)
    
    * Functions used across scripts (`functions.R`)

## How to work with the provided R scripts

  1) Read the manuscript and browse the introductory vignette `LineBreeding.Rmd` (and its output `LineBreeding.html`).

  1) Download this whole repository
  
  2) In RStudio/Posit, open project file `jbancic_alphasimr_plants.Rproj`
  
  3) For each of the three types of plant breeding programs, select a folder
     and work through the `00RUNME.R` file to run simulations and then work through
     the `ANALYZERESULTS.R` file to analyze the simulations' results.
