## Supplementary figures and images for "Plant breeding simulations with AlphaSimR"

### 06_Figure.png

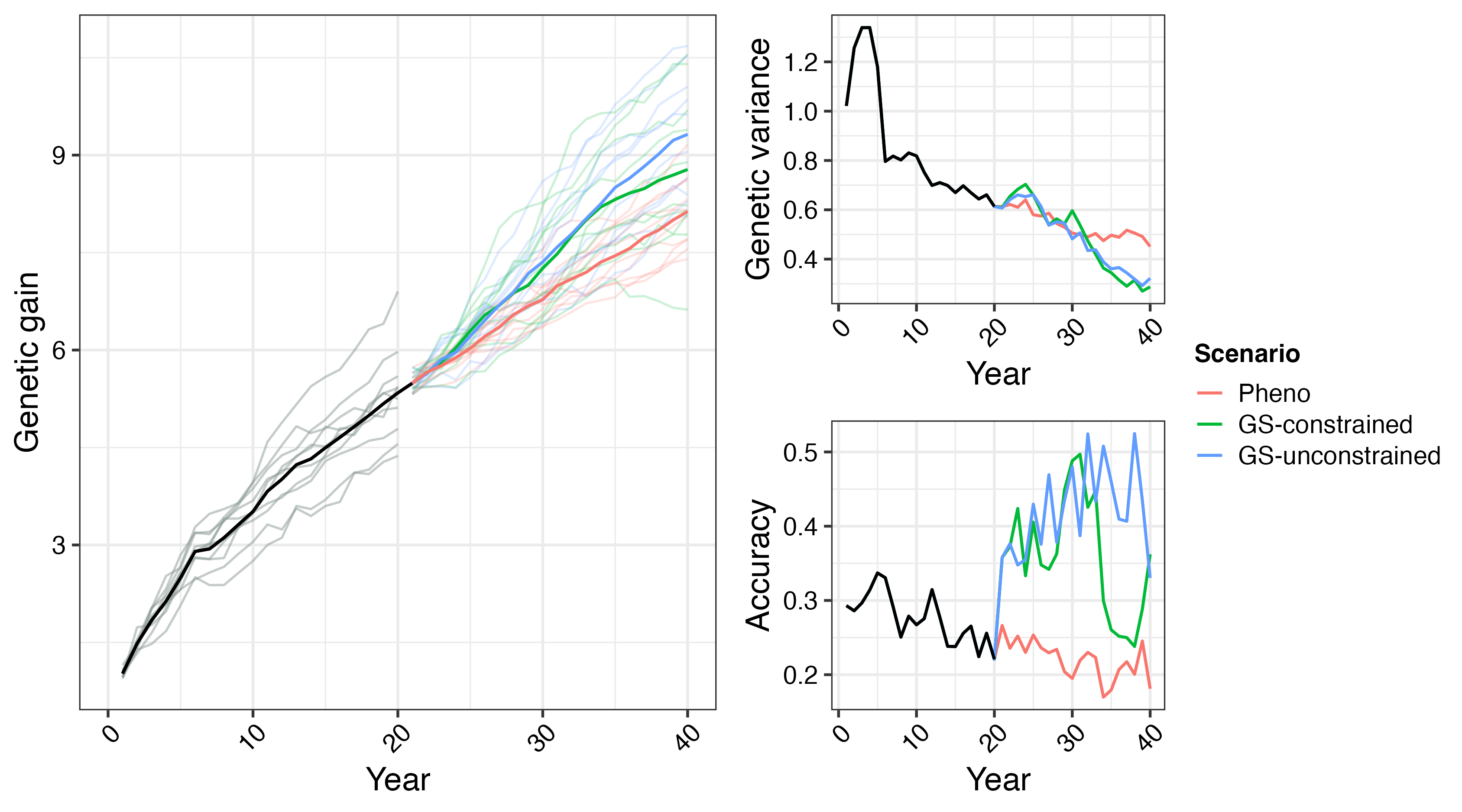

### Results.png

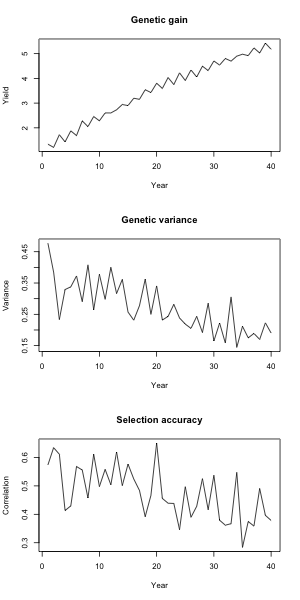

### Results.png

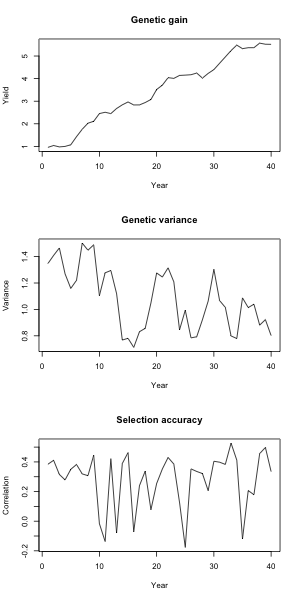

### Results.png

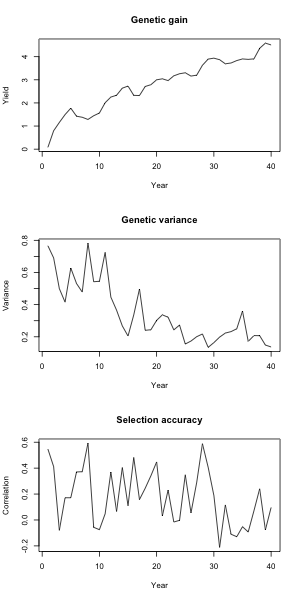

### Results.png

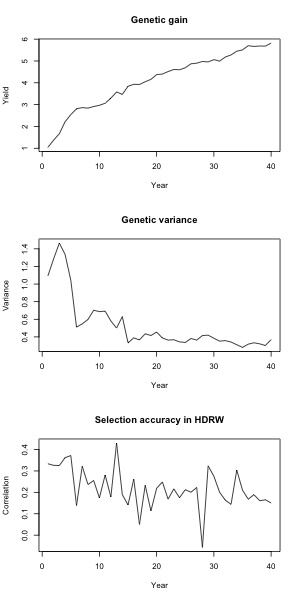

### Results.png

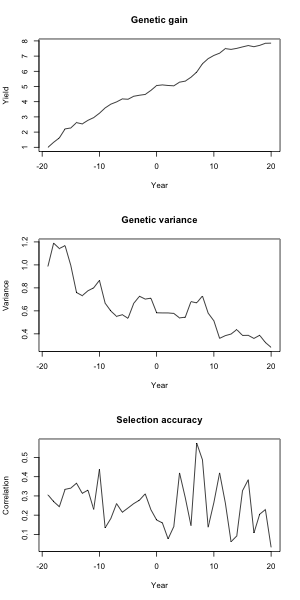

### Results.png

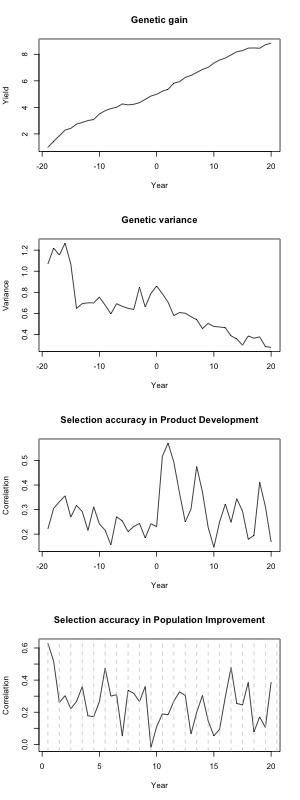

### Results.png

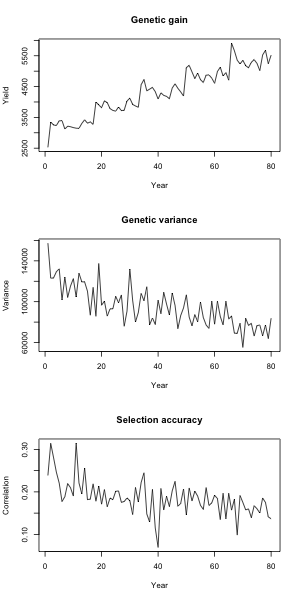

### Results.png

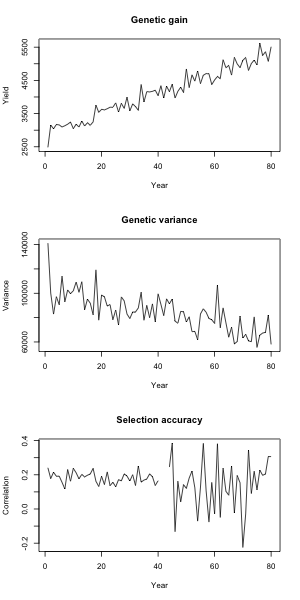

### Results.png

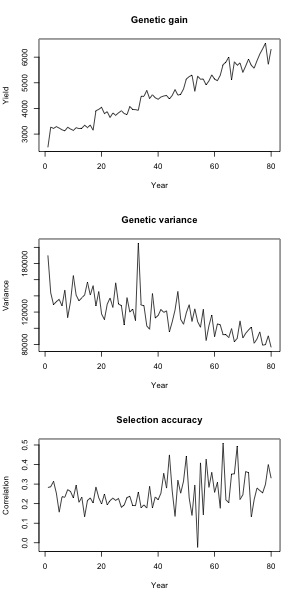

### Results.png

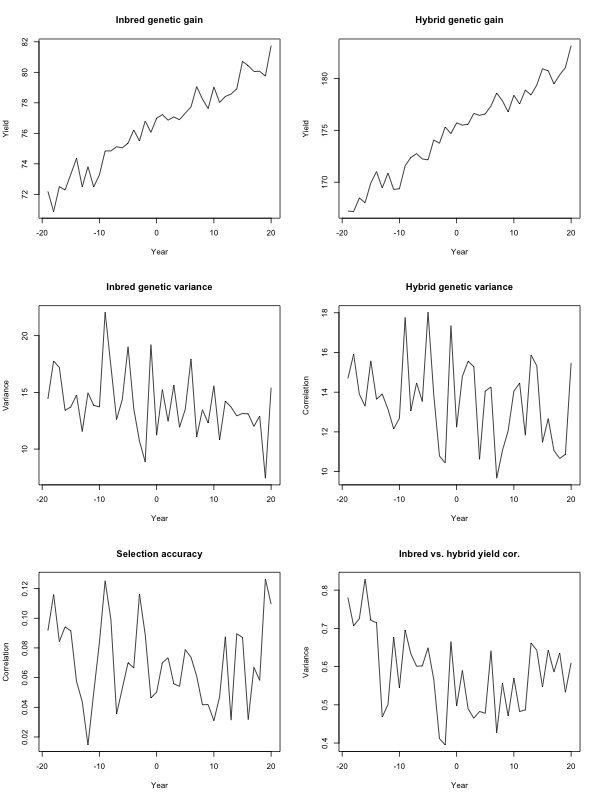

### Results.png

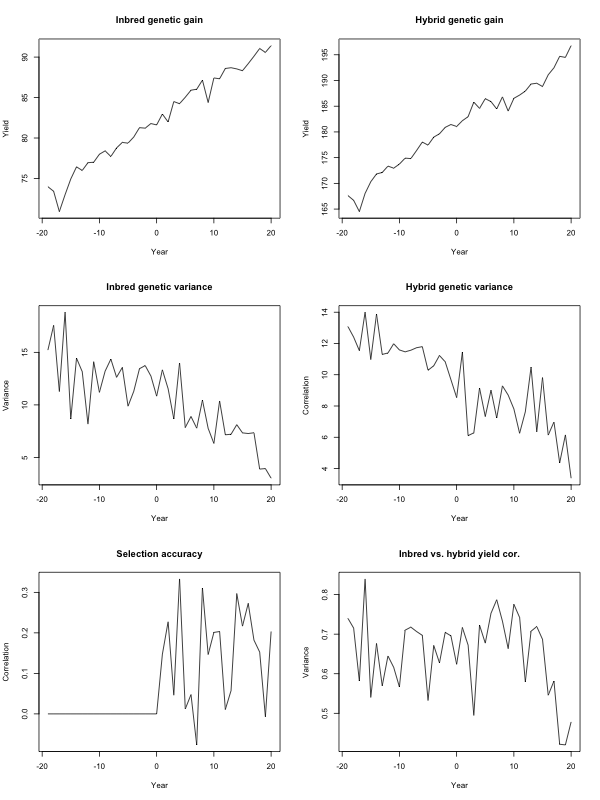

### Results.png

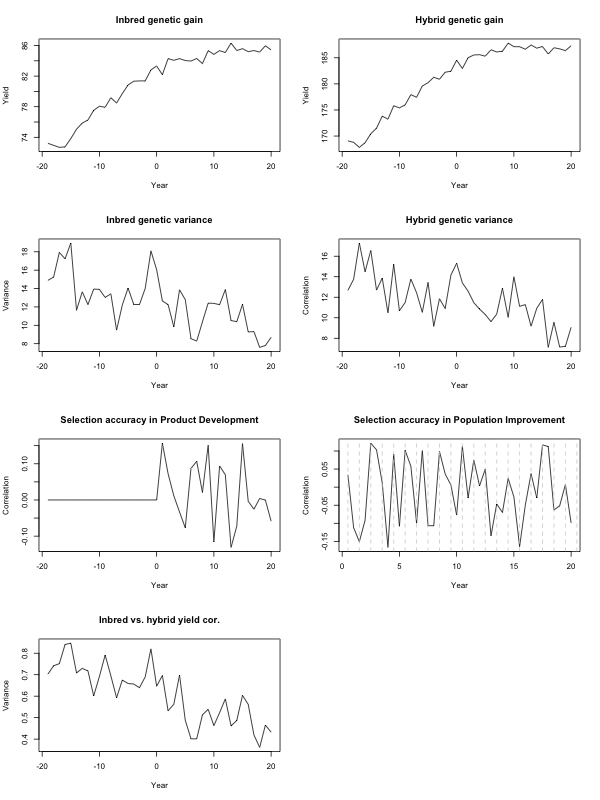
